# Fall of giants: European diversity resistance to waterscape degradation and its potential continental drivers

**DOI:** 10.1101/2024.01.11.574179

**Authors:** David Cunillera-Montcusí, Ana-Inés Borthagaray, Jordi Bou, Matías Arim

**Affiliations:** Institute of Aquatic Ecology. HUN-REN Centre for Ecological Research, Budapest, Hungary; Departamento de Ecología y Gestión Ambiental, Centro Universitario Regional del Este (CURE), Universidad de la República, Tacuarembó s/n, Maldonado, Uruguay; FEHM-Lab. Section of Ecology, Department of Evolutionary Biology, Ecology and Environmental Sciences. University of Barcelona, Diagonal, 643 (08028 Barcelona, Catalonia/Spain); GRECO, Institute of Aquatic Ecology, University of Girona, Girona, Spain; LAGP-Flora and Vegetation, Institute of the Environment, University of Girona, Girona, Catalonia, Spain

**Keywords:** Diversity decay, landscape fragmentation, habitat loss, landscape degradation, landscape conservation, aquatic habitats

## Abstract

Aquatic landscapes, or waterscapes, face severe threats from human activities propelling their deterioration. Waterscape degradation represents a main driver of the current diversity crisis, although its long-term consequences are difficult to quantify. In addition, the understanding of the potential effects of waterscape degradation on biodiversity at large spatial scales, such as biomes and continents, remains limited. In this work, we explore the potential trends in diversity decay in response to waterscape degradation across Europe to provide a first answer to these main threads. We reconstructed the European waterscape based on available satellite data and simulated diversity patterns using a coalescent metacommunity model run for several European ecoregions and considering nine dispersal abilities. Subsequently, we generated a gradient of waterscape degradation by systematically removing a percentage of available habitat and recalculating diversity metrics. For each ecoregion, dispersal ability, and degradation level we obtained a theoretical gamma diversity value. We synthesized the captured diversity decay trends in two parameters: the proportional decay rate and the collapsing rate, which respectively inform about the speed of diversity loss and its acceleration as waterscape degradation progresses. Finally, we mapped these parameters across Europe and related them with ecoregion structural descriptors (i.e. geographic location, water cover, connectivity, and size). Through this exercise, we could identify ecoregions’ sensitivities and their continental variation based on their diversity decay parameters. Connectivity and water cover emerged as primary descriptors of diversity decay, with more heterogeneous ecoregions generally exhibiting greater resistance to waterscape degradation. Our study provides a first order approximation to an urgently needed information: the continental-scale consequences of waterscape degradation for biodiversity. This data has the potential to improve conservation practices and facilitate the integration of innovative approaches in management, thereby enhancing our understanding of the consequences posed by one of the principal threats to freshwater diversity.

## Introduction

Landscape structure defines the main canvas where populations grow and decay, communities assemble and ecosystems function (Gounand et al., 2018; Turner & Gardner, 2015). Consequently, landscape degradation is one of the most pervasive problematics affecting most of the world natural systems (Fuller et al., 2015; Haddad et al., 2015; Wade et al., 2003). Because of this, understanding the potential responses of diversity structure and functioning to landscape degradation constitutes an imperious need for ecology and conservation biology against human-mediated changes (IPCC, 2022; Maasri et al., 2022; Reid et al., 2019). However, landscape can be understood in many ways, especially when considering the point of view of an organism and its dispersal ability (Bilton et al., 2001; Cañedo-Argüelles et al., 2015; Phillipsen et al., 2015). Dispersal has largely been acknowledged as one of the main metacommunity driving forces (Leibold & Chase, 2018; Thompson et al., 2020), and its roots as a key element of assembly and diversity go deep into some of the main seminal works (Hubbell, 2001; MacArthur & Wilson, 1963; Mouquet et al., 2005). However, its connection to the landscape has remained difficult to quantify and analyse (Jacobson & Peres-Neto, 2010; Leibold et al., 2022; Turner & Gardner, 2015). Part of this complexity is explained because dispersal depends on both the ability of a species to move (Cunillera-Montcusí et al., 2021; Heino, 2013; Sarremejane et al., 2020) and the spatial distribution of habitat patches through the landscape (Cañedo-Argüelles et al., 2015; Juračka et al., 2019; Ryberg & Fitzgerald, 2016). Indeed, how habitat distribution favours the exchange of organisms between them can greatly determine diversity patterns at a regional scale, for example by fostering source-sink dynamics or arrival of invasive species (Borthagaray, Pinelli, et al., 2015; Brooks et al., 2018; Cunillera-Montcusí et al., 2023; Manenti et al., 2020). The quantification of this landscape structure is generally done at a determined scale, based on an specific study-case or site spanning a spatially defined region (e.g., Arntzen et al., 2017; Bishop-Taylor et al., 2017; Cañedo-Argüelles et al., 2015; Cunillera-Montcusí et al., 2021; Lai et al., 2022; Seymour et al., 2015). However, the scale that defines a particular landscape structure, and consequently its assembly patterns, can be much bigger and be related to continental-scale descriptors (Carroll et al., 2018; Heino et al., 2015; Santini et al., 2016). Such a biogeographic point of view on the relevance of landscape structure using a network perspective has been done in a descriptive way, for example to detect the main connectivity hotspots or more important dispersal routes (Carroll et al., 2018; Cunillera-Montcusí et al., 2023). These approximations represent major steps in quantifying, at a continental-scale, main dispersal fluxes or source-sink dynamics which are key to improve conservation policies (Chase, Jeliazkov, et al., 2020; Cid et al., 2020; Santini et al., 2016) and acknowledge the changing nature of landscape fostered by human impacts (Fardila et al., 2017; Segan et al., 2016). However, the consideration of habitat loss and fragmentation impacts on a biogeographic level still remains as an avenue to further explore (Pardini et al., 2018).

Freshwater ecosystems are one of the most endangered systems due to global change (Albert et al., 2021; IPCC, 2022; Rounsevell et al., 2018). Human activity has impacted freshwater ecosystems through changes in the natural concentration of nutrients (de Vries et al., 2013; Meerhoff et al., 2022), other chemical ions (Kaushal et al., 2021), the spread of invasive species (Drake et al., 2017; Radinger & García-Berthou, 2020), land-use change and habitat loss (Albert et al., 2021; Chase, Blowes, et al., 2020; Mouton et al., 2022; P. J. Wood et al., 2003), and the complex interactions among these main drivers (Brauns et al., 2022; Jackson et al., 2016). Although these drivers are generally shared among all types of freshwaters, some systems have received less attention than others, with permanent systems being generally more studied than temporary or ephemeral ones (Bagella et al., 2016; Bonada et al., 2020; Calhoun et al., 2017; Datry et al., 2014). This bias is particularly worrying when assessing landscape degradation, as smaller and more temporary systems host an important fraction of freshwater biodiversity (Arntzen et al., 2017; Biggs et al., 2017; Boix et al., 2017; Hill et al., 2021; P. J. Wood et al., 2003). These understudied systems play a key role at the regional scale as they act as stepping-stones that allow dispersal across other freshwater habitats, thus, holding overall meta-system dynamics (Borthagaray, Cunillera-Montcusí, Bou, Biggs, et al., 2023; Cid et al., 2022; Gounand et al., 2018; Wells & Richmond, 1995). Due to this main role in bounding habitats at a large-scale (e.g. ecoregion), research aimed at understanding waterscape dynamics should consider a range of types of freshwaters, from permanent to ephemeral (Biggs et al., 2017; Borthagaray, Cunillera-Montcusí, Bou, Biggs, et al., 2023; Williams et al., 2004).

The degradation of landscapes implies the loss of habitat and its concomitant fragmentation, both recognized as main drivers of the diversity crisis. Consequently, they must be considered together to capture the potential decline in diversity due to landscape degradation (Fagan, 2012; Hanski & Ovaskainen, 2000; Horváth et al., 2019). However, the assessment of landscape degradation has been generally approached at relative small spatial scales (Barbosa & Marquet, 2002; Carrara et al., 2014; Chase, Blowes, et al., 2020; C. Chisholm et al., 2011; Horváth et al., 2019). Some biogeographic quantifications have been done in terrestrial ecosystems (Kuipers et al., 2021) but remain not fully explored for inland waters at such scales, even though these systems are markedly influenced by their spatial structure (Borthagaray, Cunillera-Montcusí, Bou, Biggs, et al., 2023; Brauer & Beheregaray, 2020; Buono et al., 2019; Fuller et al., 2015, p. 20; Tonkin et al., 2018). Species inhabiting these ecosystems depend on the landscape (or waterscape) structure as they rely on the existence of wet corridors to maintain viable populations, communities, and ecosystems. This is true even for species that disperse by flight, for which wet corridors such as river courses are key for their diversity patterns (Bogan & Boersma, 2012; Cañedo-Argüelles et al., 2015; Sarremejane et al., 2017). Therefore, an assessment of the relevance of landscape degradation on freshwater ecosystems has not been fully explored yet, especially considering a biogeographic-scale (Abell et al., 2008) and accounting with different levels of landscape degradation to quantify overall ecoregion’s sensitivity to this disturbance.

In this work, we aim to quantify the impact on diversity generated by landscape degradation, hereafter referred to as diversity decay patterns. To achieve this goal, we used the freshwater distribution of European inland waters (Pekel et al., 2016) and quantified diversity decay patterns based on a coalescent metacommunity model. We simulated proportional losses of freshwater habitats (from 0% loss up to 99%) to evaluate the trend in diversity as waterscape degraded. Two parameters were used to describe this declining trend in diversity, the proportional decay rate, which is the mean reduction in diversity for each fraction of lost waterscape, and the collapsing rate, which estimates the acceleration in the previous rate with each novel fraction of lost waterscape. By means of these two parameters, we intend to quantify the sensitivity of freshwater ecoregions in Europe to landscape degradation considering all freshwaters together or discerning among permanent, temporary, and ephemeral. Finally, we assessed the potential role of geographic location, connectivity, water cover and ecoregion size as determinants of each ecoregion diversity decay pattern. Overall, we aim to incorporate a larger-scale perspective into conservation of freshwater ecosystems, highlighting ecoregional fragility to one of the main global change impacts (Carpenter et al., 1992; IPCC, 2022).

## Methods

This study is organized in several parts that jointly contribute to quantify the impact on diversity generated by landscape degradation across Europe (see Figure 1). The first part refers to the definition of the used databases (Figure 1 Habitat quantification, Dispersal and Regional Pool). Secondly, we introduce the simulation model used and the setup of degradation scenarios (Figure 1 Coalescent model and Scenarios simulation). Thirdly, we explain how results are summarised in two informative parameters capturing the diversity decay patterns (Figure 1 Diversity decay parameters).

**Figure 1:**
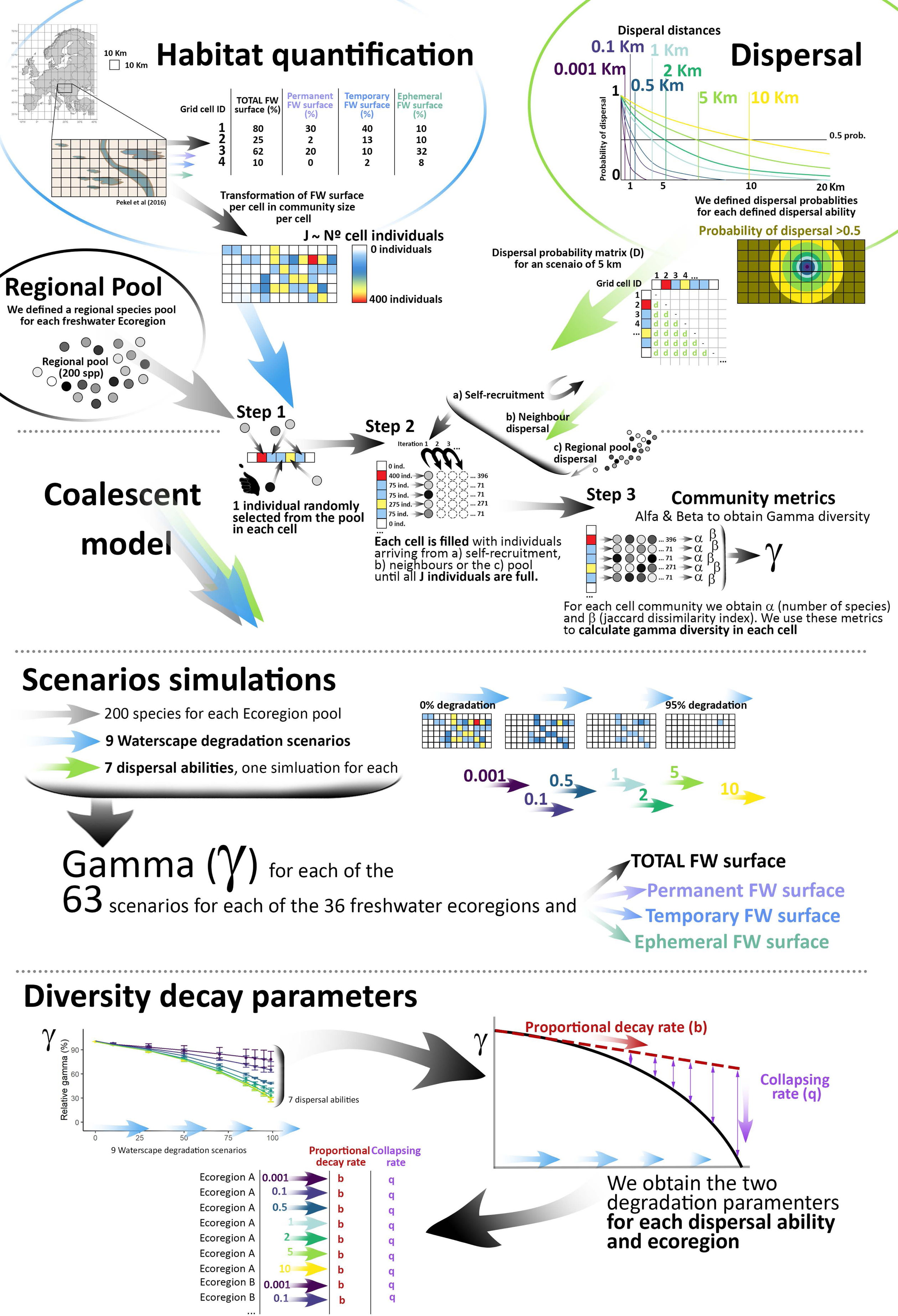
Methodological schema of the different steps followed in this study. Step 1: data obtention and model settings design (Habitat quantification, dispersal abilities, pool). Step 2: model structure and scenario definitions (Coalescent model and Scenarios simulation). Step 3: results processing and calculation of diversity decay parameters: the proportional decay rate (b) and the collapsing rate (q). See methods sections to a detailed description of each step.

### Habitat quantification and ecoregion delimitation

We retrieved the European continent surface freshwater data from the databases linked to Pekel et al. (2016). In this work, the authors quantified surface water based on Landsat satellite images with a resolution of 30-m per pixel. We used the Occurrence dataset, which considers all freshwater surface data from 1984 until 2015. This dataset also accounts for water seasonality, interpreted as the total frequency in which each pixel has remained with water during the study-period (i.e., 0 being a land pixel and 100 a permanent water body that has not been dry in the study period). We treated, depurated, and calculated water surfaces using ArcGIS Pro and the ArcToolbox>Spatial Analyst Tools (ESRI Inc., 2020). In summary, we extracted and used all water bodies included in the considered study area, the European continent. We then built a grid encompassing all this area and having cells of 10×10km. Finally, we merged these two layers and obtained a freshwater surface value for each cell of the grid, which later constituted the community size for model simulations (see Cunillera-Montcusí et al., 2023). We calculated the surface for the 4 considered types of freshwaters: a) total freshwater surface; b) permanent freshwater surface (i.e., pixels presenting water most of the captured time, 90 to 100 values); c) temporary freshwater surface (i.e., pixels presenting water once per year at least, 10 to 89 values), and d) ephemeral freshwaters surface--i.e., pixels presenting water less than once a year, 1 to 9 values).

All the studied area was divided in 36 working units that corresponded to the main freshwater ecoregions present in the European continent (Abell et al., 2008). This division was done to work within ecologically consistent units that were defined by common phylogenetic, biogeographical and/or climatic criteria. The classification criteria as well as the procedure followed to define these ecoregions can be found in Abell et al. (2008). We used ecoregions as working units to acknowledge the fact that waterscape degradation and its consequent diversity loss will have a common impact and respond to the same drivers within a determined biogeographical unit (e.g., waterscape degradation will not have the same consequences nor affect the same species in Western Iberia than in the Northern Baltic Drainages as ecological, evolutionary, or climatic backgrounds of these ecoregions are not the same either). The 36 selected ecoregions and their complete names can be found in Supplementary material 1.

### Dispersal abilities

Within each simulation, we modelled dispersal as a kernel with exponential decay based on the distance between cells (Borthagaray, Cunillera-Montcusí, Bou, Biggs, et al., 2023; Cunillera-Montcusí et al., 2023; Muneepeerakul et al., 2008). We considered a range of dispersal decay patterns to capture different dispersal abilities that could interact with freshwater habitats. The distance at which the probability of dispersal was 0.5 was used as an indicator of the dispersal ability of the species under analysis (Borthagaray, Cunillera-Montcusí, Bou, Tornero, et al., 2023); Supplementary material 2). We defined 7 different dispersal abilities ranging from 0.001km, which would represent organisms with very limited movement capacity (e.g., Incagnone et al. 2015), to 10km, which would represent organisms able to cover large distances across the waterscape (e.g., Figuerola & Green, 2002; Minot et al., 2020) including intermediate dispersal ranges (i.e. 0.1, 0.5, 1, 2, 5km).

### Species pool

For each ecoregion we defined 200 as the total number of species of the regional pool. This number was considered the same for all ecoregions to set a common level of regional diversity as a reference for comparing results. Consequently, we assessed only the decay in diversity linked to waterscape degradation, not including prior-diversity differences per ecoregion even though we acknowledge their existence (Abell et al., 2008).

### Coalescent model

We used a spatially explicit coalescent model (Borthagaray, Cunillera-Montcusí, Bou, Biggs, et al., 2023; Borthagaray, Cunillera-Montcusí, Bou, Tornero, et al., 2023; Cunillera-Montcusí et al., 2021; Hubbell, 2001; Worm & Tittensor, 2018). The model considers each cell of the grid as a community with its size determined by the cell freshwater surface, *AA*_*j*_. Community size, hereafter *J*, was set at a maximum of 400 individuals per cell. The number of individuals in each specific cell based on the area occupied by freshwater was, 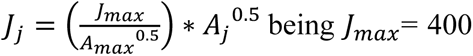 *AA*_max_=100. (Supplementary material 2). Therefore, for each type of freshwater (total, permanent, temporary, and ephemeral) we associated the maximum surface values (i.e. the cell is fully covered by freshwater) to the greatest community size (400 individuals). It should be highlighted that the individuals in the model are not biological individuals. Indeed, each individual in the model represents a large aggregation of biological individuals from the same species—e.g. local populations (Borthagaray, Cunillera-Montcusí, Bou, Biggs, et al., 2023; Worm & Tittensor, 2018).

The coalescent model consists of three main steps (Figure 1 Coalescent model): Step 1: initial species random sowing; Step 2: community filling until *Jj*; Step 3: calculation of community metrics. In step 1 the model randomly selects one individual from the regional pool for each one of the cells with J>0. Then, in step 2 an iterative filling process is started, in which for each iteration one individual is added in each cell with available space in function of three probabilities (Figure 1): a) self-recruitment, b) neighbouring dispersal and c) regional pool dispersal. The iterative filling continues until all available spaces in all cells have been filled (for the specific probabilities and detailed description of coalescent filling see Supplementary material 2 and in (Borthagaray, Cunillera-Montcusí, Bou, Biggs, et al., 2023). Finally, in step 3, when all ecoregion cells have been filled, we calculated for each cell its alpha diversity as the number of species and beta diversity as the average Jaccard dissimilarity index.

### Simulation of degradation scenarios

We defined 9 degradation levels from a 0% loss (i.e., original water surface) to 99% loss, includig intermediate values of 10%, 30%, 50%, 70%, 85%, 90% and 95%. For each level of degradation, a fraction of the of water surface was reduced along all cells concomitantly reducing cell’s community size *J*. When *J* falls below eight individuals, 2% of maximum community size, that cell was considered as a dry cell, *J*=0. Therefore, degradation scenarios implied habitat loss (i.e. overall community size decrease) but also cell loss and the concomitant fragmentation of the waterscape. The remaining cells with water become more and more distant from one another making more difficult the exchange of individuals between them.

We run several replicates of a coalescent model for each combination of dispersal abilities and degradation levels (7 dispersal abilities x 9 degradation levels= 63 scenarios),for each one of the 36 ecoregions and the 4 types of freshwaters. Then, for each ecoregion we obtained the mean alpha and beta diversity values and their maximum and minimum values. Based on the two metrics obtained, we calculated gamma diversity as: 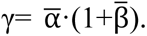. Finally, to make results comparable among ecoregions and organisms dispersal abilities, we scaled this gamma with respect to the gamma obtained when no waterscape degradation was involved (0% loss of water). We used this relative gamma diversity as a joint indicator of both alpha and beta diversities, summarizing for each degradation level, dispersal, and ecoregion the local and regional diversity interaction. This indicator was more informative than the real gamma diversity, which was constant in all simulations as the coalescent model did not incorporate species extinctions (Supplementary S2).

### Diversity decay parameters

The pattern of decay of regional diversity with the loss and fragmentation of habitat (Figure 1) was characterized by means of fitting a nonlinear model to each diversity decay curve: *γ*∼(*a* − *b* ∗ *f*) − exp (*f*^*q*^), where *γγ* is the relative gamma diversity; *aa* is the gamma diversity for the current waterscape estimated from alpha and beta diversities; *b* captures the average decline in diversity for each percentage of lost waterscape at the start of the degradation, hereafter *proportional decay rate*; *f* is the percentage of waterscape degraded; and *q* resumes the acceleration rate in diversity loss for each increase in the % of waterscape lost, hereafter *collapsing rate*. In summary, the *proportional decay rate* (*b*) is informing about the average trend in diversity loss, the lager the magnitude of *b* the stepper the species loss due to waterscape degradation. On the other hand, the *collapsing rate* (*q)* is informing on the acceleration of diversity loss along the habitat degradation gradient, higher values of *q* will imply a faster and steeper fall in diversity with each advance in waterscape degradation (i.e. a collapsing trend).

We retained the parameters *b* and *q* for characterizing the relationship between diversity loss and waterscape degradation for each ecoregion, dispersal ability, and freshwater type. Then, we represented these parameters across the continent to illustrate differences between ecoregions and identify more sensitive ones. Furthermore, for each ecoregion we calculated several structural descriptors summarizing ecoregion geographic location (i.e. mean latitude and longitude), ecoregion connectivity (i.e. the median and the coefficient of variation of betweenness centrality and outdegree centrality), ecoregion water cover (i.e. the mean and the coefficient of variation of water cover among cells) and ecoregion total size (i.e. number of cells with freshwater contained in the ecoregion). We obtained ecoregion geographic location from Pekel et al. (2008) databases. We obtained ecoregion connectivity metrics from Cunillera-Montcusí et al. (2023). We obtained the average water surface of ecoregions cells and the total ecoregion size (total number of cells in each ecoregion) from the water cover cells herein estimated. We related these ecoregion structural descriptors with the two diversity decay parameters using a generalized additive model (gam; mgvc R package; S. Wood & Scheipl, 2013) combined with a gam selection to identify significant descriptors and their trends. We conducted a model for each type of freshwaters and dispersal ability scaling descriptors with the *scale* function from the base R package (R-Core Team, 2019) but not including highly correlated descriptors (cutoff=0.8, in the *findCorrelation* function from caret R package; Kuhn et al., 2023).

## Results

### Alpha, beta, gamma results

The effect of waterscape degradation on diversity patterns was contingent along the considered dispersal abilities and ecoregions (Figure 2, 3, and Supplementary material 3). Overall, for the smaller dispersal abilities (e.g. 0.001 km and 0.1 km) low mean alpha and high beta diversities were observed, and the opposite was true for higher dispersal abilities (e.g. 10 km). This pattern was consistently observed along the four freshwater types considered (Figure 2). Along the gradient of habitat degradation all dispersal abilities were impacted decreasing their average alpha diversity and increasing their community differentiation, thus impacting regional gamma diversity (Figure 2 and supplementary material 4). The fitted nonlinear least squares model properly captured the loss of gamma diversity for all dispersal abilities. The higher organism’s dispersal ability, the greater the change in regional diversity due to waterscape degradation; a pattern generally observed throughout all freshwater types and ecoregions (Supplementary material 4). Despite this, some regions had a collapsing rate (q) equal to 0 implying that the loss of diversity was constant through the gradient of landscape degradation (Supplementary material 4). All simulation outputs as well as the corresponding code can be found online (Online supplementary) and all freshwater ecoregion plots reporting alpha, beta and relative gamma diversities can be found in Supplementary material 3 and 4.

**Figure 2:**
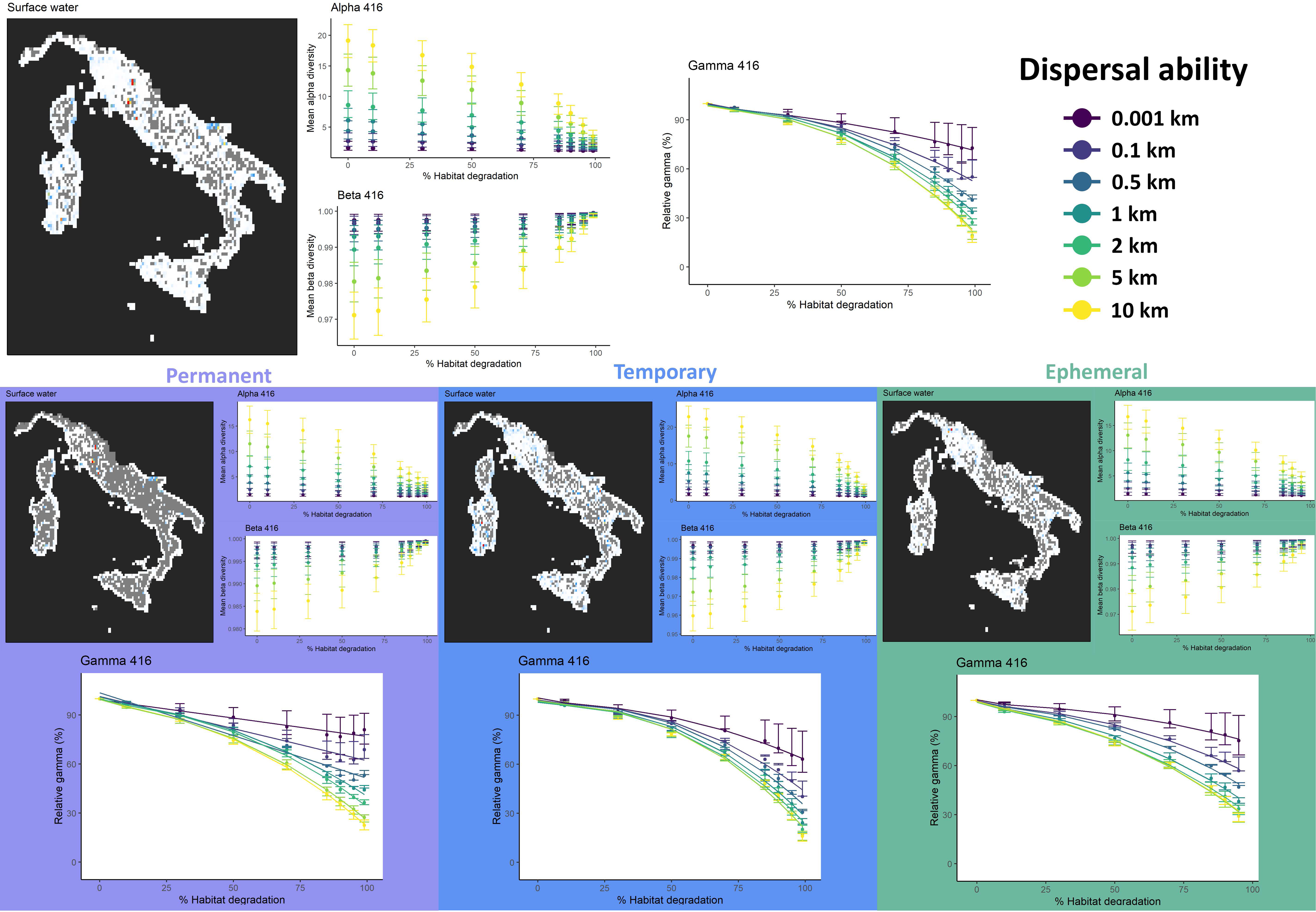
Example of the results obtained after the simulation for the ecoregion 416, Italian Peninsula & Islands and its four types of freshwaters: Total (top plot), Permanent (purple plots), Temporary (blue plots) and Ephemeral (green plots). We represent the database values for this region already converted in the grid of 10×10 km with cells with high surface water values in red colours and cells with low surface water values in blue-white colours, grey cells represent cells without that type of habitat. Alpha, Beta, and relative Gamma diversities (y axis) for the 7 dispersal abilities (coloured scatterplots) are represented along the landscape degradation gradient (% of habitat degradation).

### Diversity decay across the European continent

The diversity decay parameters *b* and *q* for each one of the 36 ecoregions and the 7 dispersal abilities are presented in the Supplementary material 5. Both the proportional decay rate (*b*) and the collapsing rate (*q*) are presented across ecoregions in Figure 3 and supplementary material 6. Overall, we observe an interesting heterogeneity in the decay parameters through the European freshwater ecoregions (Figure 3 and Supplementary material 6). Total freshwaters had an average *b* value of ™0.58 and *q* value of 0.34, permanent freshwaters a *b* value of ™0,56 and a *q* value of 0,34, temporary freshwaters a *b* value of ™0,58 and a *q* value of 0,34, and ephemeral freshwaters a *b* value of ™0,36 and a *q* value of 0,32. Some ecoregions indicated a high sensitivity to waterscape degradation, particularly captured in the proportional decay rate (*b*). For example, Vardar, situated across North Macedonia, Serbia, and Greece, had *b* values ranging around ™0.9 for total, ™0.6 for permanent, ™0.7 for temporary freshwater but around ™0.4 for ephemeral freshwaters. In the same line, Western Iberia, situated between Spain and Portugal, had *b* values of ™0.7 for total, ™0.6 for permanent, ™0.6 for temporary, and ™0.4 for ephemeral freshwaters. On the other hand, the collapsing rate (*q*) ranged between 0.35 and 0.25. It should be highlighted that the *q* parameter is a potential exponent within an exponential term, therefore small differences in the parameter values can produce large changes in the shape of the curve. Ecoregions from Norwegian Sea Drainages, and the Northern Baltic Drainages, spanning Norway, Sweden, Finland, and Russia had relatively large *q* values (i.e. ranging around 0.348 for all freshwater types). Overall, ephemeral systems presented smaller proportional decay rates than the other freshwaters (Figure 3 and supplementary material 6) but not for collapsing rates. Furthermore, the differences among dispersal abilities were quite small and they generally highlighted similar regions as the more sensitive, being higher dispersal abilities the ones mostly experiencing greater diversity decay parameters, with the exception of the smallest dispersal abilities (i.e. 0.001 and 0.1 km).

**Figure 3:**
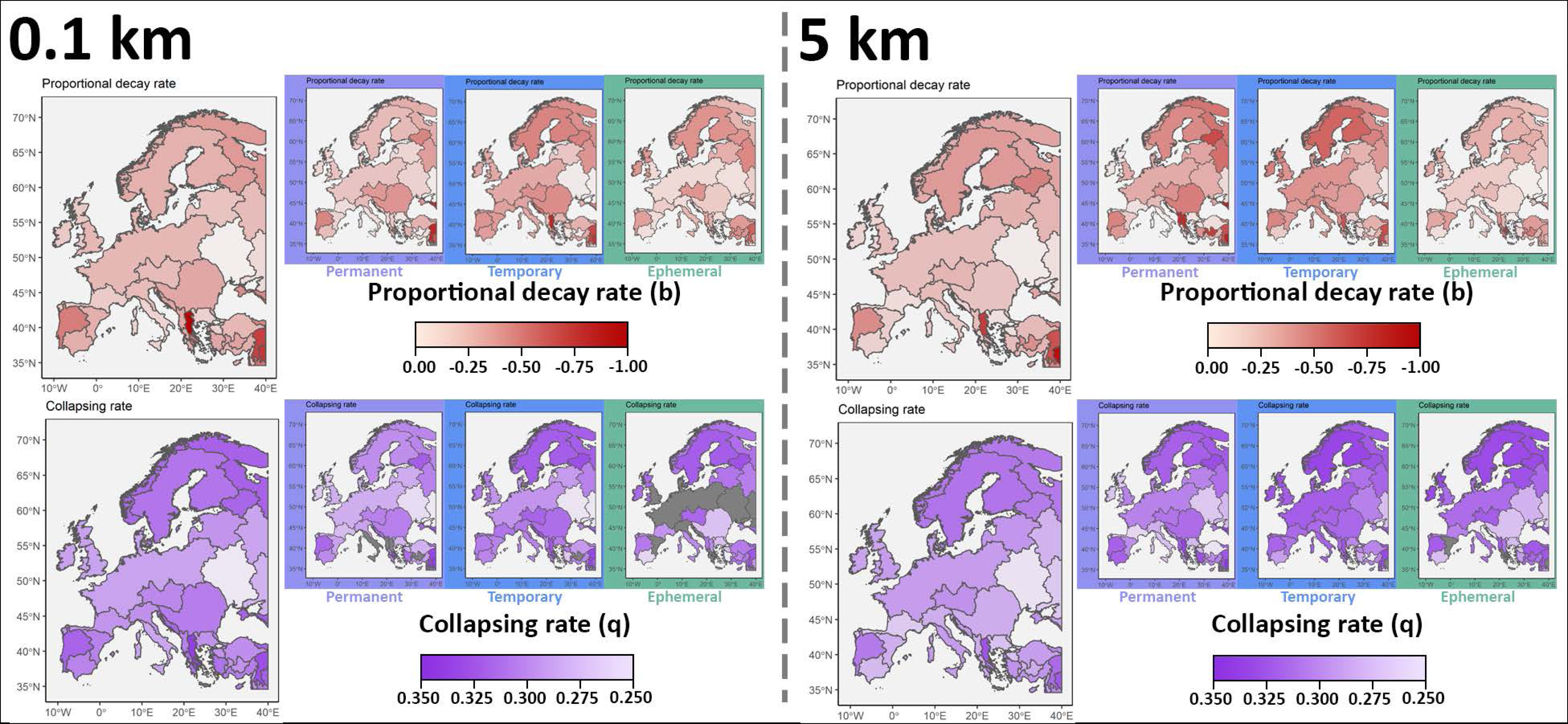
European ecoregions diversity decay parameters for dispersal abilities of 0.1km (low dispersal capacity) and 5 km (high dispersal capacity). From left to right, Total freshwater, Permanent freshwaters (purple background), Temporary freshwaters (blue background) and Ephemeral freshwaters (green background). First plot row represents the proportional decay rate (b) in red gradient colours and the second row represents collapsing rates (q) in purple colours.

### Diversity decay trends in relation to ecoregion structural descriptors

We could observe some general common trends across the different dispersal abilities despite the presence of strong differences among the significant structural descriptors for each freshwater type (Figure 4 and Supplementary material 7, 8, 9). Total freshwaters *b* and *q* values appeared related with connectivity (median betweenness and median out degree) and water cover (mean water cover and water cover variation; Figure 4A). For permanent freshwaters, water cover drivers were the main significant ones, although variation in betweenness also showed a significant positive pattern (Figure 4B). Contrastingly, for temporary freshwaters only median out degree appeared as a main significant trend for *b* and *q* values (Figure 4C). Finally, ephemeral freshwaters diversity decay parameters appeared related to ecoregions geographic location (latitude), water cover (mean water cover), connectivity (median out-degree) and ecoregion size structural drivers (Figure 4D).

**Figure 4:**
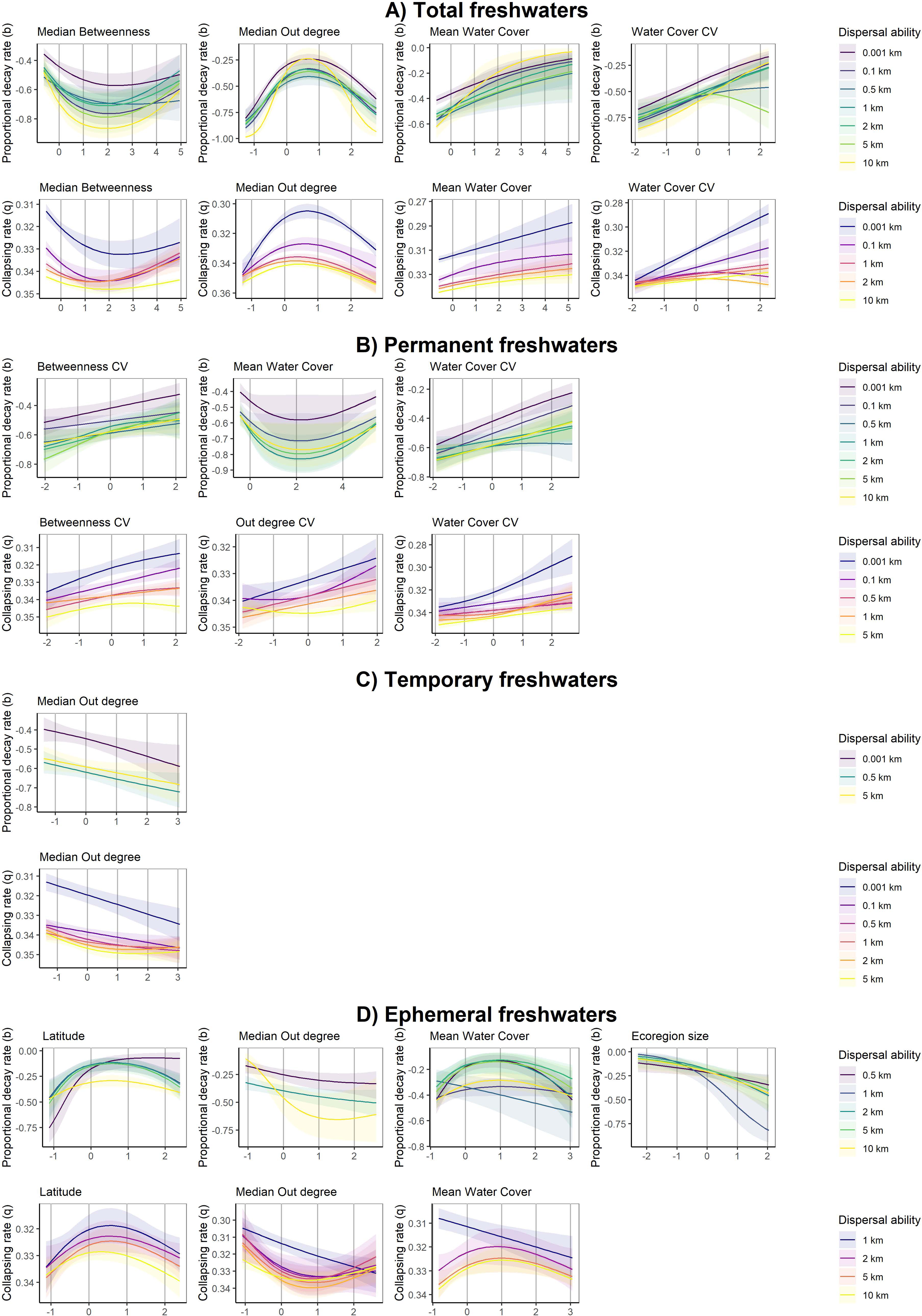
Landscape structural descriptors main significant trends against the two diversity decay parameters Proportional decay rate (top plots) and Collapsing rates (bottom plots) for each of the four freshwater types: A) Total freshwaters, B) Permanent freshwaters, C) Temporary freshwaters, D) Ephemeral freshwaters. Dispersal abilities are identified with different colour gradients from low (0.001km) to high (10km) with purple-green-yellow gradient for proportional decay rate trends and blue-red-yellow for collapsing rates to ease visual identification. Only significant trends between dispersal abilities (individual lines) and the scaled landscape structural descriptors are represented (x axes). Note that y axes are ordered differently ranging from 0 to ™1 for proportional decay rate (*b*) and from 0.2 to 0.4 for collapsing rate (*q*) to visually define high resistance to landscape degradation (decay rates close to 0 for both cases) as “high” y axis values and low resistance to landscape degradation as “low” y axis values.

## Discussion

Multiple drivers are causing the current biodiversity crisis (He et al., 2019; Maasri et al., 2022; Reid et al., 2019), as well as their complex interactions (Beermann et al., 2018; Jackson et al., 2016; Nuy et al., 2018). This complexity hinders the mechanistic understanding of how these drivers impact freshwater ecosystems (Reid et al., 2019). Among them, waterscape degradation — habitat loss and fragmentation— plays a key role and yet we are still far from comprehending its connection with diversity patterns (Albert et al., 2021; Des Roches et al., 2021; Grzybowski & Glińska-Lewczuk, 2019; Horváth et al., 2019). In this work, we precisely aimed to advance in the potential consequences of waterscape degradation on freshwater diversity and summarise the differences in the sensitivity among European ecoregions. For all ecoregions, waterscape degradation determined a decrease in alpha diversity, an increase in beta diversity, and a general loss in gamma diversity. Such generalized patterns mimic the reported consequences of landscape degradation (Halley & Iwasa, 2011; Horváth et al., 2019; Kuipers et al., 2021; Segan et al., 2016). Nevertheless, we saw how these decay patterns differ along the continent with more sensitive regions (e.g. Vardar or Western Iberia) presenting more pronounced decays. Such variation stands as the main of our contributions: the quantification of region-wide diversity response to waterscape degradation and the identification of potentially more sensitive ecoregions (Supplementary table 5). We all accept that waterscape degradation will decrease diversity, yes, but at which rate we will be losing species as landscapes are degraded? Here we gave a first order approximation to answer this question for the European continent ecoregions. Overall, European landscapes lose more than 0.5% of their diversity for every 1% of habitat that is eliminated. But this value increases around 0.3% for every 1% of habitat lost. This implies that after a 10% of freshwater loss, 12 species of every 100 would be lost only due to waterscape degradation.

The two decay parameters inform about the interaction between dispersal ability and waterscape structure and how they drive diversity decay in each ecoregion. For example, species with relatively high dispersal abilities had higher diversity values than low dispersal abilities before waterscape degradation. However, as landscape degraded, diversity values of high dispersers became similar to those of low dispersers indicating the concomitant regional isolation of high dispersers. Thus, species having greater dispersal abilities had greater decay rates because the landscape that was sustaining their diversities fragmented diminishing their values (Cañedo-Argüelles et al., 2015; Cunillera-Montcusí et al., 2021; Phillipsen et al., 2015; Riibak et al., 2017). On the other hand, species with low dispersal were formerly less dependent on connectivity (Borthagaray, Berazategui, et al., 2015; Borthagaray et al., 2012; Urban & Keitt, 2001), and consequently, although their diversity values also decayed, they did it at lower rates. For many ecoregions (e.g. Central and Western Europe, Gulf of Venice Drainages, or Lower Tigris & Euphrates), species with relative low dispersal presented a constant rate in diversity loss without evidence of acceleration in diversity decay with the increase in waterscape degradation. This could be related to a loss of diversity only related to habitat loss (i.e. habitat surface; Camelier & Zanata, 2015; Hof et al., 2008; Lomolino, 2001; Oertli et al., 2002), without the synergetic effect of a reduction in connectivity that experience organisms with larger dispersal abilities. Such patterns identified the higher vulnerability of wider range organisms against landscape degradation (Arntzen et al., 2017; Beebee & Griffiths, 2005; He et al., 2019).

Ecoregions had marked differences in their level of sensitivity to landscape degradation as the presence of freshwater habitats and their distribution varied across them. For example, permanent and ephemeral habitats were differently distributed in the Dnieper–South Bug than in Southern Anatolia (Abell et al., 2008). Indeed, ephemeral waters had globally smaller diversity decay values in comparison to temporary and permanent. Such differentiation was linked to two main processes. First, ephemeral waters presented in general smaller surfaces in each ecoregion which limits the number of species that they were able to contain (R. A. Chisholm et al., 2018; Lomolino, 2001). Second, these freshwaters were widely widespread being more connected among them and consequently being more homogeneous but also robust to landscape degradation (Cunillera-Montcusí et al., 2021; Gao et al., 2016; Santos et al., 2021). These two elements generated smaller diversity decay rates as communities with not many species needed greater levels of degradation to decay. Contrastingly, permanent habitats had a less widespread distribution through the ecoregions, which generated the opposite pattern, with permanent freshwaters experiencing the greatest decay values. Temporary systems had similar decay rates than permanent freshwaters, even though they were generally more widespread. These diverging patterns highlight the relevance of also considering different types of habitats when assessing wide-scale diversity (Borthagaray, Cunillera-Montcusí, Bou, Biggs, et al., 2023). Our framework relies on the distinction of habitats linked to their drying frequency (Pekel et al., 2016), but it aims to capture the overall diversity provided by the way in which these habitats are distributed (Borthagaray, Cunillera-Montcusí, Bou, Tornero, et al., 2023; Faggioni et al., 2021; Worm & Tittensor, 2018). Such approximations have been already proposed for habitats within a determined studied regions (e.g. a natural park as in Carroll et al. (2018), pondscape as in Borthagaray, Berazategui, et al. (2015) or an ecoregion as in Kuipers et al. (2021). At the end, how the spatial arrangement of habitats in a region is sustaining a determined level of diversity is tightly linked to how the network is structured (Borthagaray, Cunillera-Montcusí, Bou, Tornero, et al., 2023; Drake et al., 2017; Economo & Keitt, 2008, 2010; Santos et al., 2021). Nevertheless, in the current study we went a step further and explored these patterns on a degrading landscape, which allowed us to quantify the global consequence of structural network breakage. Such approximation, with the calculation of decay parameters, summarized a dynamic process by distilling landscape degradation consequences on diversity in two specific parameters: the proportional decay (b) and collapsing (q) rates. Thus, our work goes beyond a static approximation, opening the door to incorporating more dynamic and realistic scenarios with the aim of understanding the general response patterns to dynamic processes (Hubbell, 2001; Worm & Tittensor, 2018)

Differences between ecoregions response to degradations were associated with original features of their landscapes. In general, ecoregions with greater values of water coverage and with greater spatial variation in this cover were more resistant to landscape degradation. In addition, ecoregions having an heterogeneous distribution of water through the landscape (i.e. variation in centrality), were also more resistant to landscape degradation (Borthagaray, Cunillera-Montcusí, Bou, Tornero, et al., 2023; Savary et al., 2023). This link with heterogeneity emphasizes the key role of centrality patterns in resistance to disturbances (Carroll et al., 2018; Gastauer et al., 2021; Kuipers et al., 2021). Heterogeneity in the distribution of habitats could favour the existence of a gradient of spatial relevance throughout the landscape, which could concomitantly favour the coexistence of different strengths on assembly mechanisms (e.g. mass-effects in more central habitats, priority effects in more isolated habitats) and therefore foster a more diverse and imbricated system with higher global resistance to degradation (Altermatt et al., 2020; Leibold & Chase, 2018; Ritchie, 2009). Thus, to also considering coefficients of variation seems relevant to better capture network resilience (Cunillera-Montcusí et al., 2021; Economo & Keitt, 2010; Fagan, 2012; Kuipers et al., 2021). All these insights as well as the obtained diversity decay parameters could be directly incorporated in management and conservation assessments constituting a valuable resource to set priority guidelines at European level (Cunillera-Montcusí et al., 2023; Maasri et al., 2022; Reid et al., 2019). Furthermore, the approximation and the decay parameters used in the current study represent a novel way through which we could assess these values at other landscape scales (e.g. countries) or systems (e.g. only fluvial systems) that opens a promising window for conservation and management as it incorporates regional scale and metacommunity-based dynamics (Chase, Jeliazkov, et al., 2020; Cid et al., 2020; Gounand et al., 2018).

The approximation we developed to assess landscape degradation impacts on diversity is of high value as far as it is complemented with other views, as in this work we focused on landscape degradation as a driver of diversity loss. Indeed, many other drivers determine aquatic diversity and its loss (Araújo et al., 2006; Didham et al., 2020; He et al., 2019; Seibold et al., 2019; Wagner et al., 2021). For example, fluvial fragmentation, change in land cover, urbanization, pollution, and species invasions, which were not explicitly included in the present analyses (Belletti et al., 2020). The contribution of the current results relies on their contextualization with other available information, for example climatic vulnerability index (Edmonds et al., 2020), natural parks protection (Rivers-Moore et al., 2016; Santini et al., 2016), vulnerable species (Angelibert & Giani, 2003; Marini et al., 2019) or human impacts (Hintz & Relyea, 2017; Reid et al., 2019). Therefore, the patterns herein reported must be understood as another layer to consider on conservation planning (Chase, Jeliazkov, et al., 2020; Cid et al., 2020), where landscape degradation has seldom been considered in spite of being a recognized driver of diversity loss (Kuipers et al., 2021). Therefore, we aimed to capture the pattern caused by landscape degradation and provide general indicators informing about its impacts in biodiversity patterns, as well as where it may have greater impacts (Worm & Tittensor, 2018).

The links between landscape structure, dispersal patterns, and biodiversity architecture and stability remain to be fully unfolded at the theoretical and empirical levels. However, nowadays there is an imperious need to better comprehend it for aquatic landscapes as they are being largely impacted by human intervention (Gozlan et al., 2019; IPCC, 2022; Tuytens et al., 2014). Future assessments at the European level should consider ecoregion sensitivity to focus conservation efforts in areas with higher sensitivity to landscape degradation. These ecoregions might greatly benefit from interventions favouring freshwater connectivity (e.g. addition of stepping-stone habitats), and particularly favouring their overall waterscape spatial heterogeneity. Thus, both mean and variation coefficients from network indices should be considered in conservation planning (Barnett & Belote, 2021; Borthagaray, Cunillera-Montcusí, Bou, Tornero, et al., 2023; Santini et al., 2016; Savary et al., 2023). At the end, this pushes forward the need to consider the overall network structure in order to sustain systems functioning despite the loss of habitat (Gonzalez et al., 2017; Marini et al., 2019). In this work, we did a first step towards a wide-scale quantification of landscape degradation impacts. Some efforts have already been proposed in these line (Fagan, 2012; Horváth et al., 2019; Rodeles et al., 2021), but there is still work to do in order to fully unfold landscape degradation impacts to face global change challenges (Parra et al., 2021; Reid et al., 2019; Segan et al., 2016). Future works could benefit from the current approximation to design more realistic scenarios considering anthropogenic landscape alterations (e.g. urbanisation, agricultural landscapes, protected areas) to modulate fragmentation levels. Overall, by incorporating these new perspectives, we were able to incorporate new layers of understanding for aquatic habitat conservation that might be key to overcome conservation current halts (Haase et al., 2023) and that may play a key role in the preservation of European, and global, waterscapes against landscape degradation and the concomitant diversity crisis.

## Acknowledgments

This study was funded by H2020 EU-funded project PONDERFUL (869296), CSIC I+D 2020_ID_188 and CSIC groups (ID 657725) UDELAR, and FCE_1_2021_1_167009. DCM was supported by the European Union - NextGenerationEU, Ministry of Universities and Recovery, Transformation and Resilience Plan, through a call from Universitat de Girona and from the European Union’s Horizon 2020 research and innovation programme under the Marie Skłodowska-Curie grant agreement No 101062388. The authors do not have any conflict of interest to declare. We would like to express our gratitude to Felipe Maresca for his valuable comments that have significantly improved the readability of our manuscript.

## SUPPLEMENTARY MATERIAL

**Supplementary material 1:** Table with Ecoregions names and reference.

| **Supplementary material 1: Table with Ecoregions names and reference.** |  |
| --- | --- |
| **ID** | **Ecoregion name (Abell et al. 2008)** |
| 436 | Coastal Levant |
| 440 | Arabian Interior |
| 433 | Western Transcaucasia |
| 426 | Crimea Peninsula |
| 420 | Southeast Adriatic Drainages |
| 422 | Vardar |
| 431 | Central Anatolia |
| 429 | Western Anatolia |
| 423 | Thrace |
| 414 | Eastern Iberia |
| 409 | Lake Onega Lake Ladoga |
| 416 | Italian Peninsula & Islands |
| 403 | Cantabric Coast Languedoc |
| 417 | Upper Danube |
| 405 | Norwegian Sea Drainages |
| 408 | Southern Baltic Lowlands |
| 425 | Dnieper South Bug |
| 406 | Northern Baltic Drainages |
| 437 | Orontes |
| 428 | Kuban |
| 441 | Lower Tigris & Euphrates |
| 421 | Ionian Drainages |
| 424 | Aegean Drainages |
| 419 | Dalmatia |
| 442 | Upper Tigris & Euphrates |
| 432 | Southern Anatolia |
| 415 | Gulf of Venice Drainages |
| 413 | Southern Iberia |
| 430 | Northern Anatolia |
| 412 | Western Iberia |
| 427 | Don |
| 402 | Northern British Isles |
| 422 | Volga Ural |
| 418 | Dniester Lower Danube |
| 407 | Barents Sea Drainages |
| 404 | Central & Western Europe |

**Supplementary material 2:** Coalescent model specific steps described in Figure 1.

We assigned all freshwater surfaces defining community size to each cell, *J*_*i*_. This sets the number of individuals in each specific cell based on the area occupied by water, *A*_*j*_

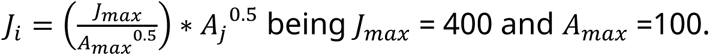

Once we defined J for each cell, communities were filled until *J* individuals. **In** figure 1 **step 1**, one individual is added in each one of the communities being selected from the pool. This step is necessary to initially sow the communities and allow the other steps to take place.

In figure 1 step 2, individuals are added to each community being sampled from three main sources: 1) from the metacommunity species pool, 2) from neighbour communities, or 3) from the same local community. The probabilities that define this process are *m*. *p*ool = 0.01) (Worm & Tittensor, 2018), *m*. neighbour, and 1 − (*m*. *p*ool+ *m*. *n*eihbour), respectively. Neighbouring dispersal is defined by a dispersal kernel between each pair of communities determining the *m*. *n*eghibour. This value is modelled with an exponential decay function as *m*. *n*eighbour= *m*. *m*ax∗ e^−*b*∗*di*j^; being *d*_*ij*_ the Euclidean distance between communities *i* and *j, m*. *m*ax the migration between ponds 0 kilometres apart (*m*. *m*ax = 1) and *bb* a migration parameter defined by the distance between ponds at which migration decays to half its maximum value, *dd*_50_. Using these dispersal processes, a community-by-community migration matrix *M* was then estimated, with elements representing the dispersal rate from community *i* to community *j*. Self-recruitment was fixed as one setting the diagonal of the migration matrix *M* to one. The incoming recruitment probabilities, *m*, for a local community are represented by the columns of the migration matrix *M* plus the migration from the pool. These values are standardized to add 1, as 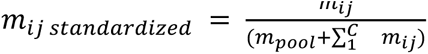 being *C* the number of communities.

The pool of neighbour species available to colonize a local community was then estimated as the matrixial product of the metacommunity abundance matrix and the standardized migration matrix. The matrix implementation of the coalescent algorithm allows to efficiently simulate the assembly of large metacommunities. This step is done until all communities (i.e. cells) have been filled with J individuals. For all simulations, the species pool was defined at 200 species.

Finally, in step 3 mean alpha and beta diversity are calculated from all ecoregion communities in each simulation. We run 10 replicates for every scenario considering a freshwater type (i.e. total, permanent, temporary and ephemeral), a determined level of landscape degradation (i.e. from 0 to 99% loss) and dispersal ability (from 0.001 km until 10 km). This makes a total of 252 simulations that were run 10 times. The median of mean ecoregion alpha and beta diversity was the final value used to later calculate gamma diversity.

All code and corresponding functions can be found in the following GitHub repository: https://github.com/Cunillera-Montcusi/PanEU_fragmentation

**Supplementary material 3:**
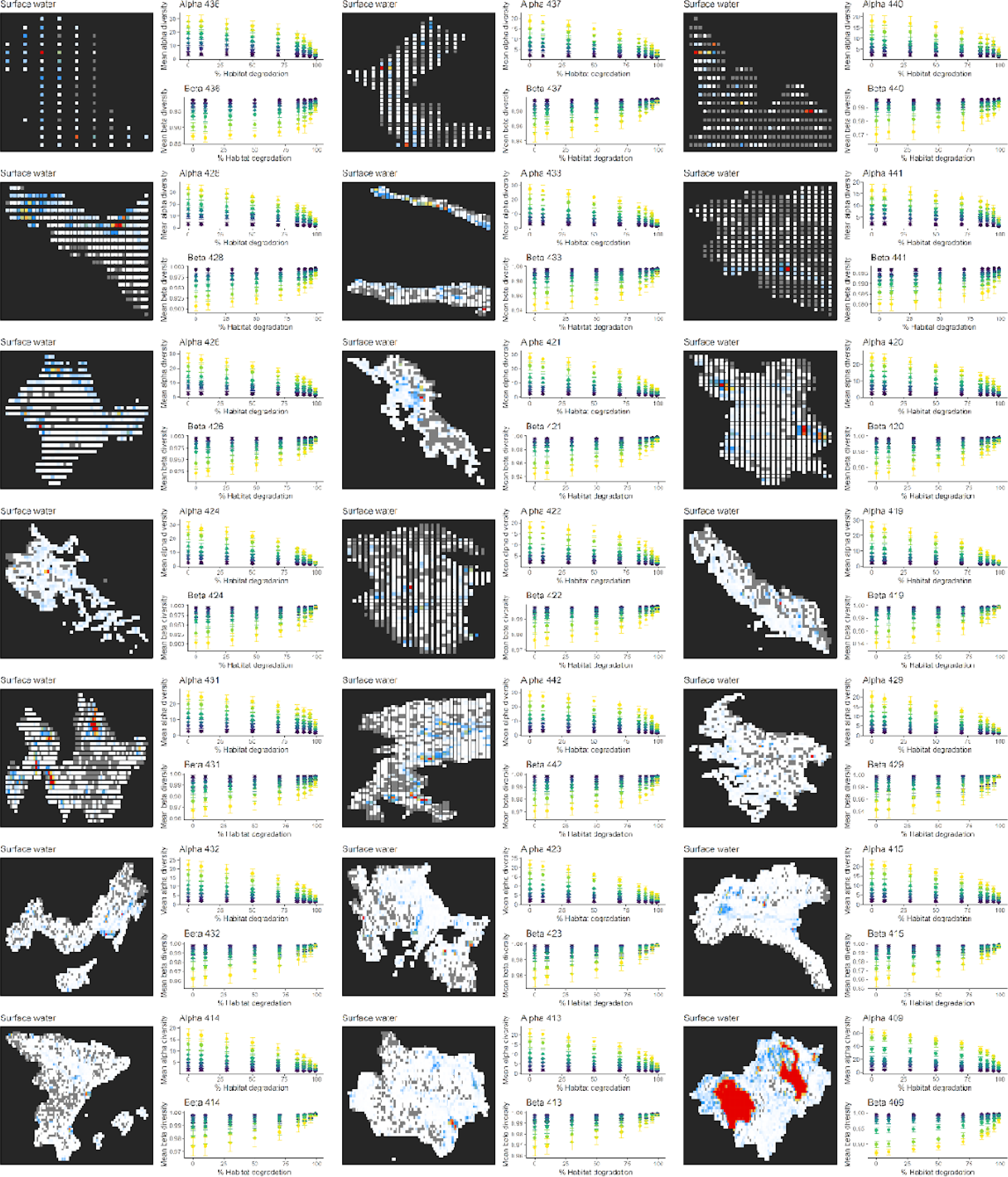
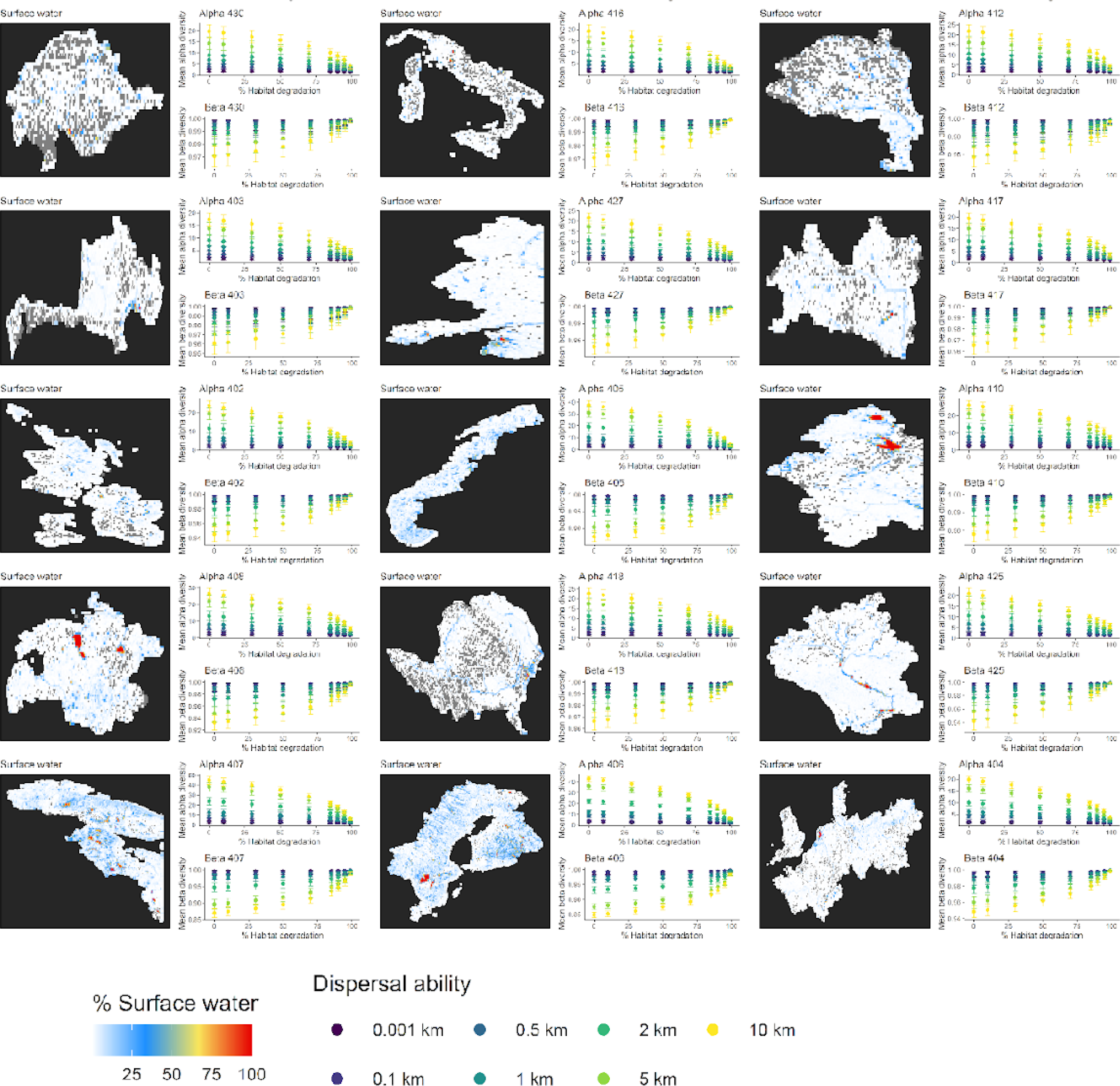
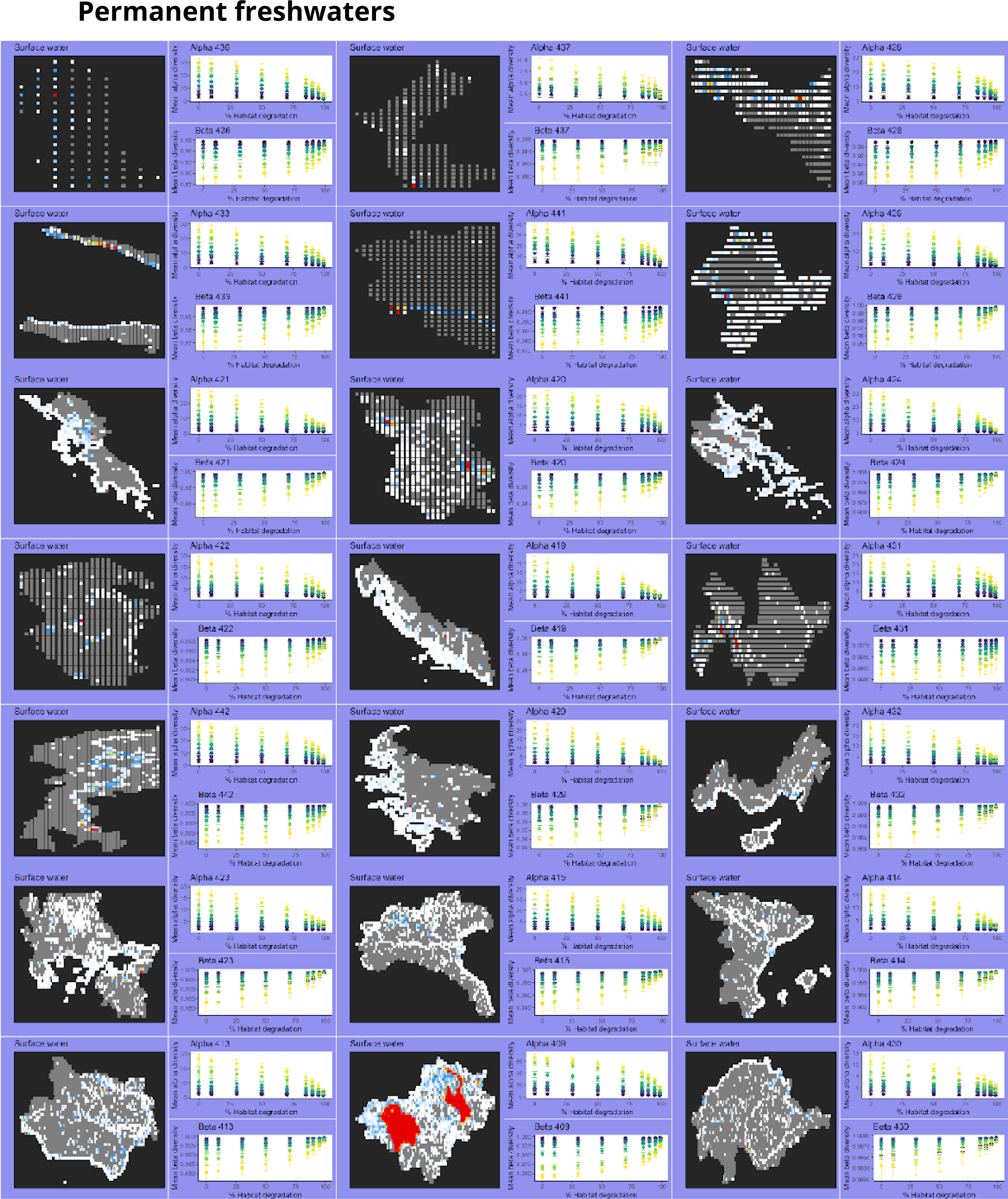
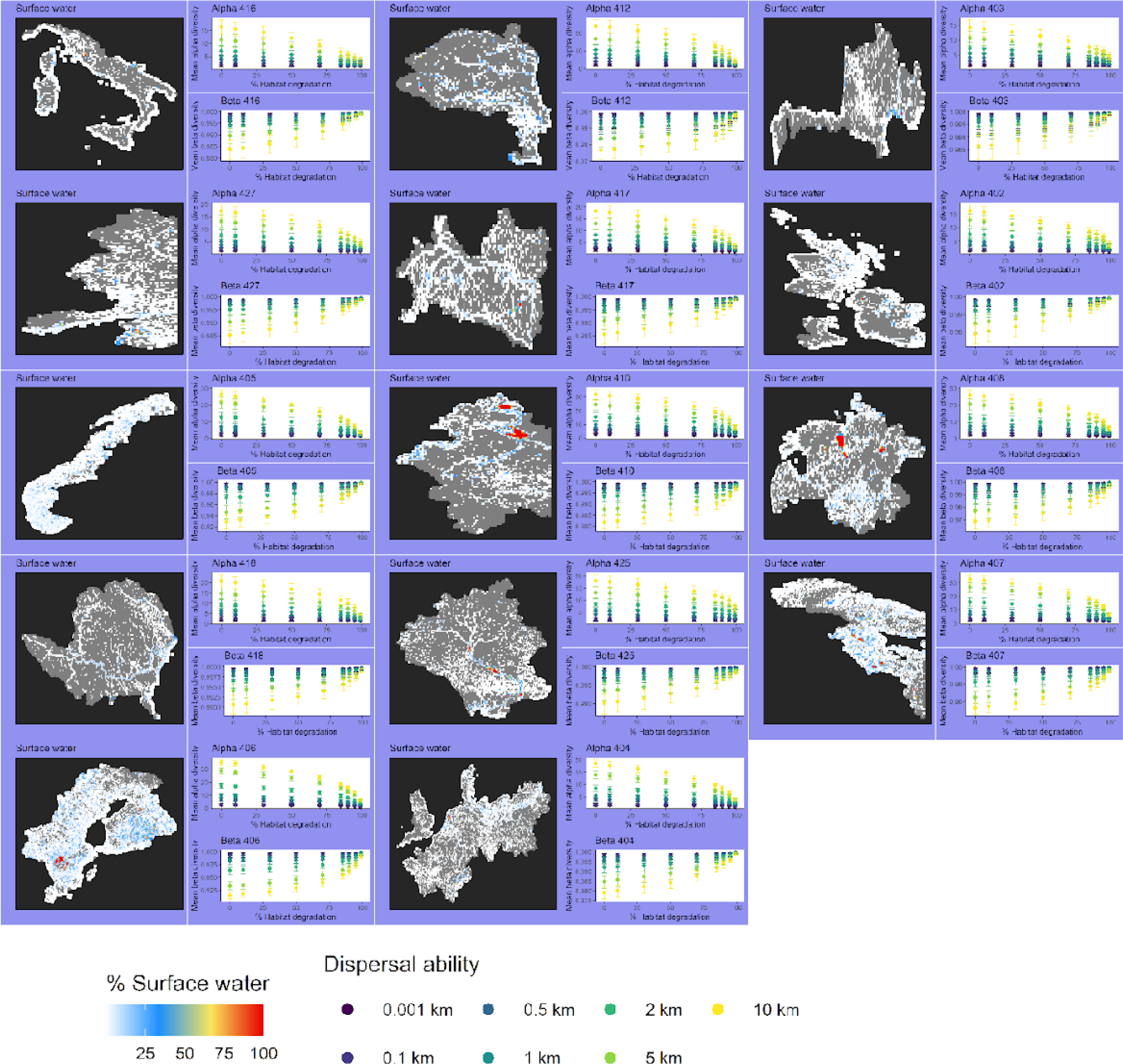
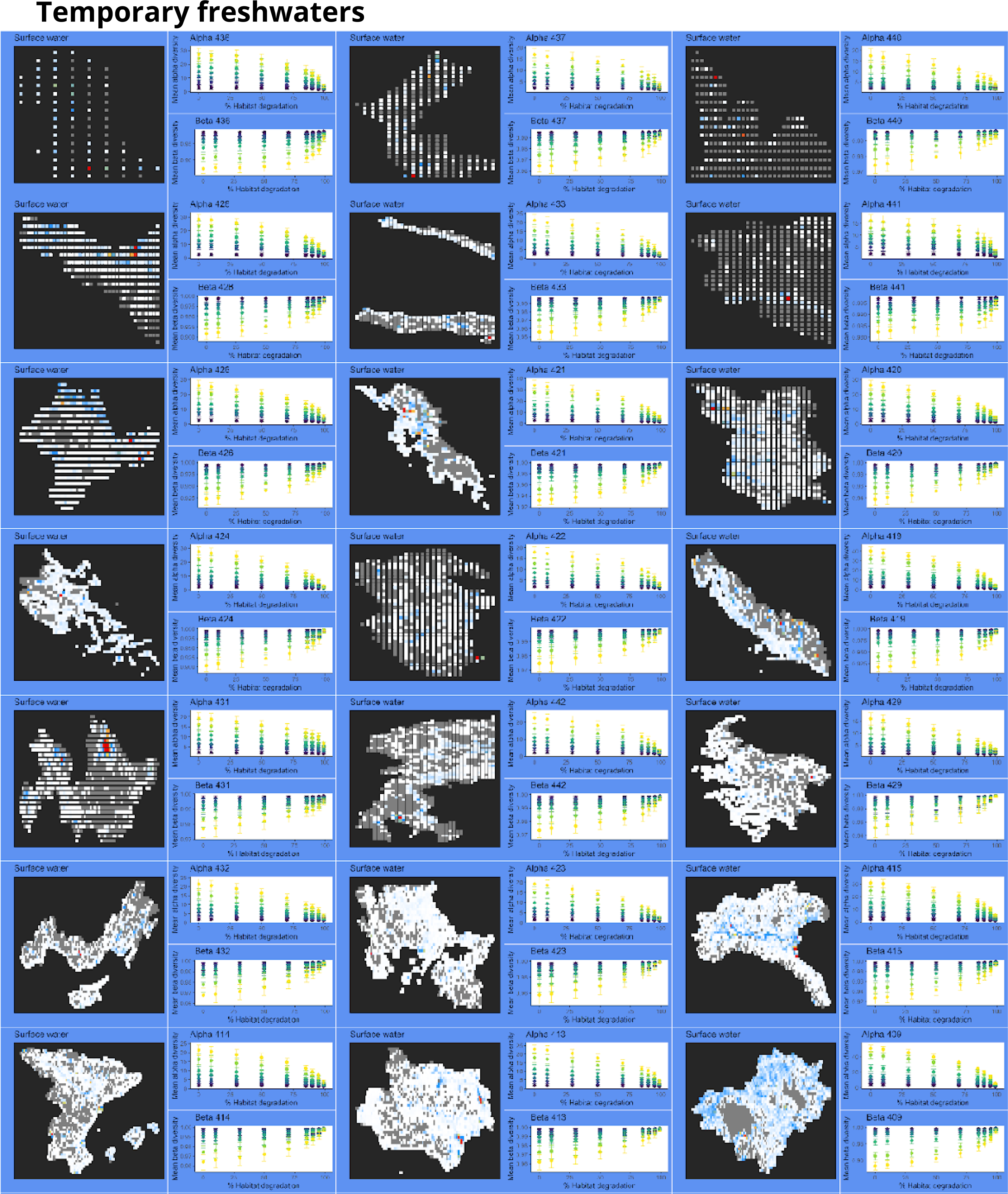
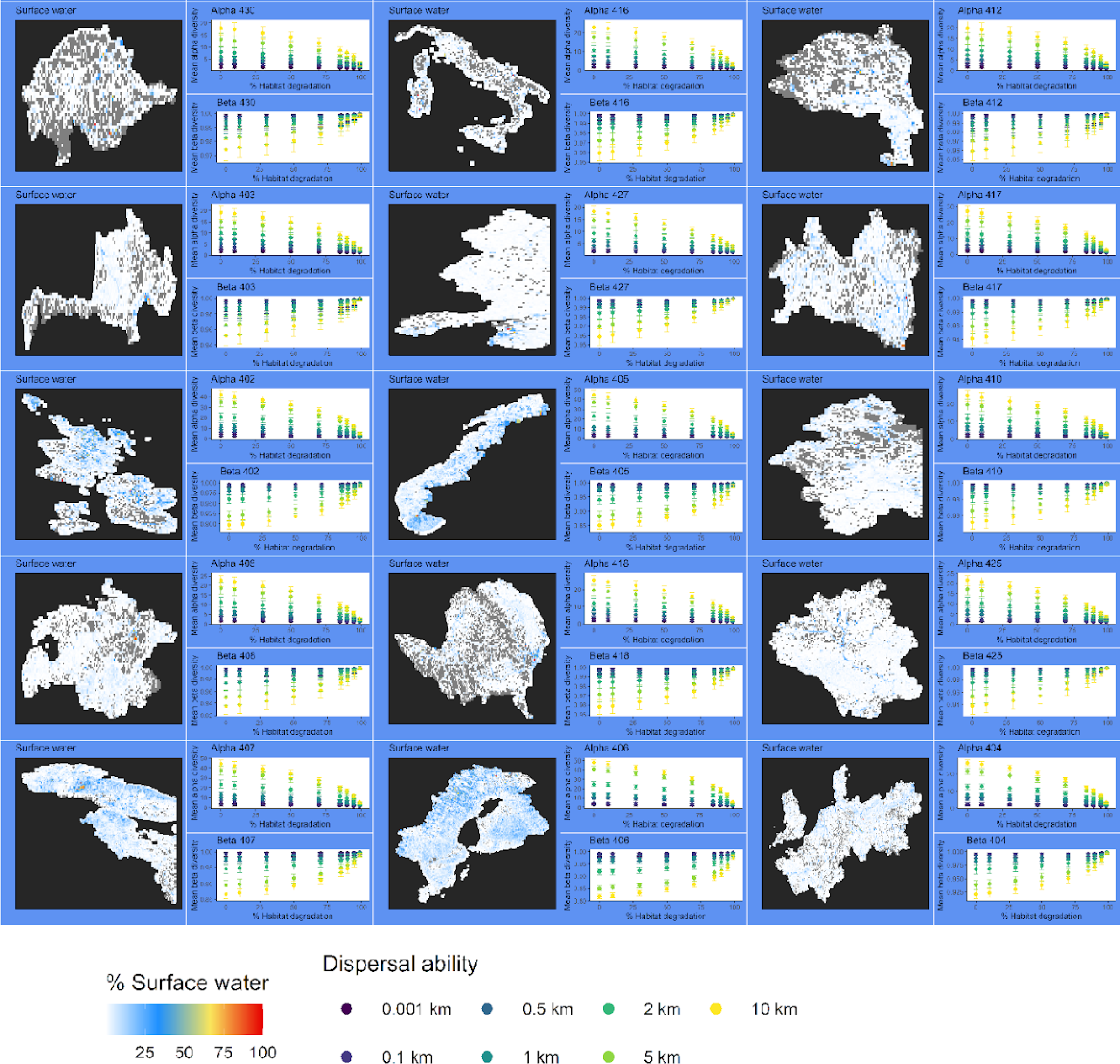
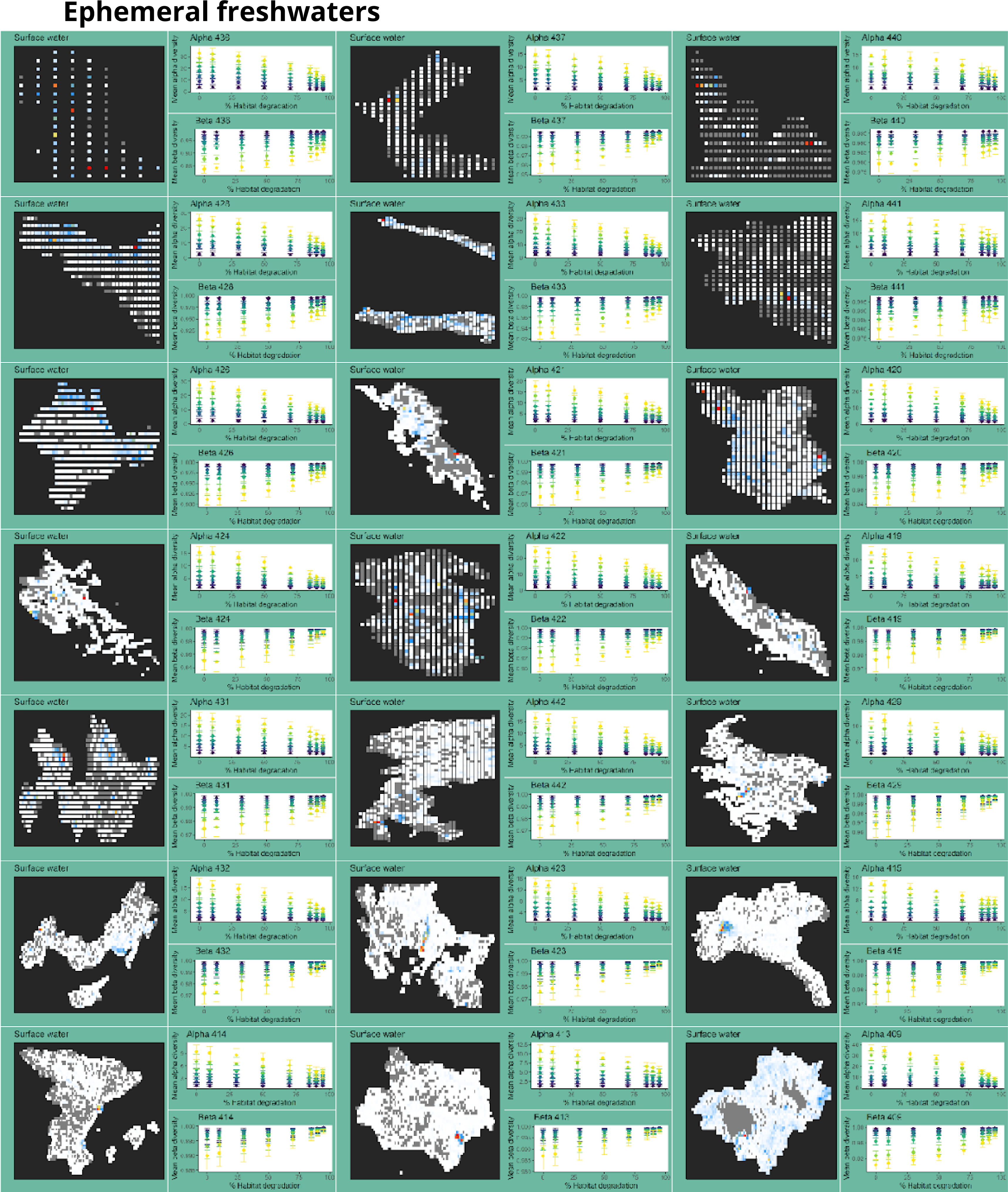
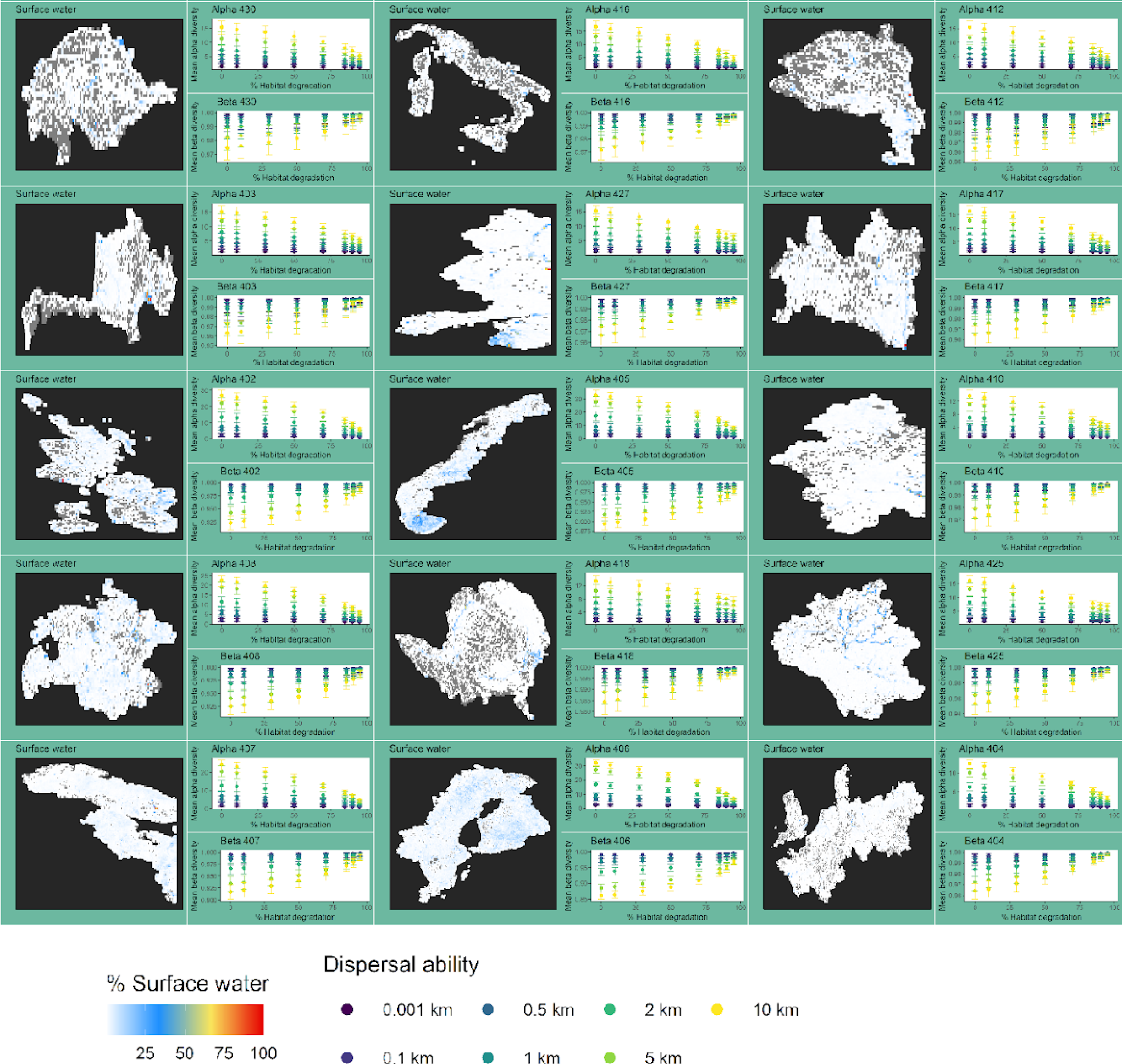
Ecoregion maps with all contained cells and their corresponding occupied surface values. Red cells indicate almost full of freshwater cells, grey cells indicate empty cells. Next to each ecoregion, mean alpha and beta diversity results for each dispersal ability (colours from yellow to purple) and along landscape degradation values are plotted. These results are shown for Total (white background), permanent (purple background), temporary (blue background) and ephemeral freshwaters (green background).

**Supplementary material 4:**
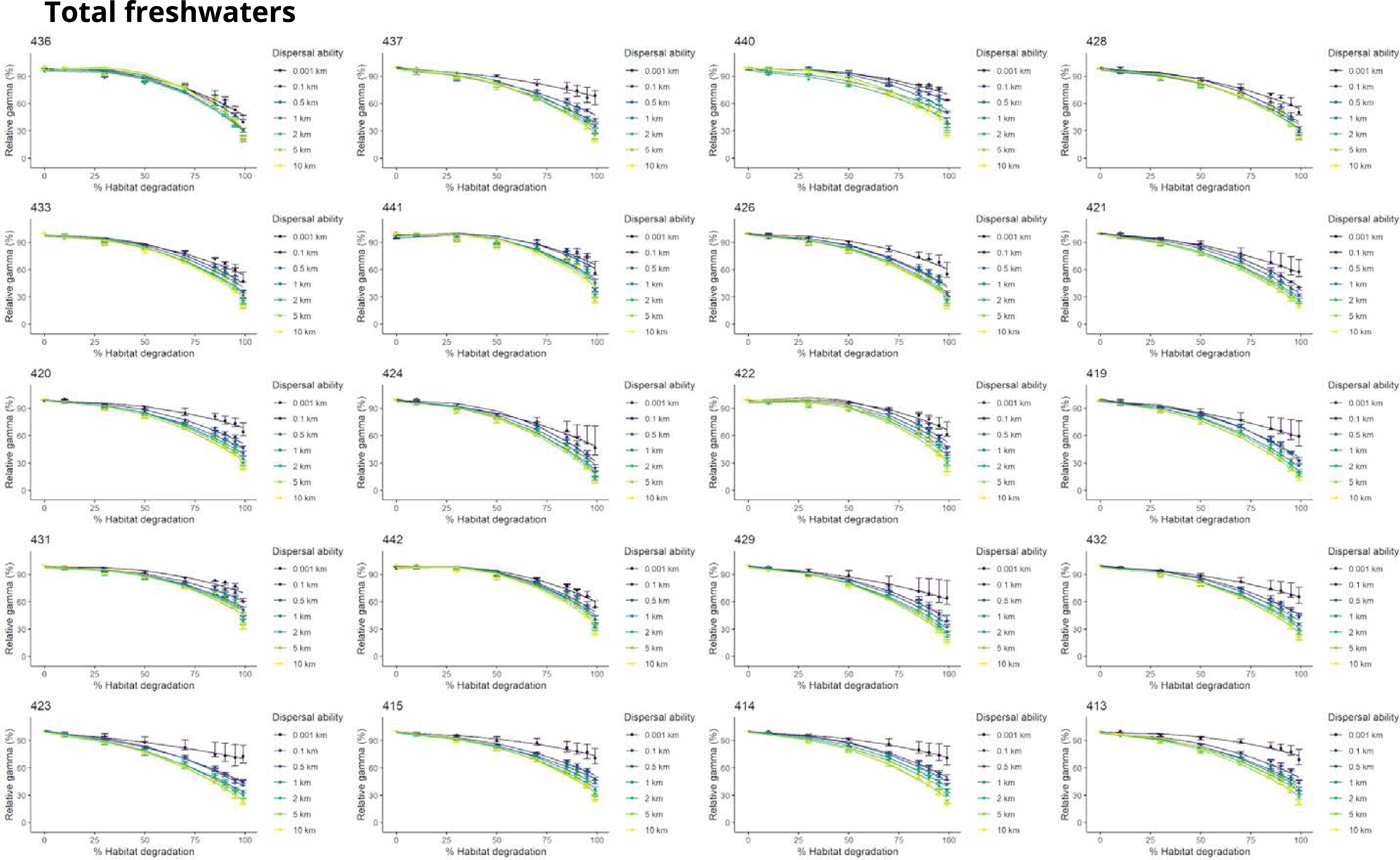
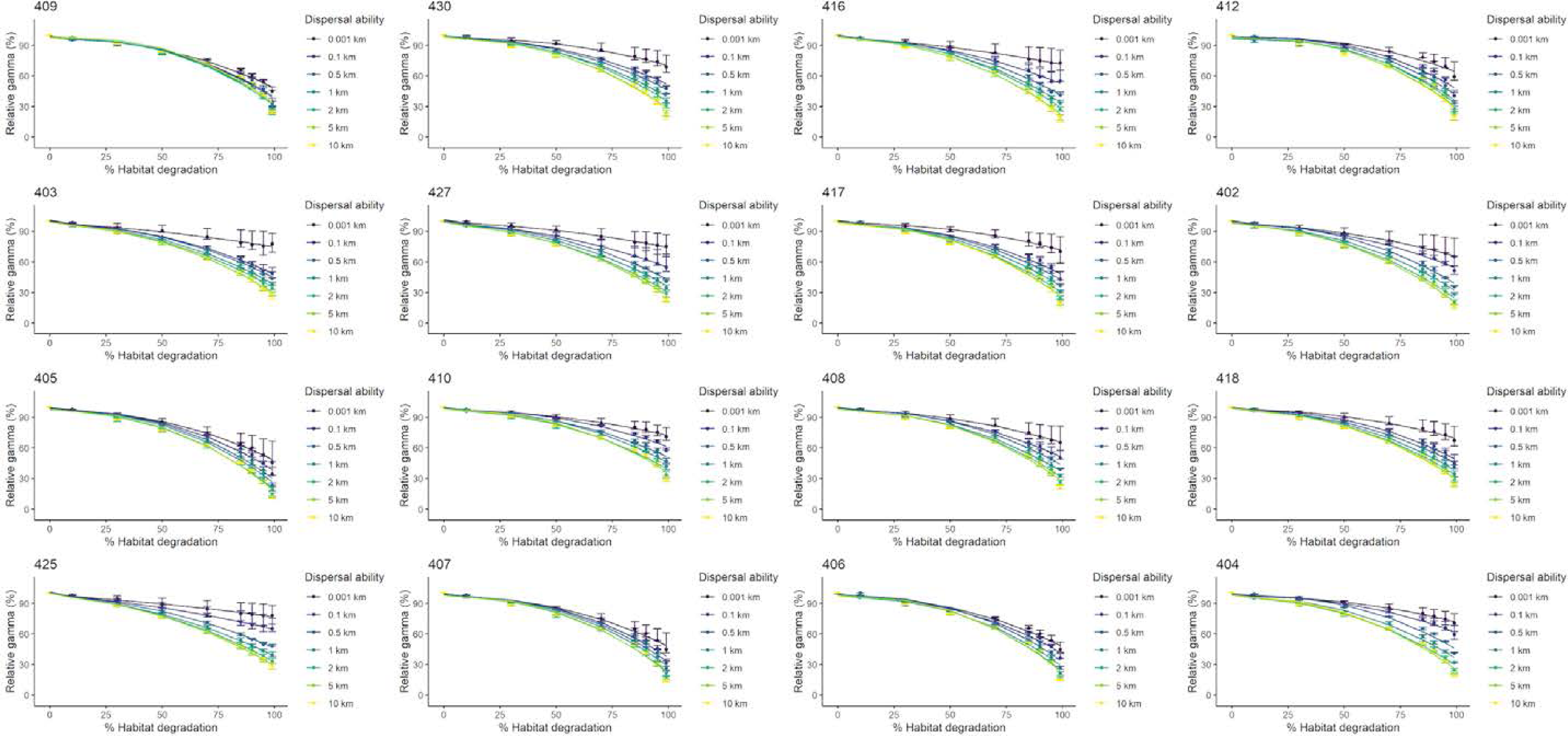
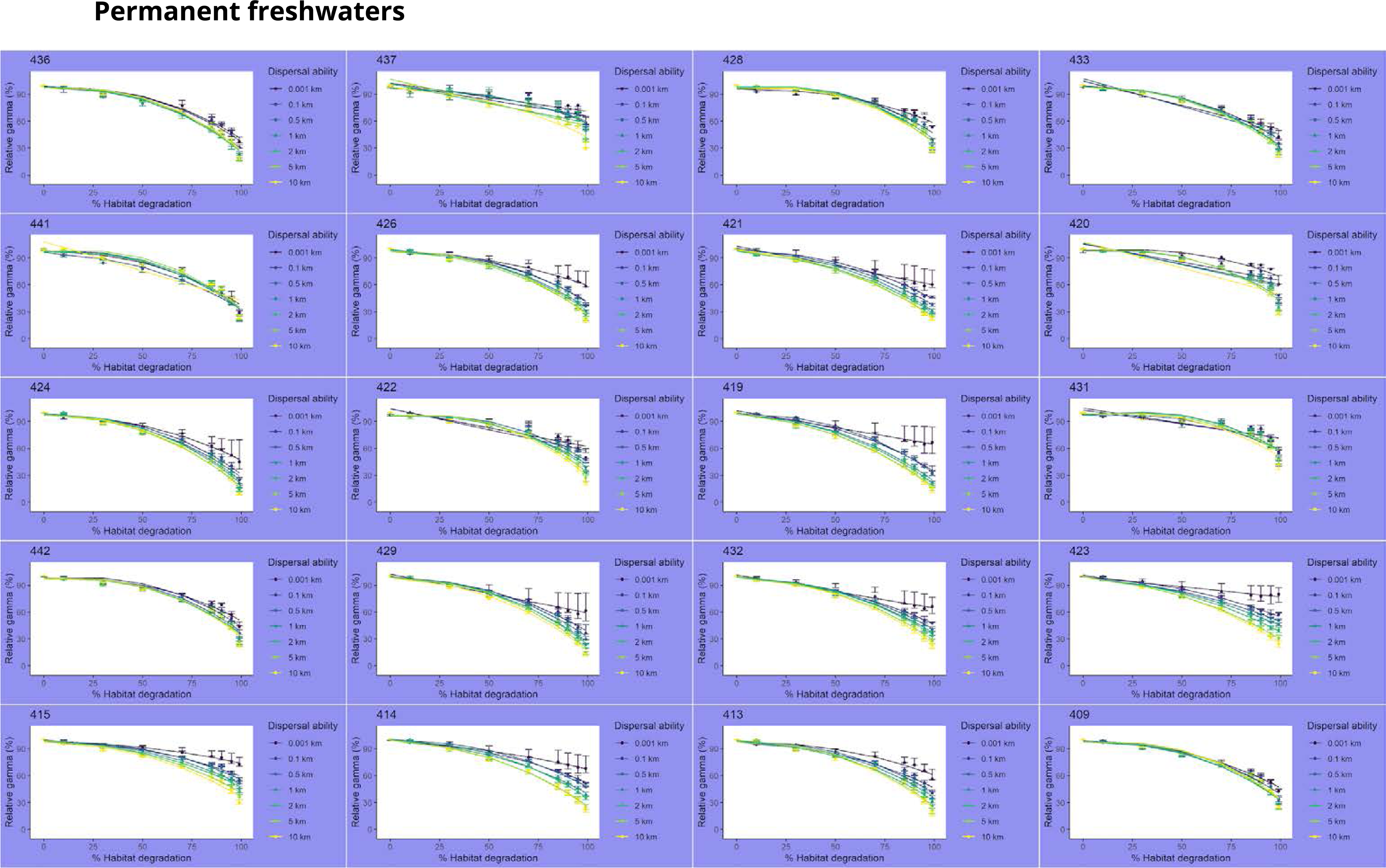
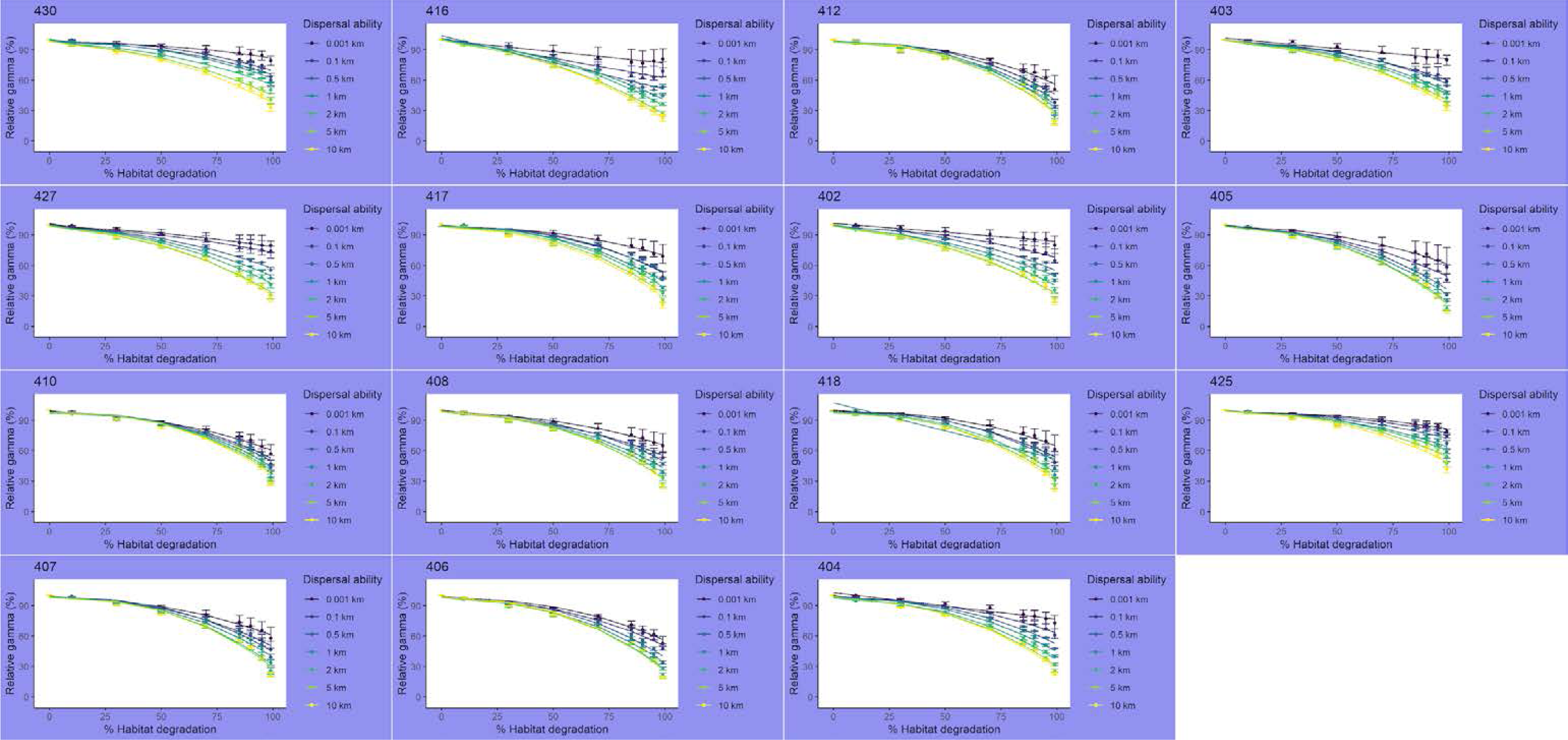
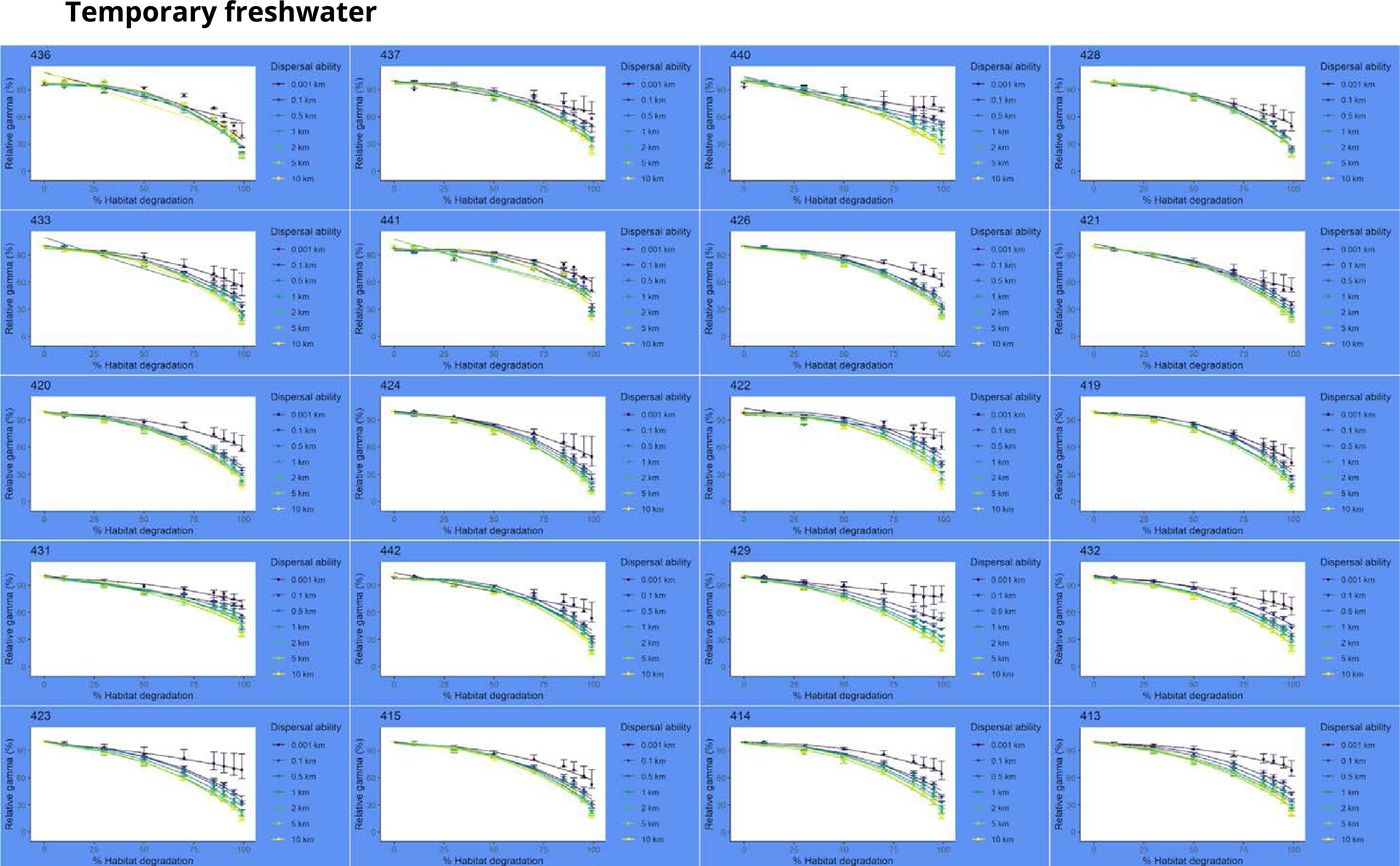
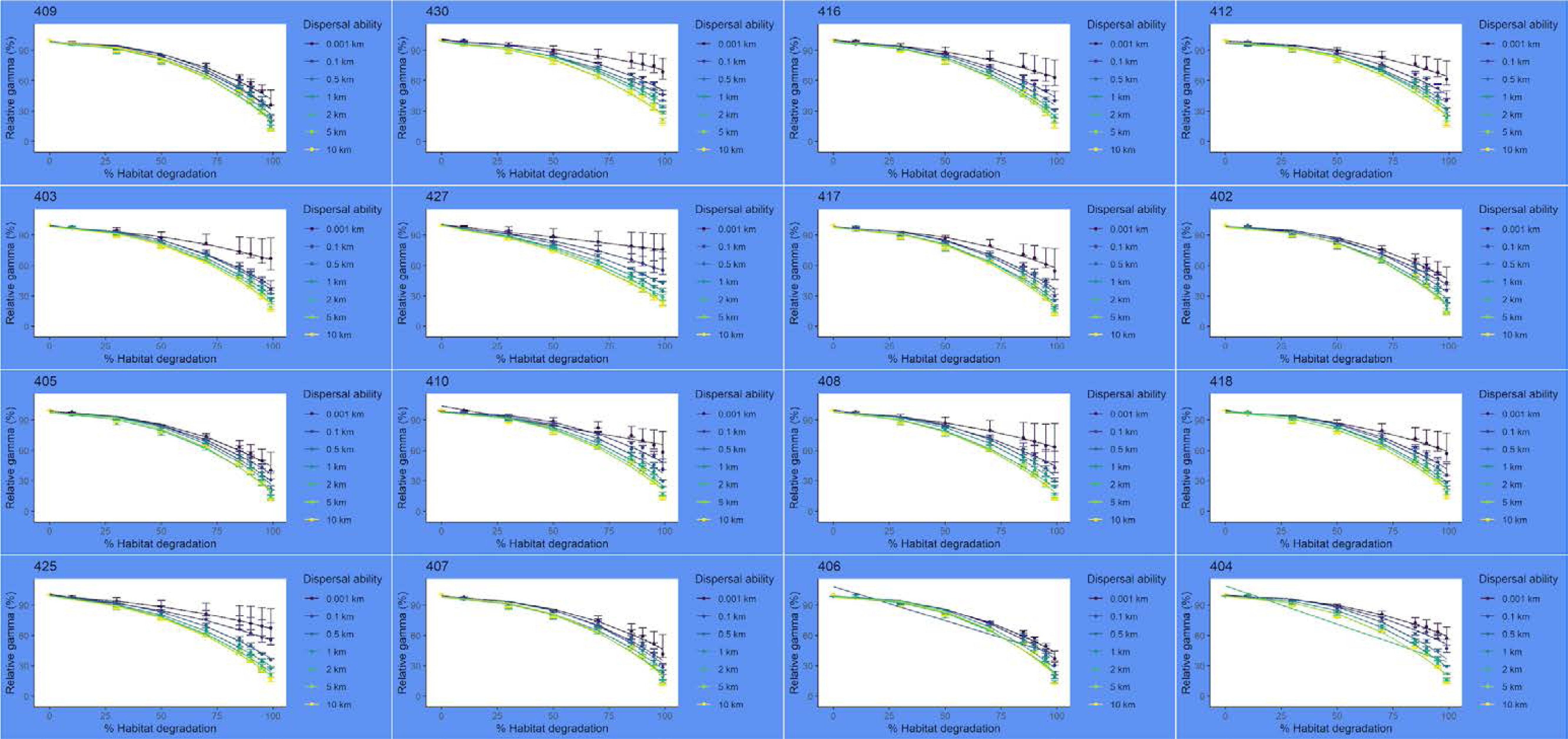
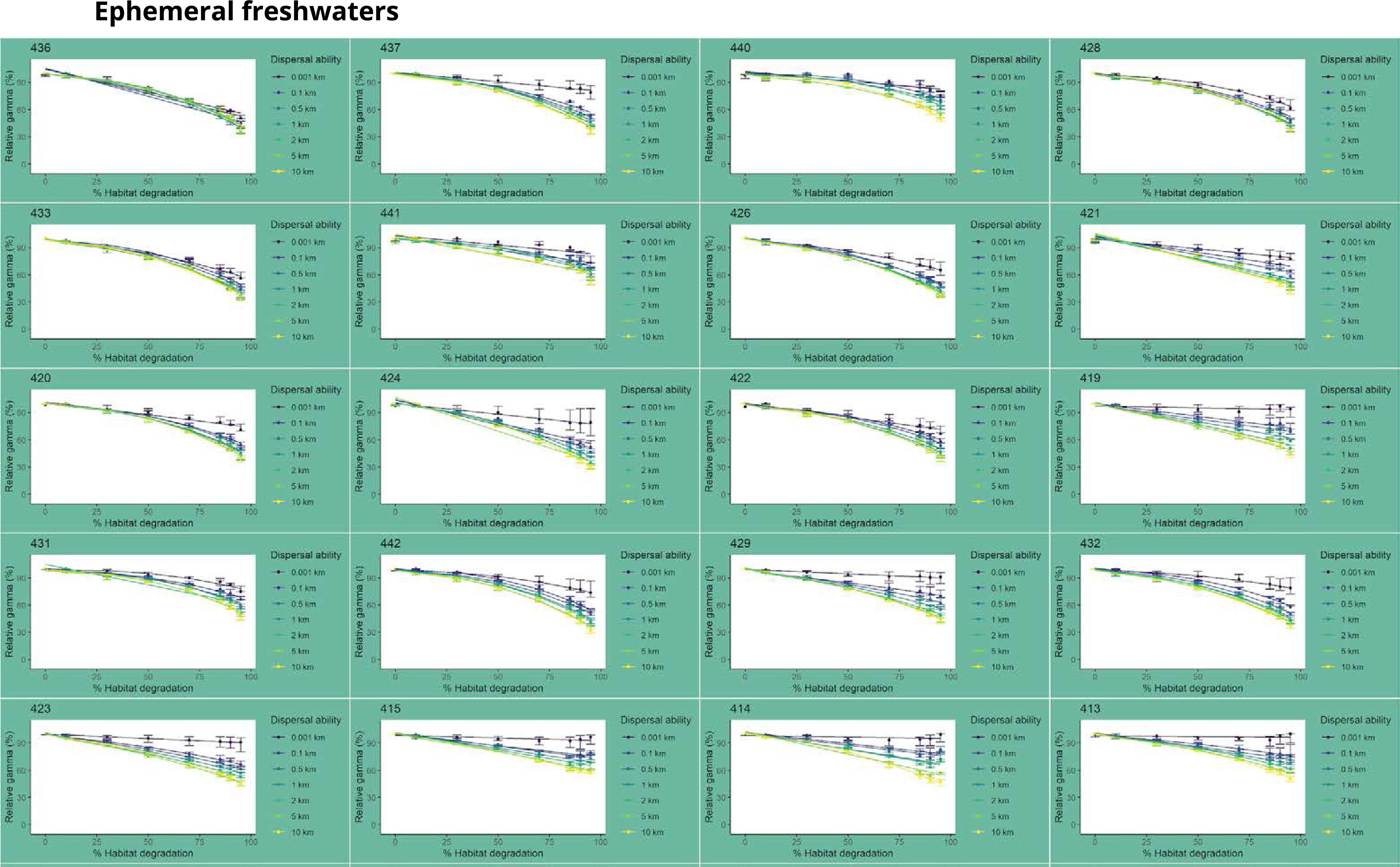
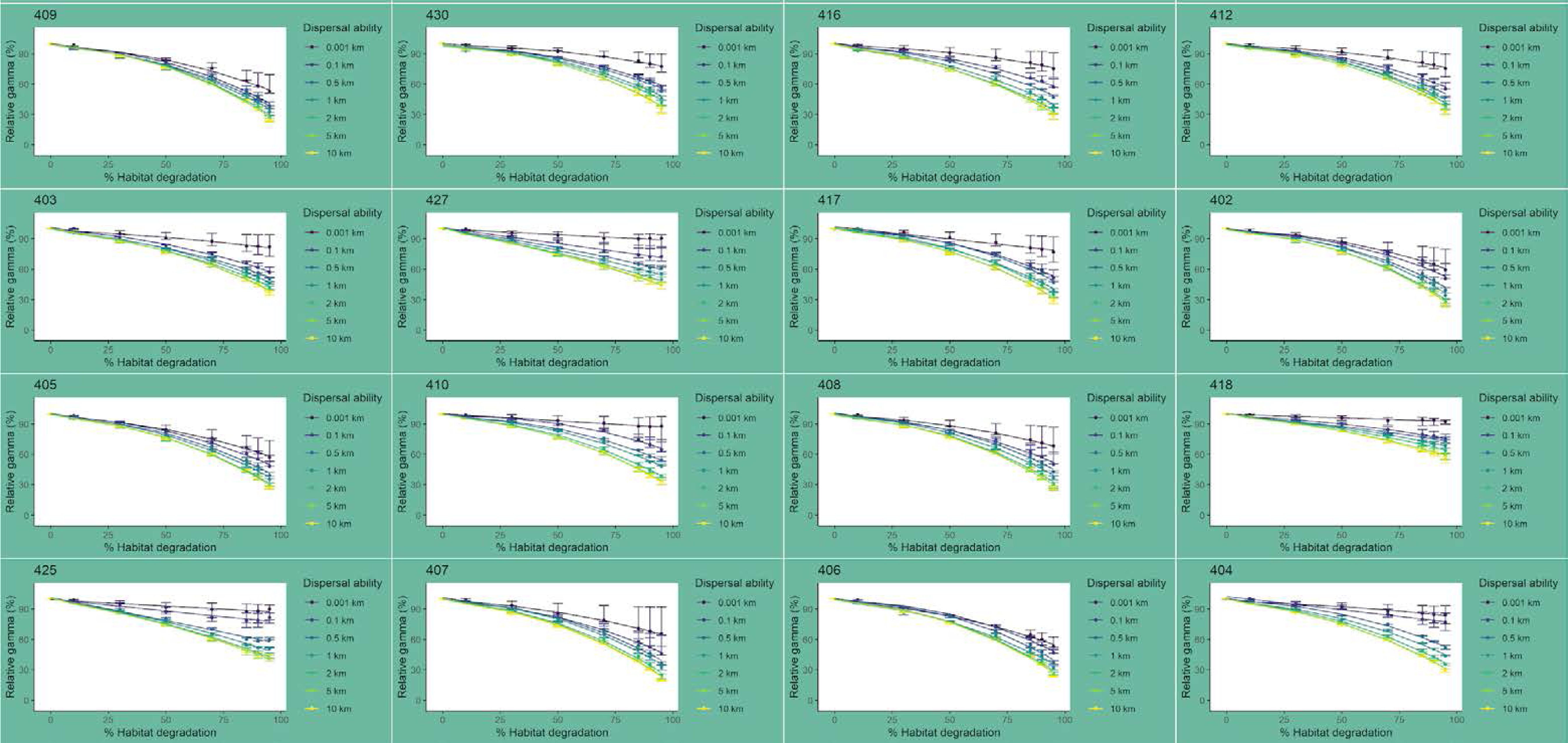
Gamma diversity results for each ecoregion and their corresponding diversity decay curves. From yellow to purple, different dispersal abilities are indicated for Total (white background), permanent (purple background), temporary (blue background) and ephemeral freshwaters (green background).

**Supplementary material 5:**
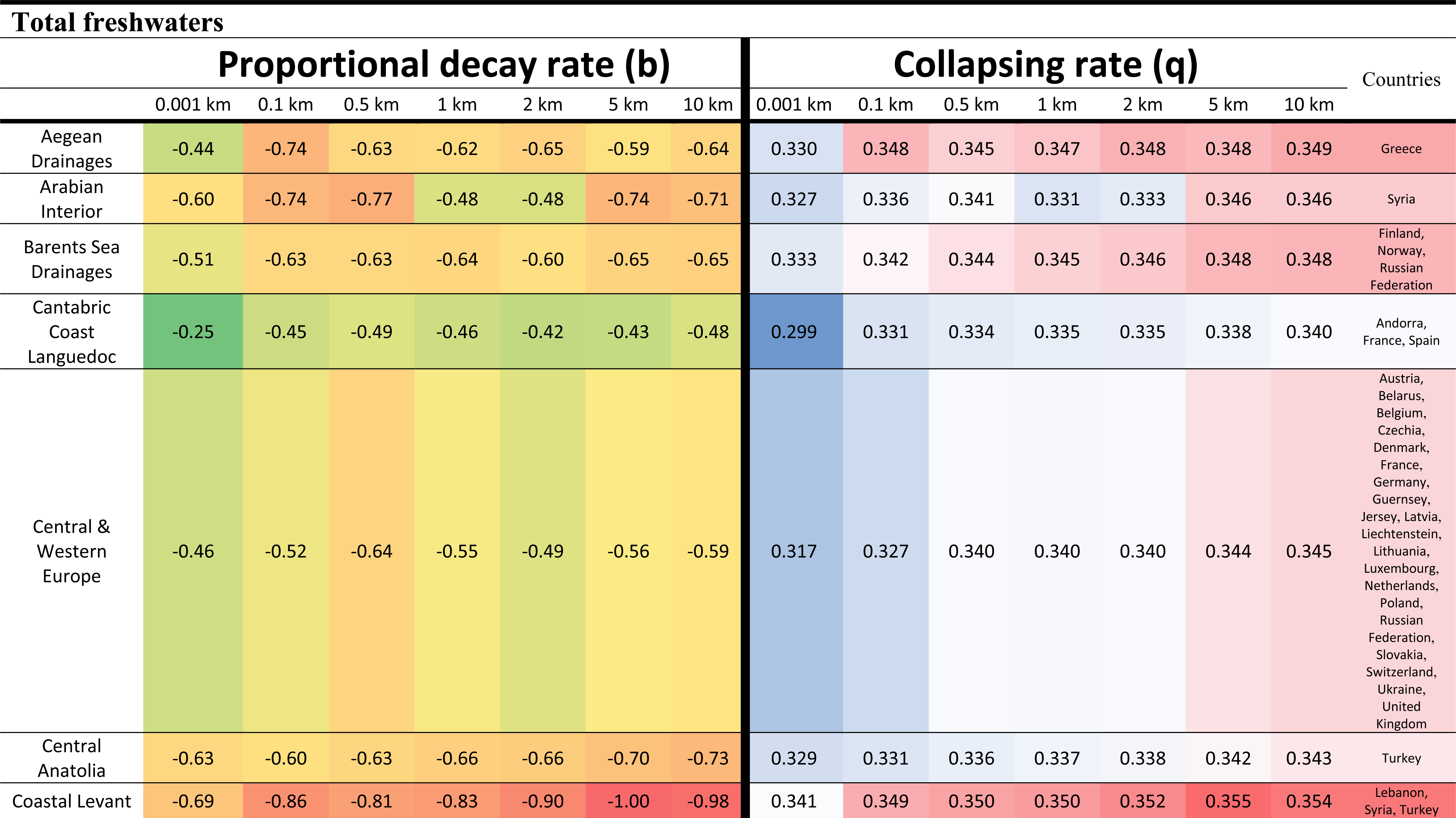
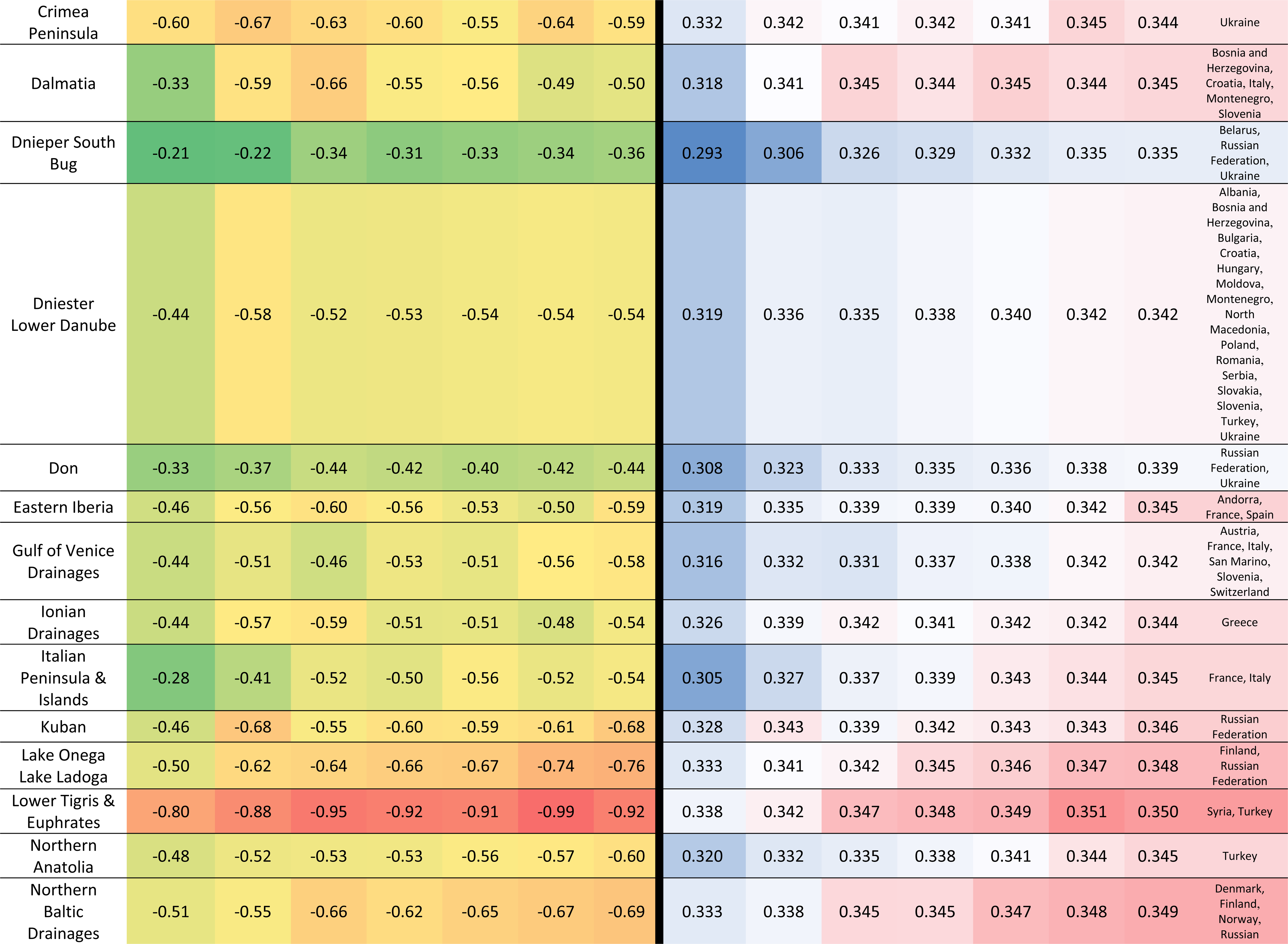
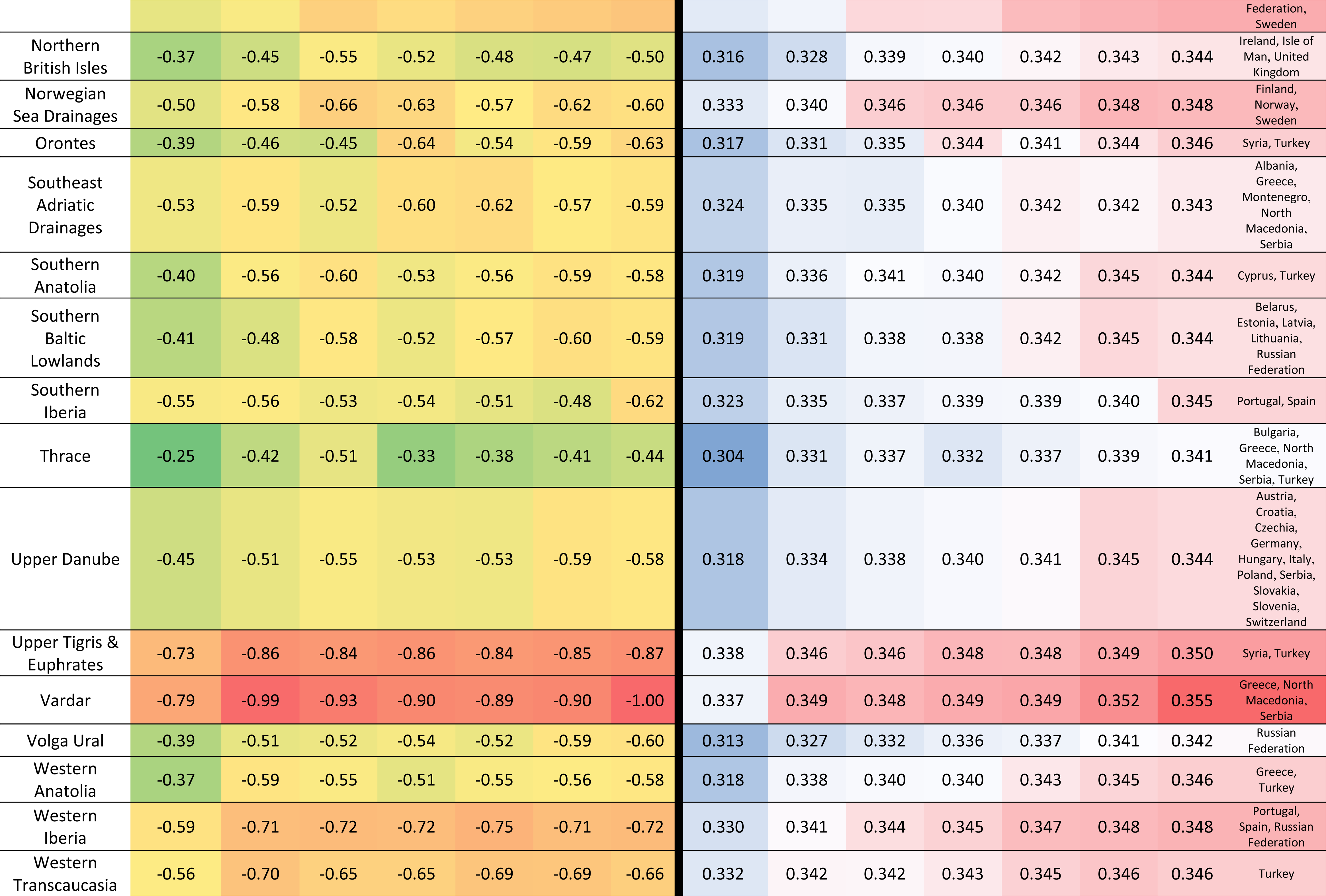
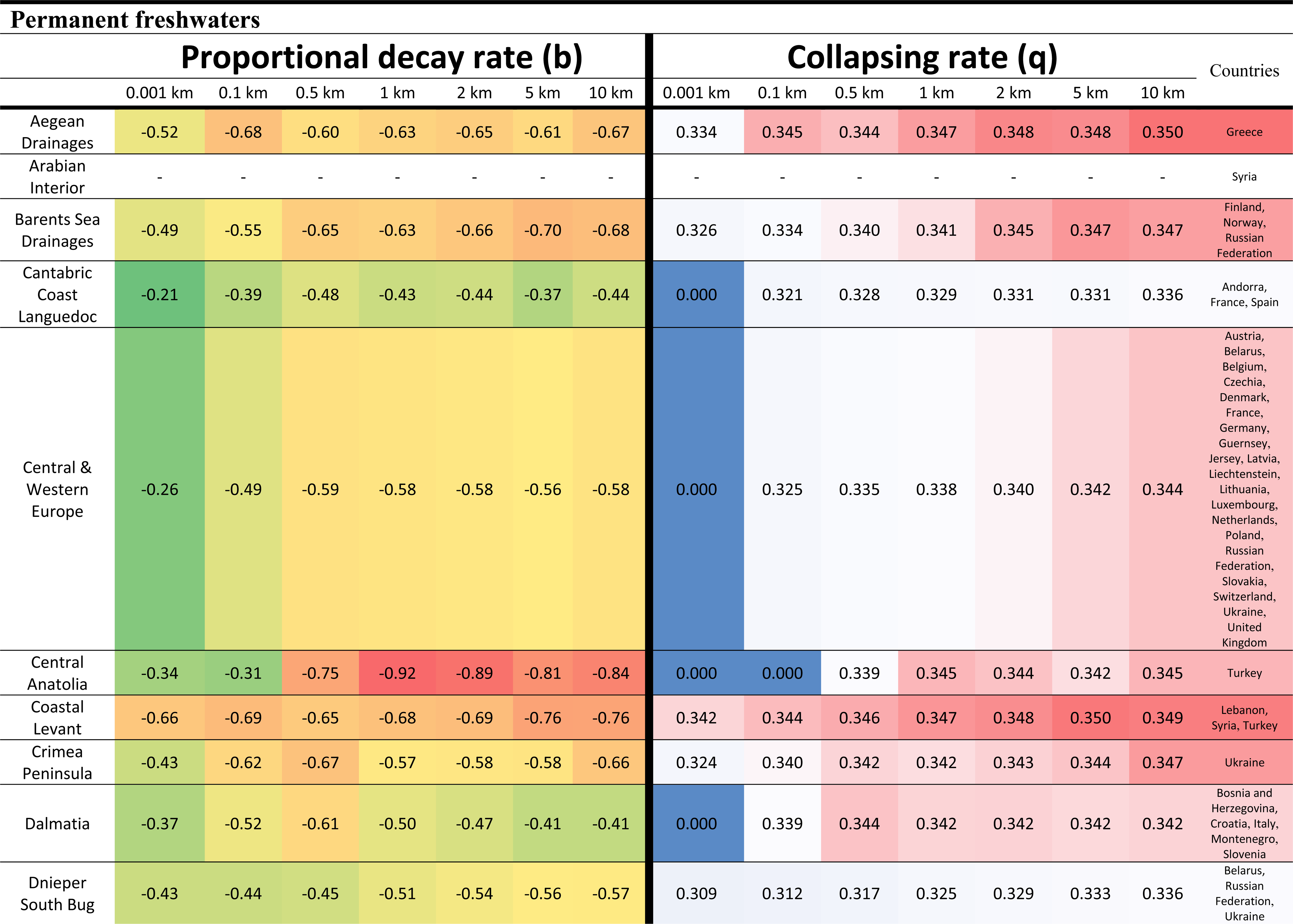
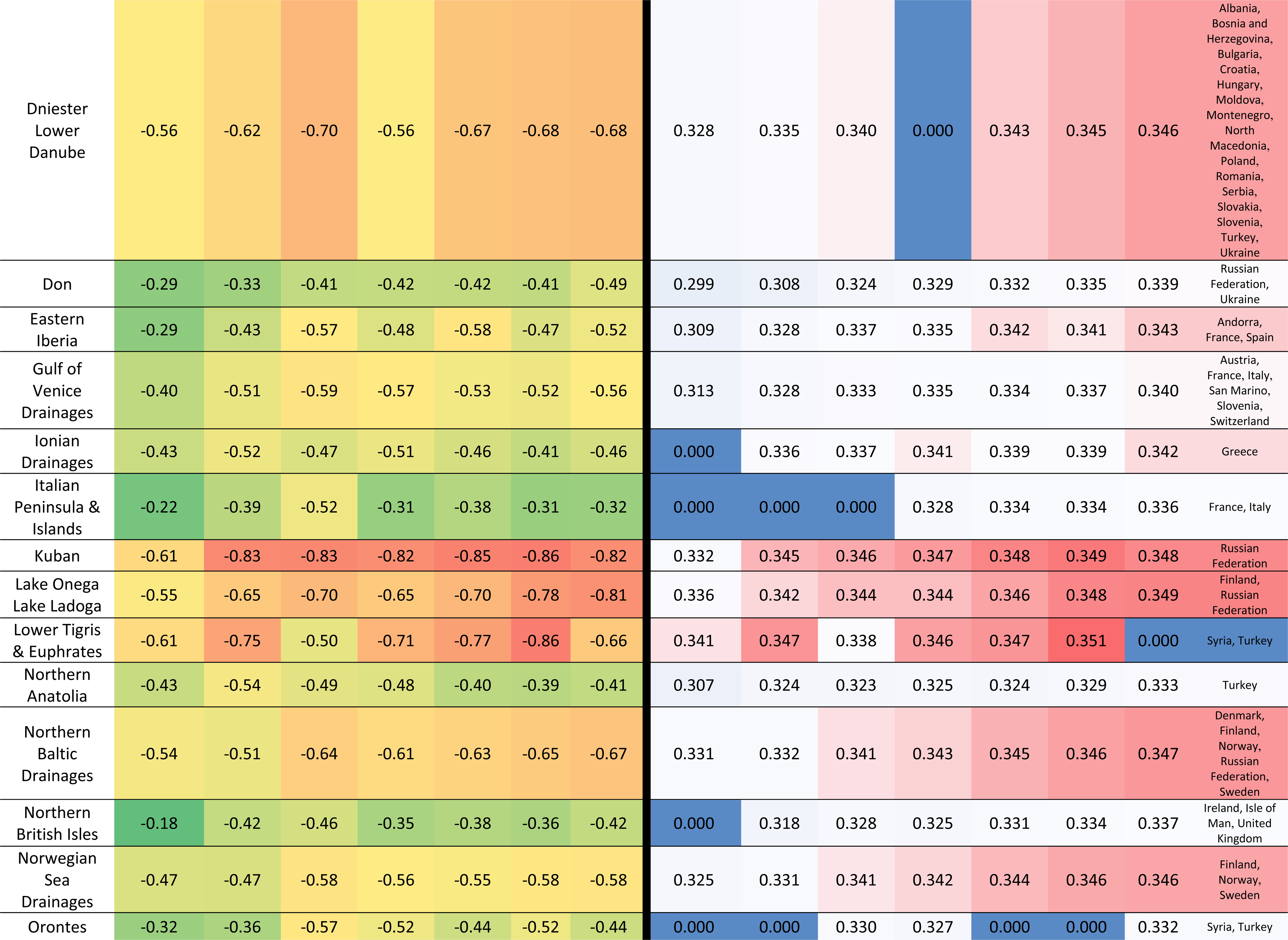
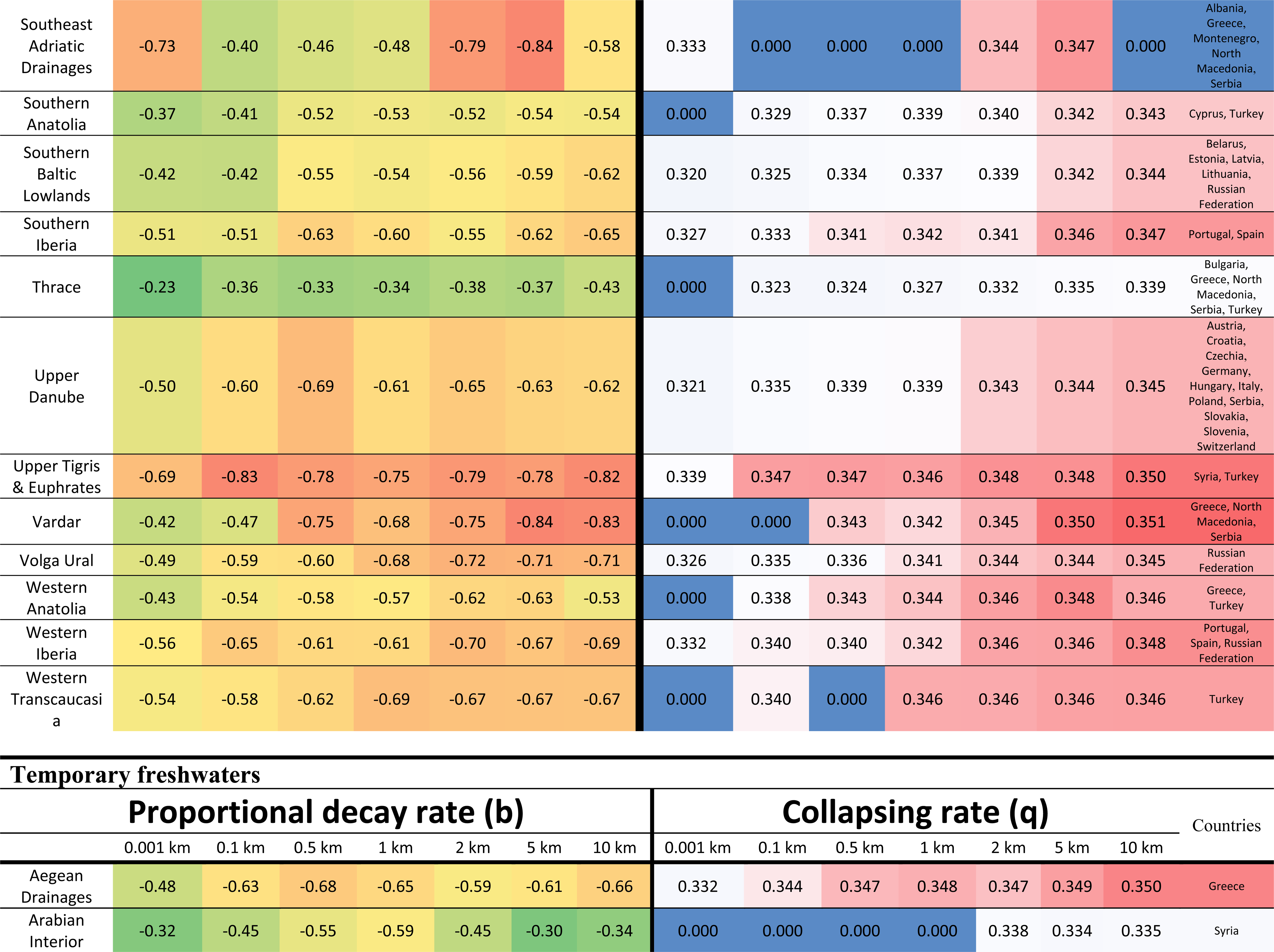
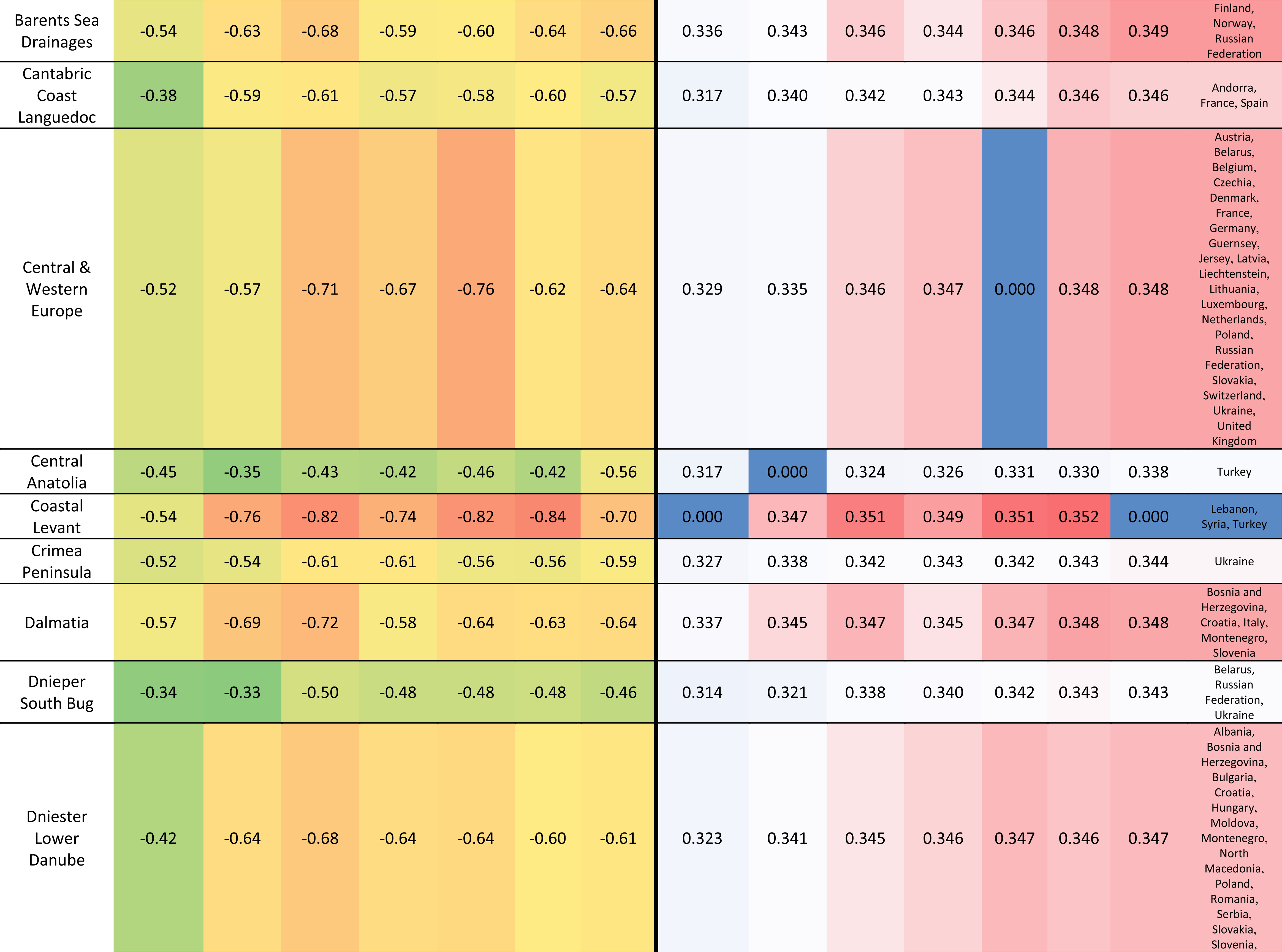
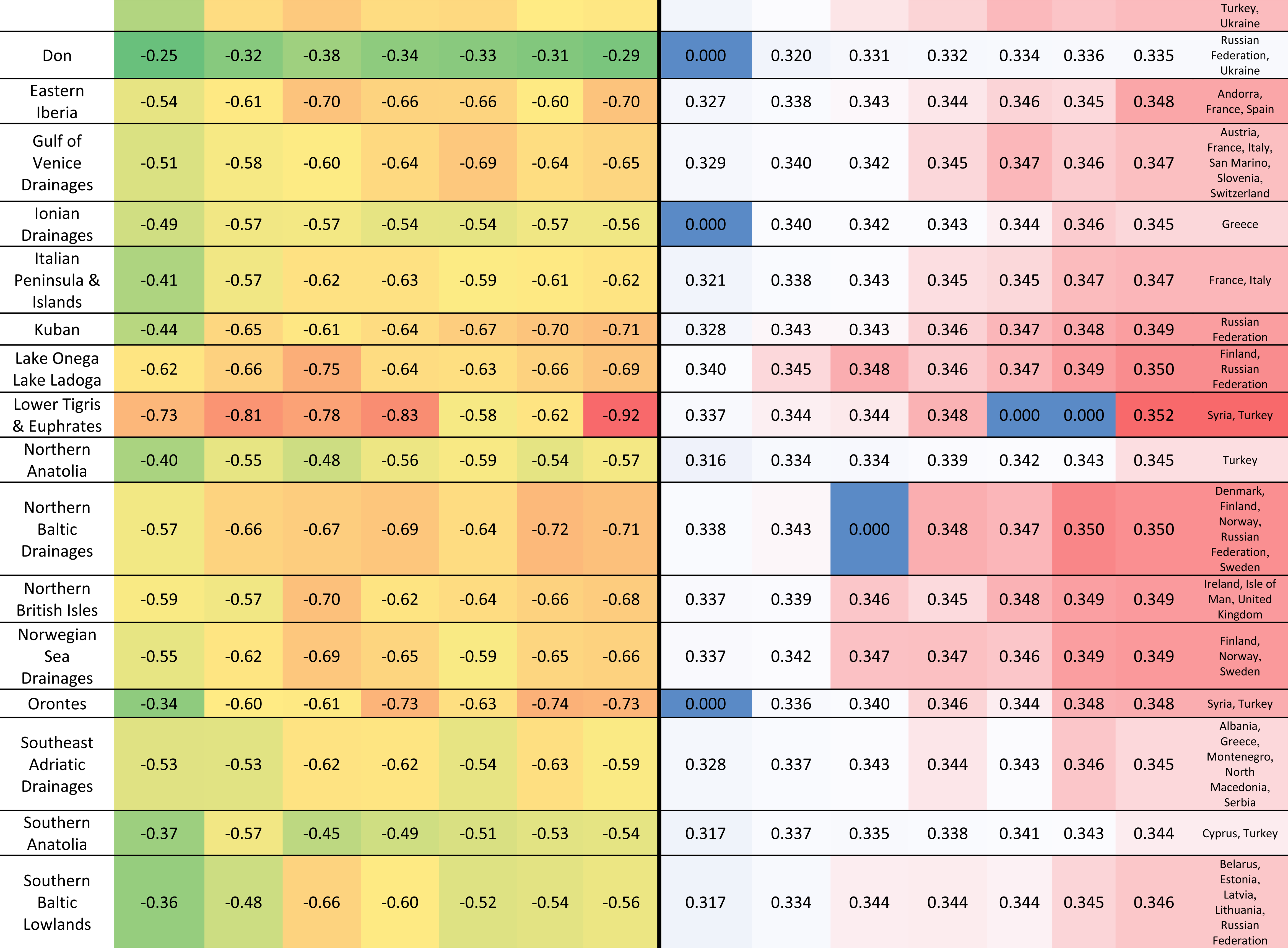
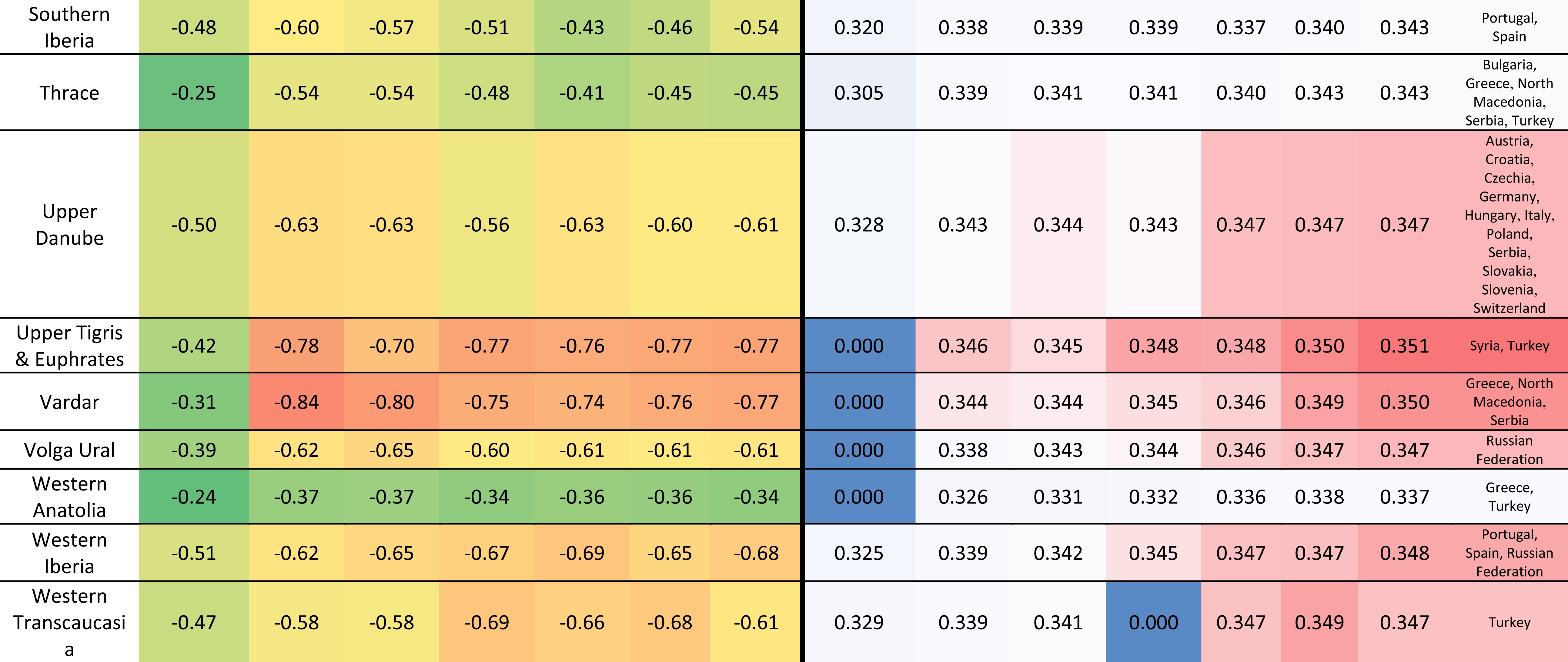
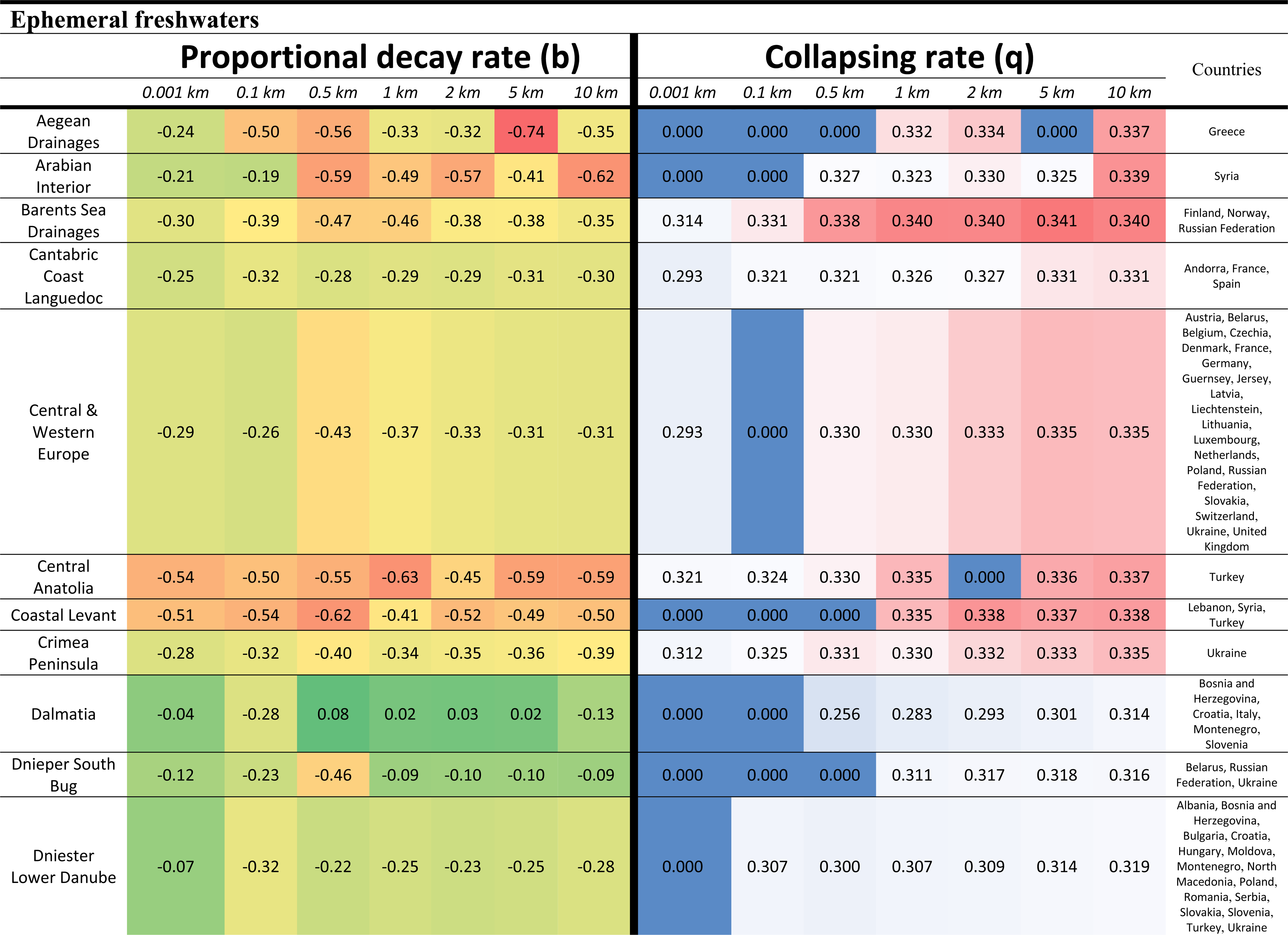
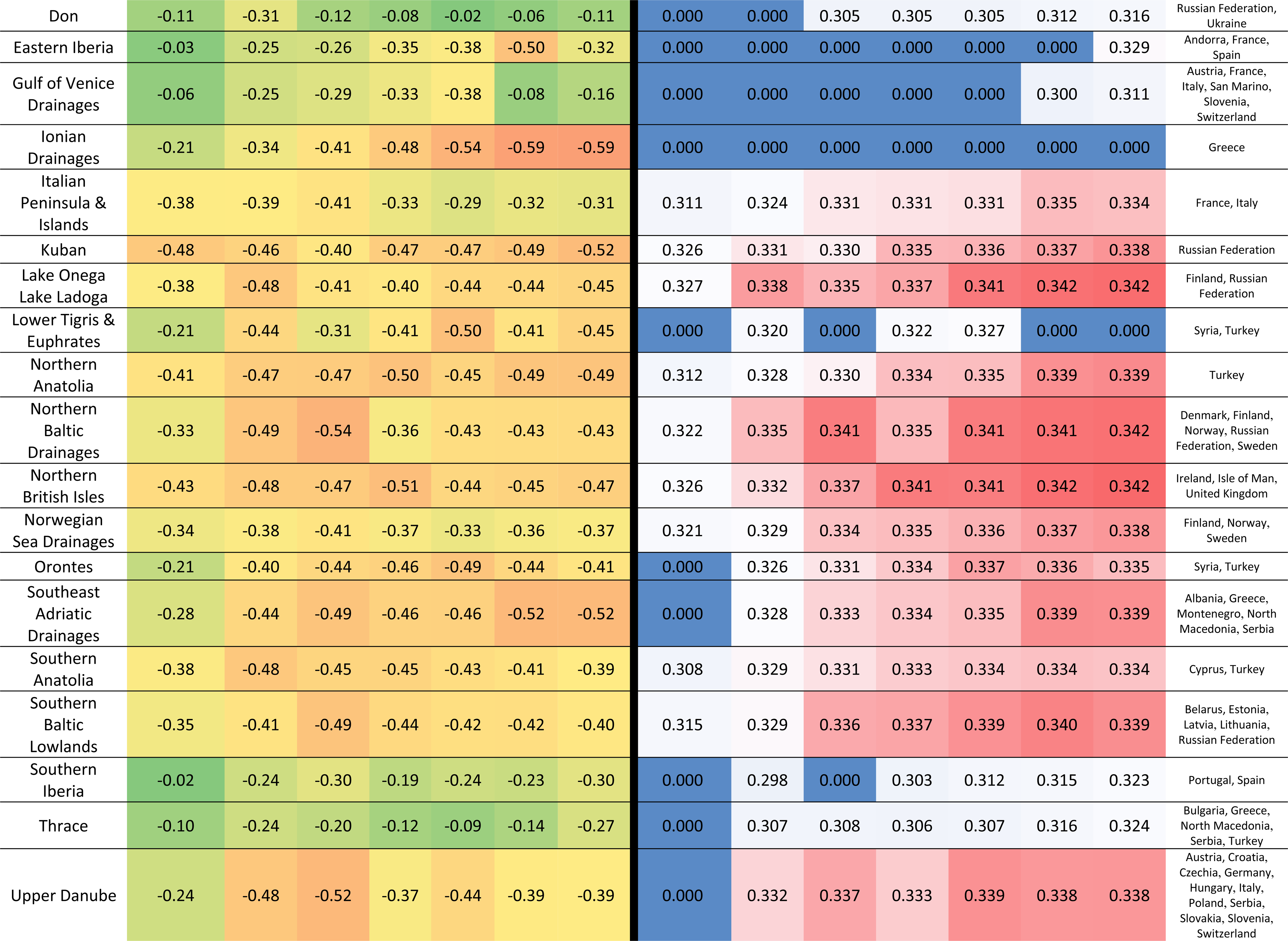
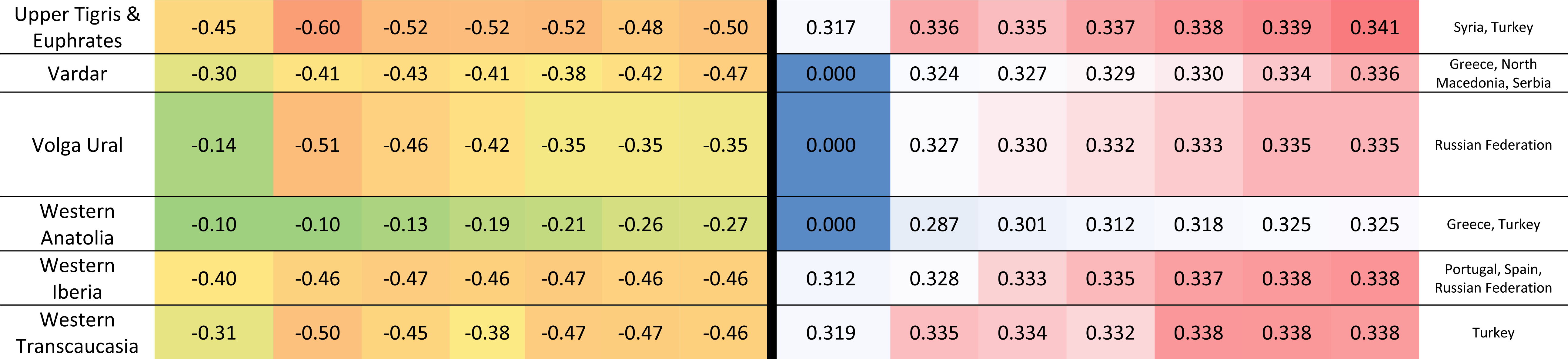
Diversity decay values for each ecoregion and dispersal ability for total, permanent, temporary, and ephemeral freshwaters. The countries where the ecoregion belongs are indicated. Colours are visual guides to indicate the range of proportional decay rate values (smaller in green and higher in red) and for collapsing rate (smaller in blue and higher in red). The proportional decay rate (b) describes the velocity at which species are lost per percentual unit of habitat lost, with values closer to ™1 meaning a loss of 1 species per each 1% of habitat lost and values closer to 0 meaning that no species is lost with landscape degradation. The collapsing rate (q) describes the acceleration of species loss per each percentual unit of habitat lost, with high values (around 0.36) implying a rapid collapse of diversity with habitat loss and values closer to 0 implying no acceleration at all or in other words, the diversity decay is constant and describes only a straight line with slope equal to the proportional decay rate (b).

**Supplementary material 6:**
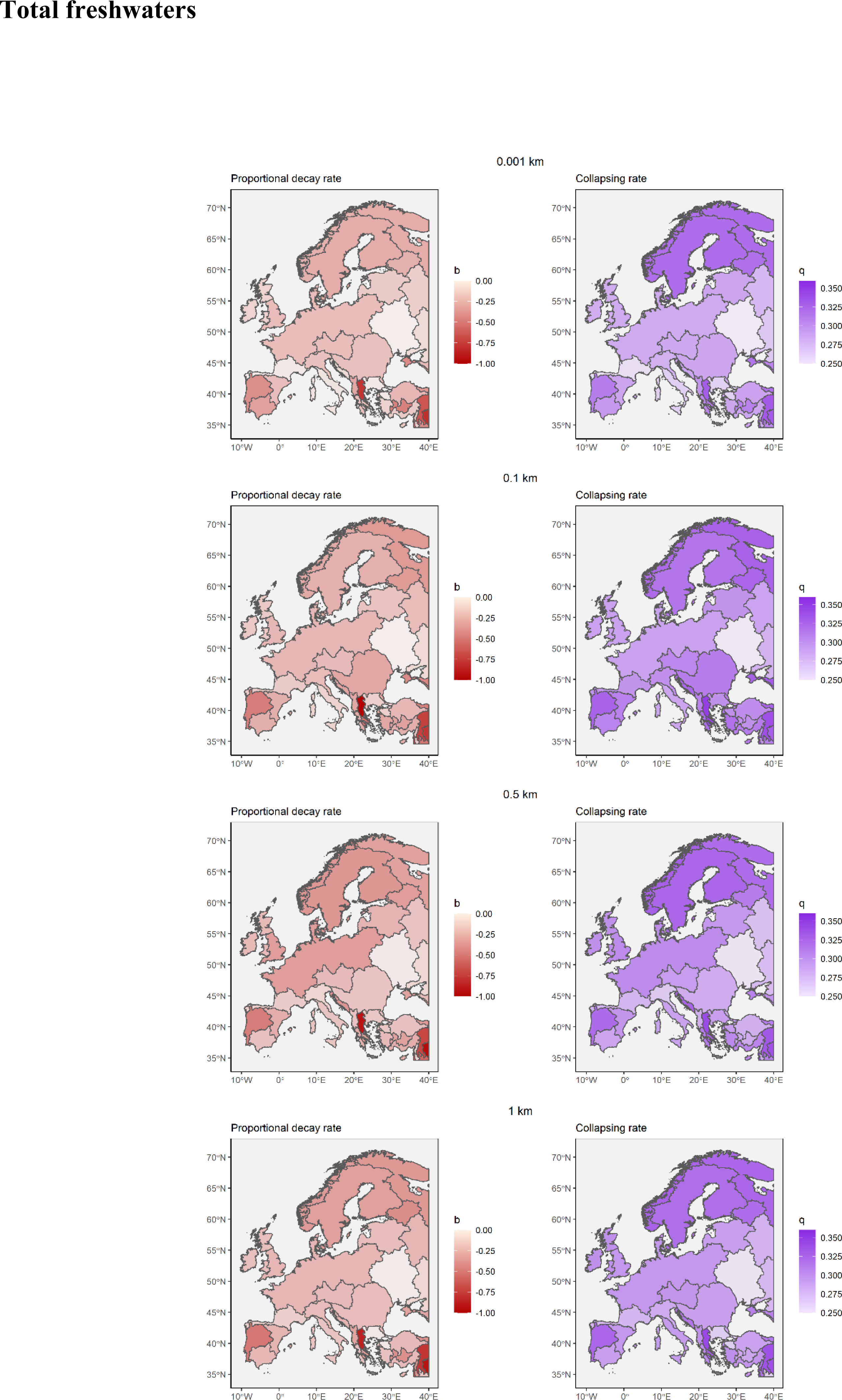
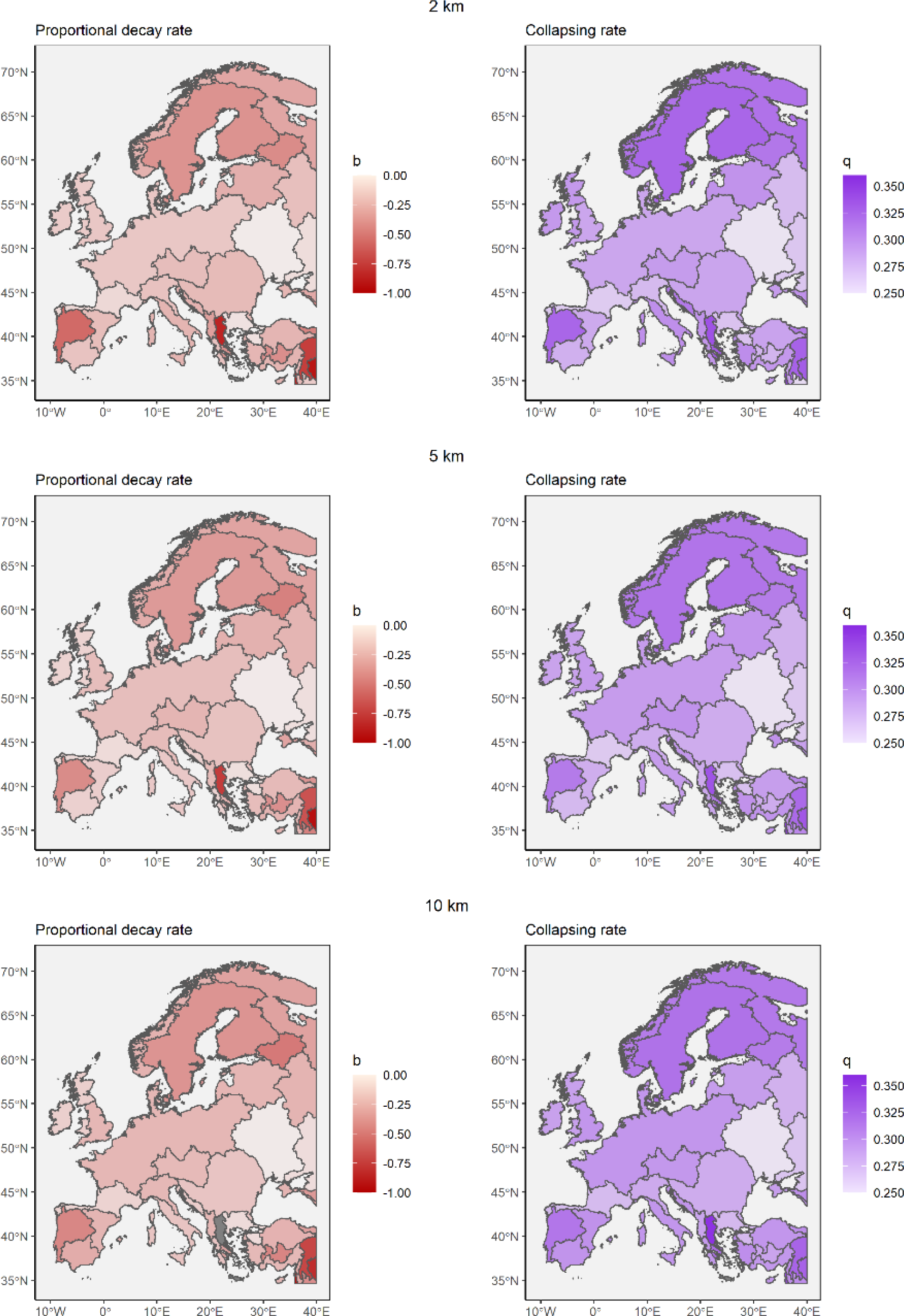
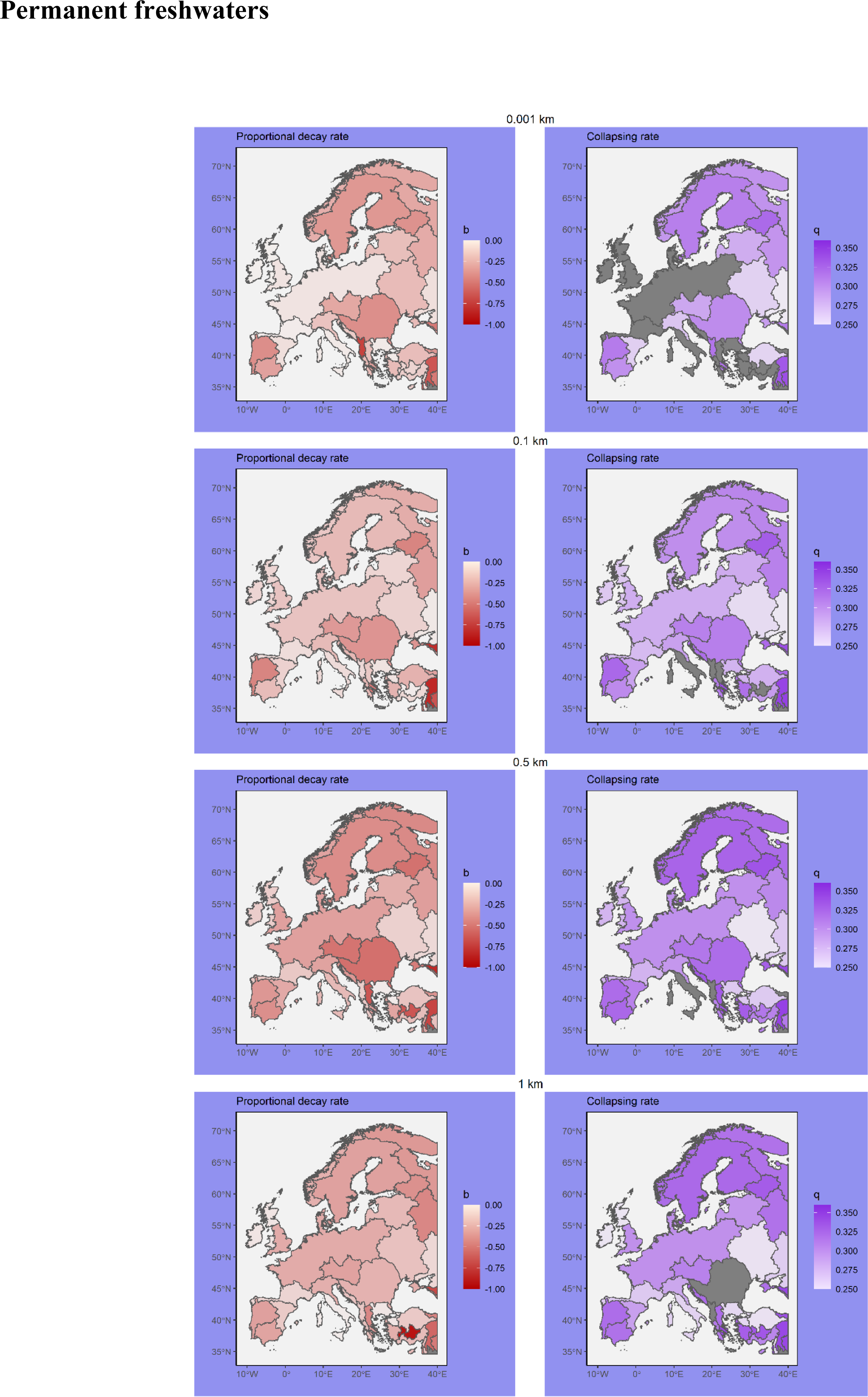
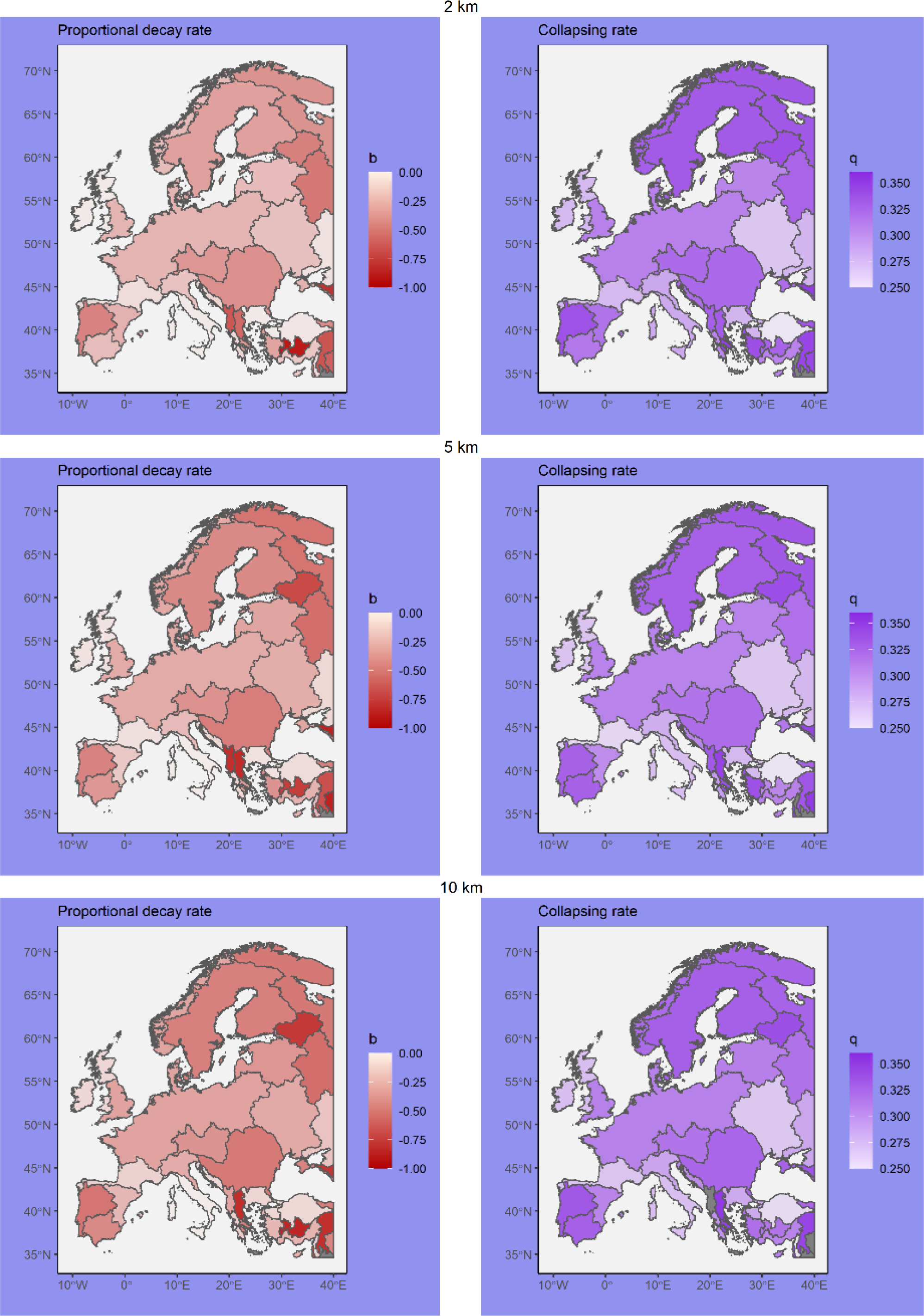
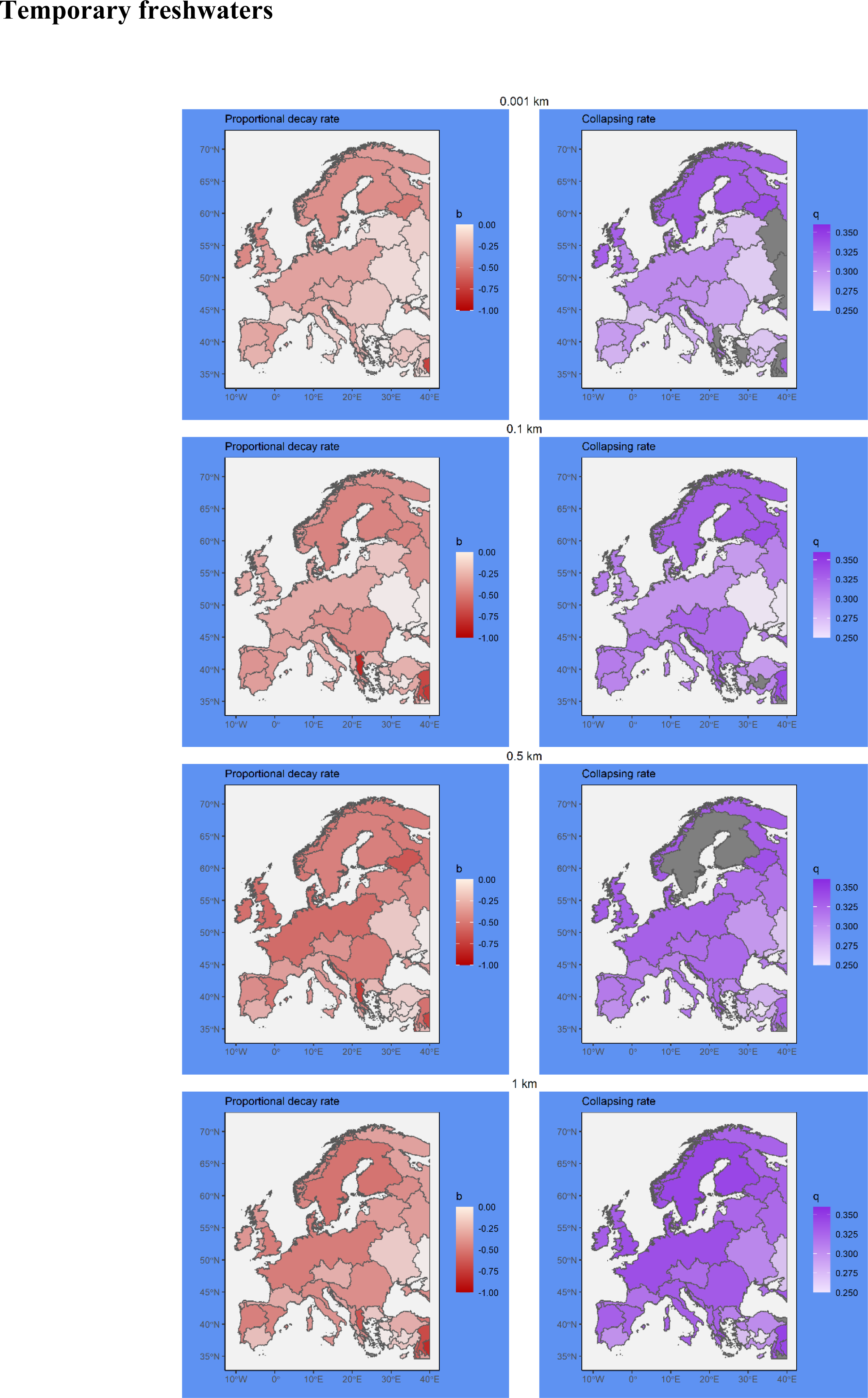
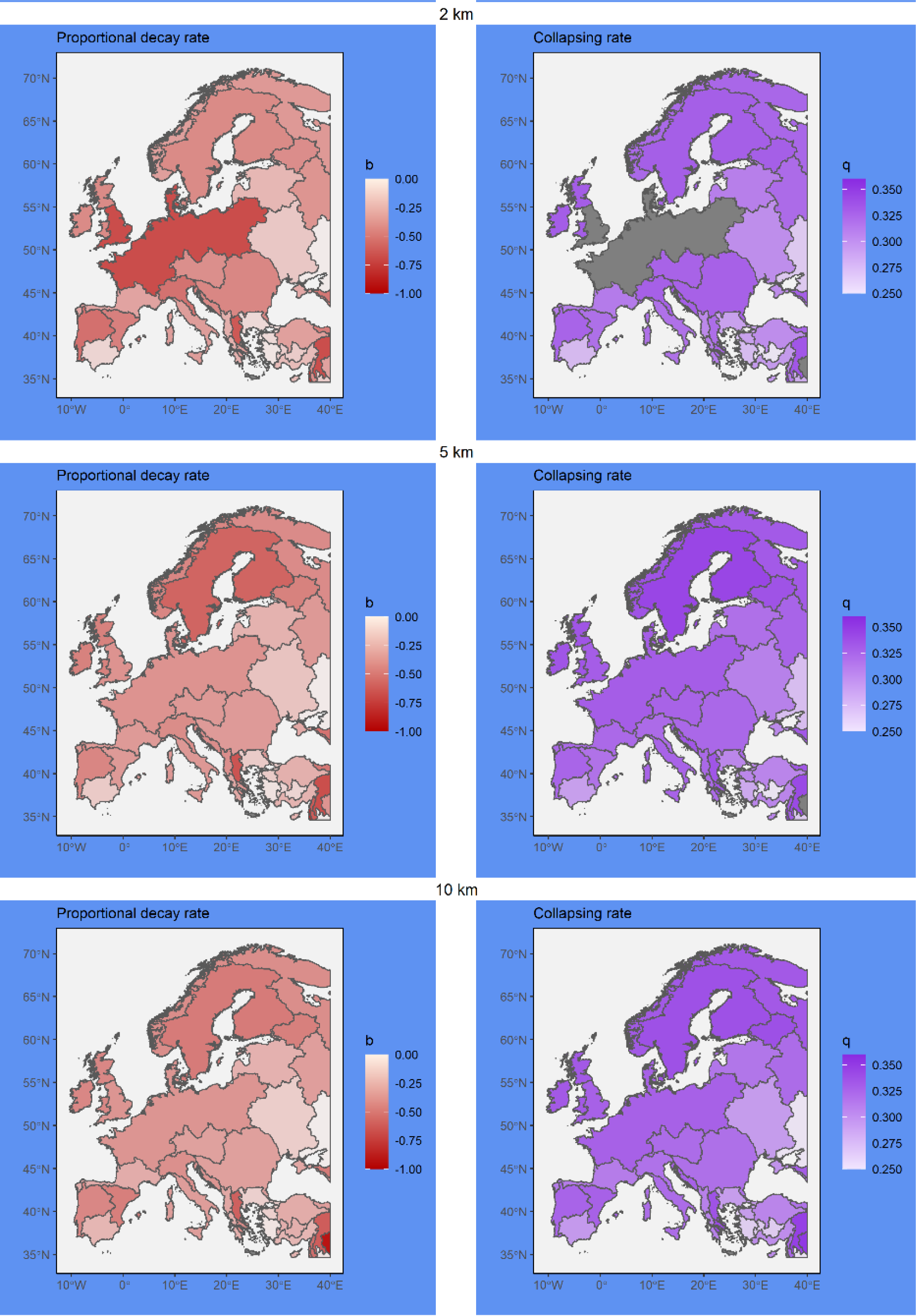
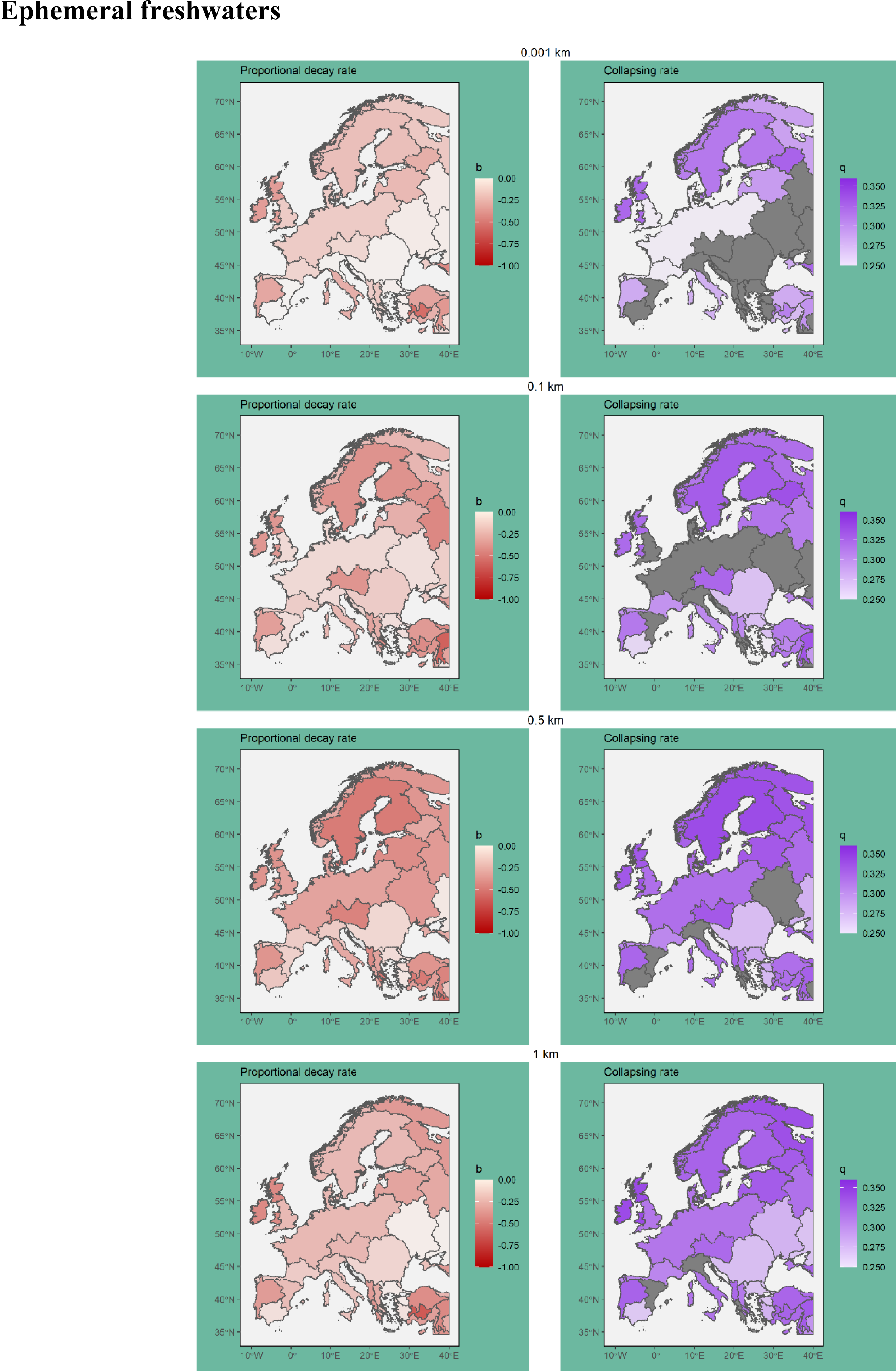
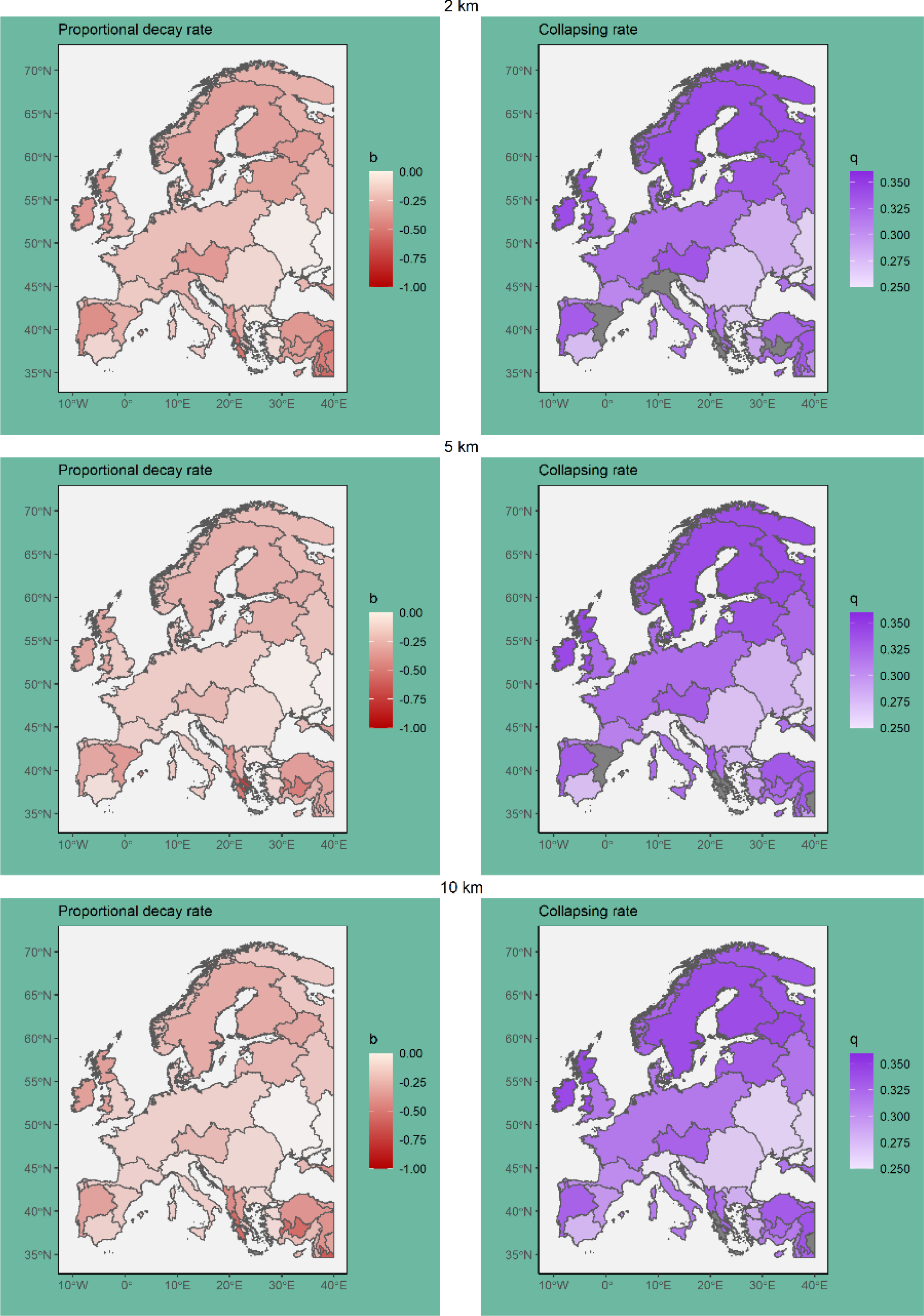
European diversity decay maps for each ecoregion and for each type of freshwaters: total, permanent, temporary, and ephemeral (white, purple, blue, and green background respectively). All maps are scaled to be comparable among them. The proportional decay rate (b) describes the velocity at which species are lost per percentual unit of habitat lost, with values closer to ™1 meaning a loss of 1 species per each 1% of habitat lost and values closer to 0 meaning that no species is lost with landscape degradation. The collapsing rate (q) describes the acceleration of species loss per each percentual unit of habitat lost, with high values (around 0.36) implying a rapid collapse of diversity with habitat loss and values closer to 0 implying no acceleration at all. In other words, when equal to 0 the diversity decay is constant and describes only a straight line with slope equal to the proportional decay rate (b).

**Supplementary material 7:**
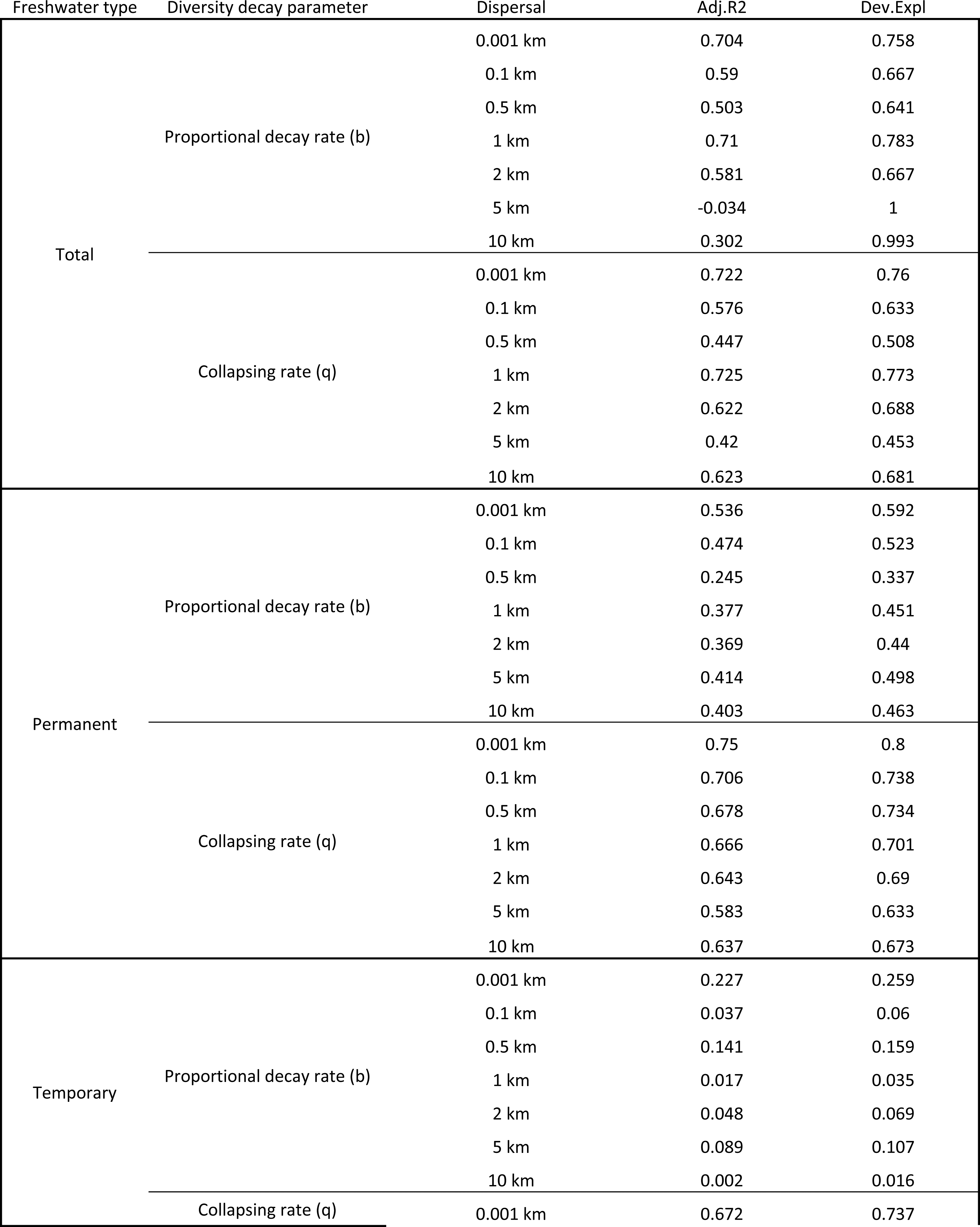
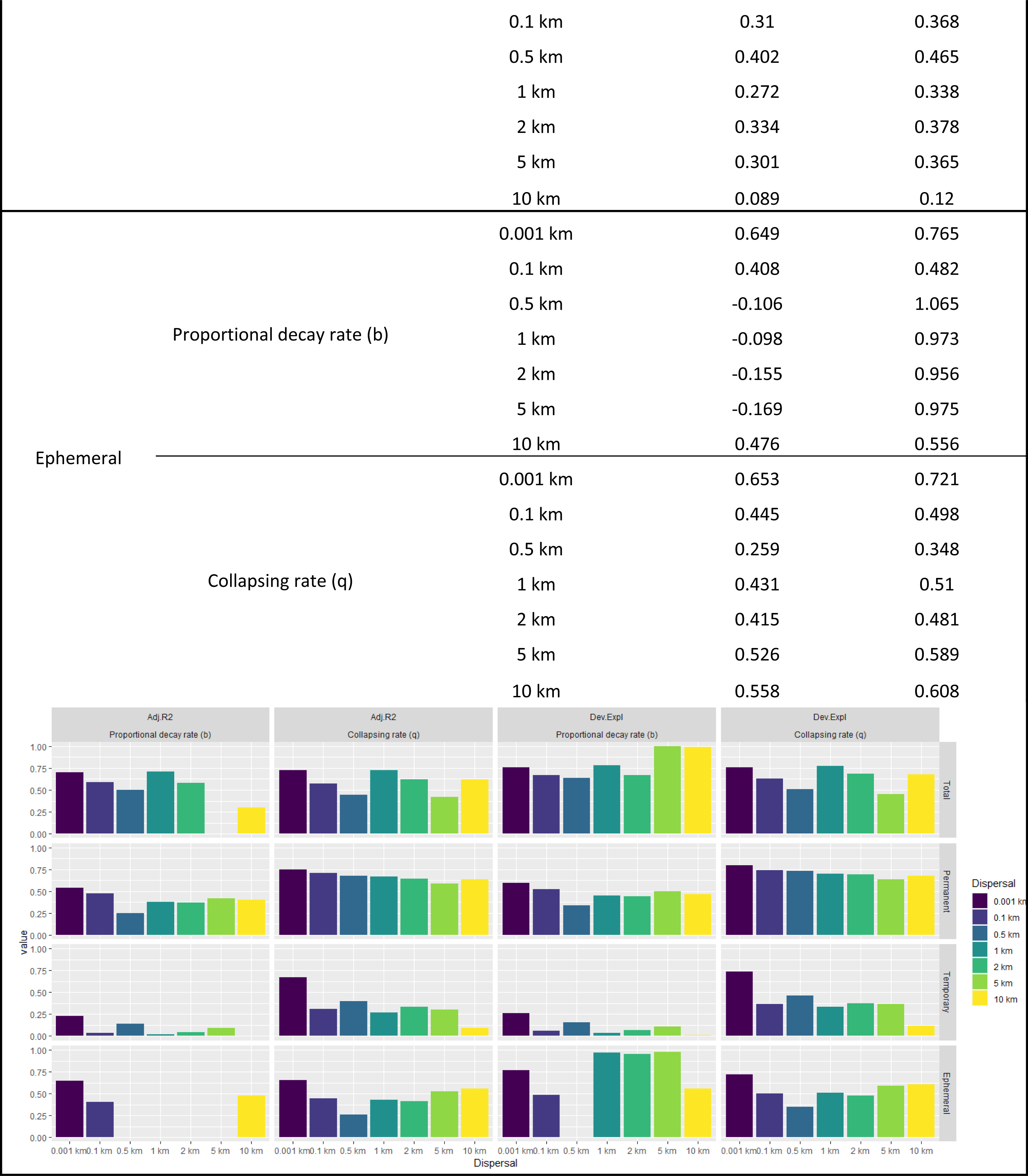
Gam model adjusted squared R and explained deviation for each type of freshwater (Total, Permanent, Temporary and Ephemera), Diversity decay parameter (Proportional decay rate and Collapsing rate) and the 7 dispersal distances (0.001 km, 0.1 km, 0.5 km, 1 km, 2 km, 5 km, 10 km). A summarizing barplot plot with the same information is also included to facilitate visual comparisons betewen models.

**Supplementary material 8:**
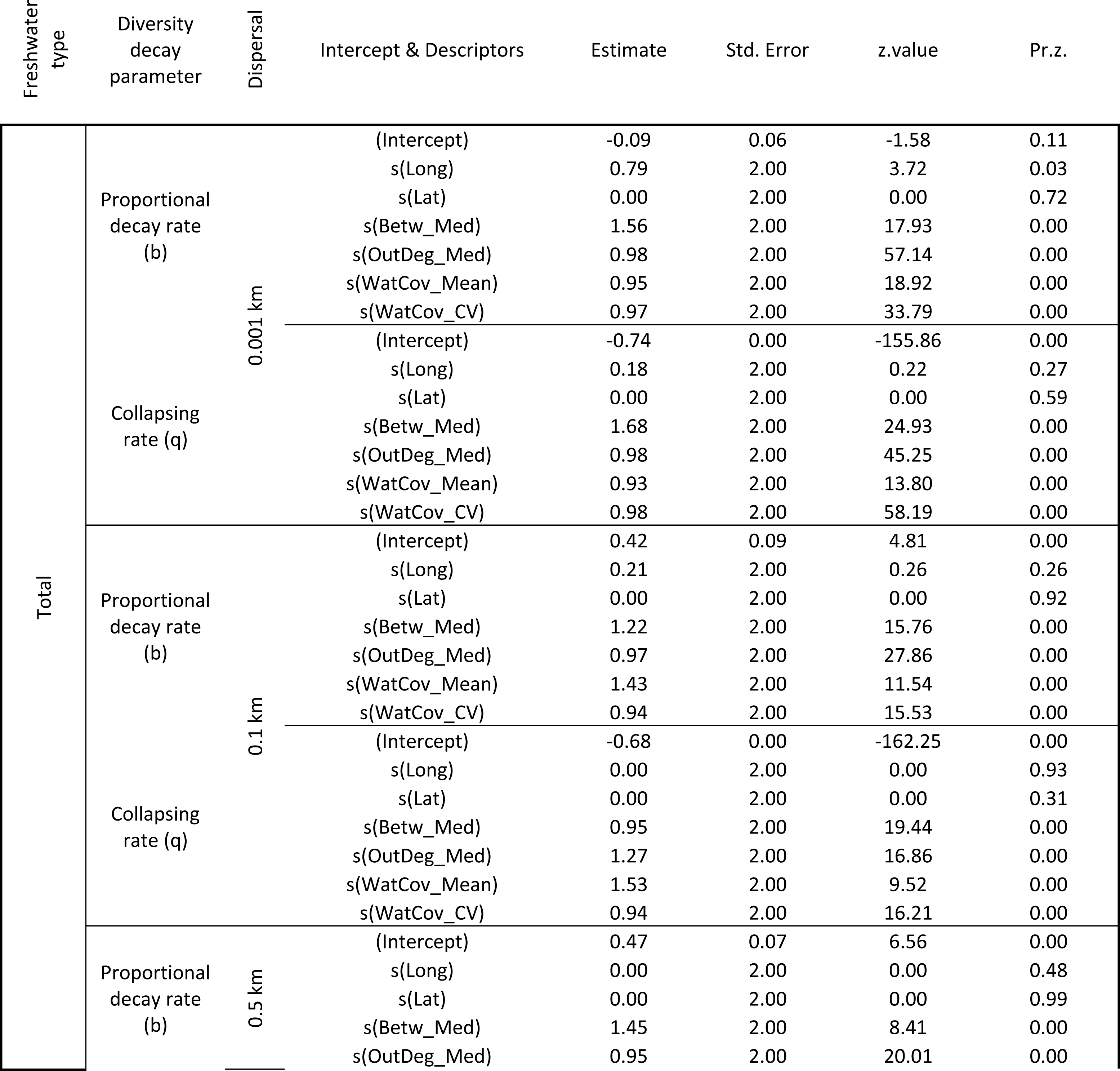
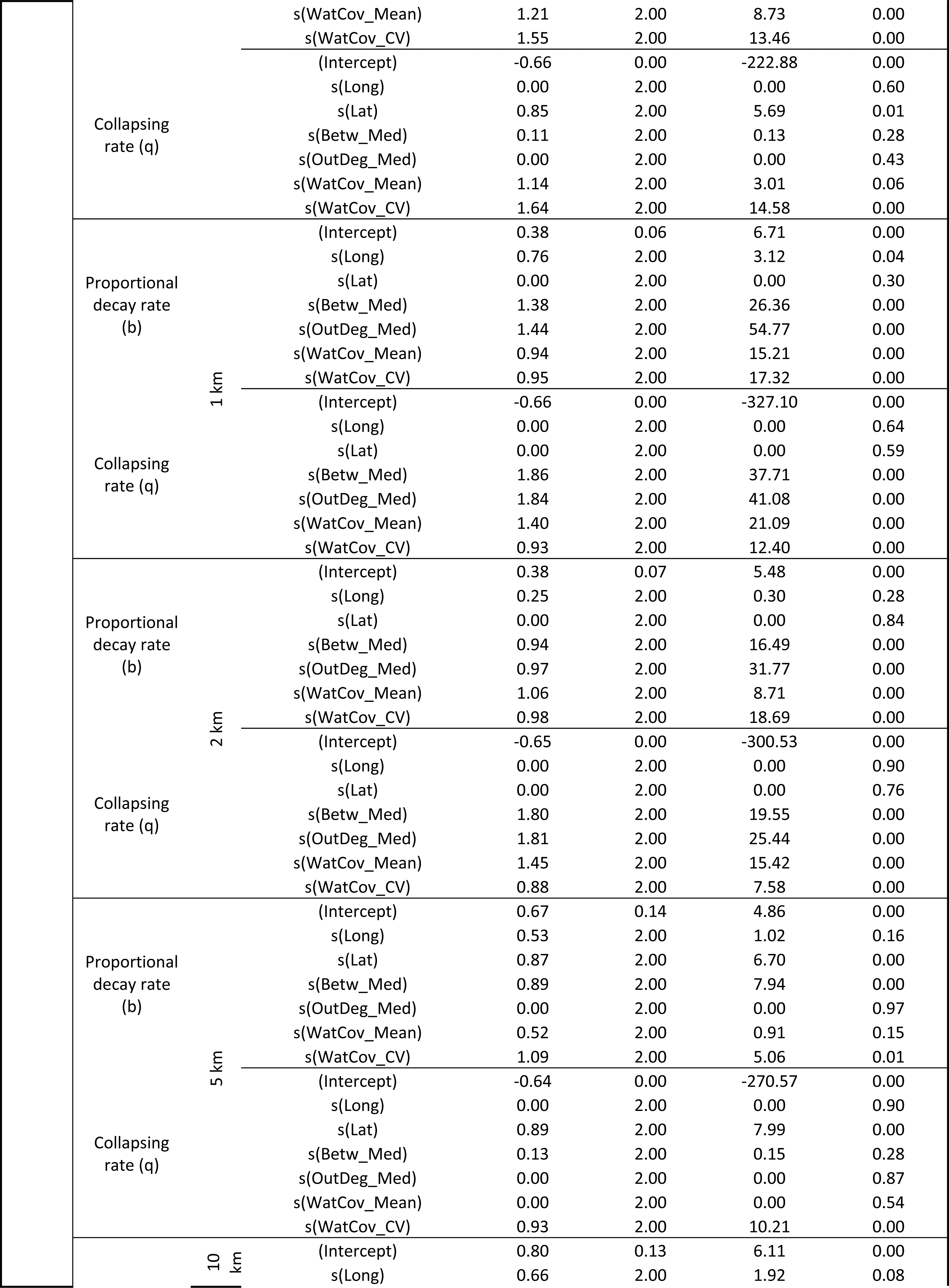
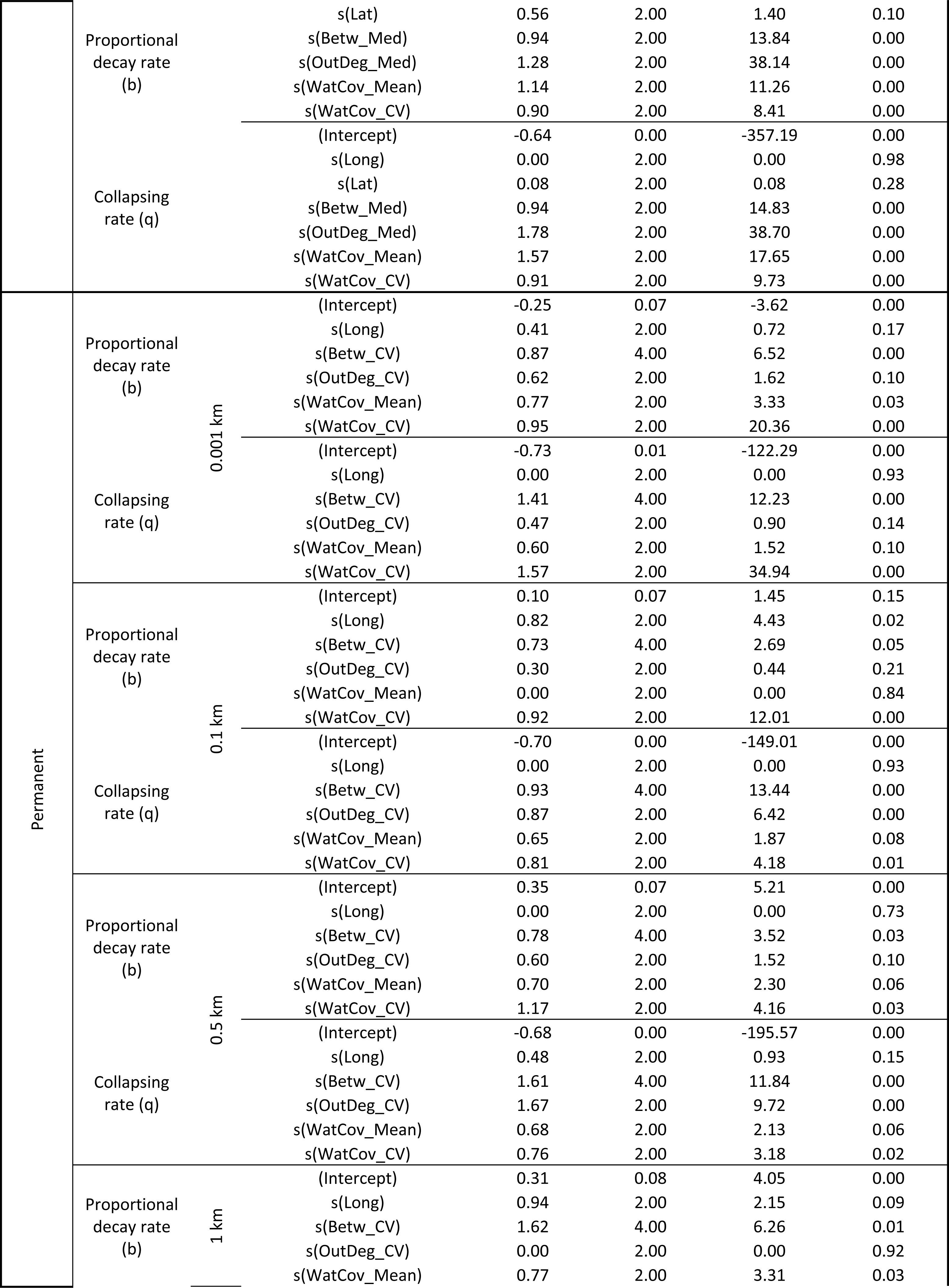
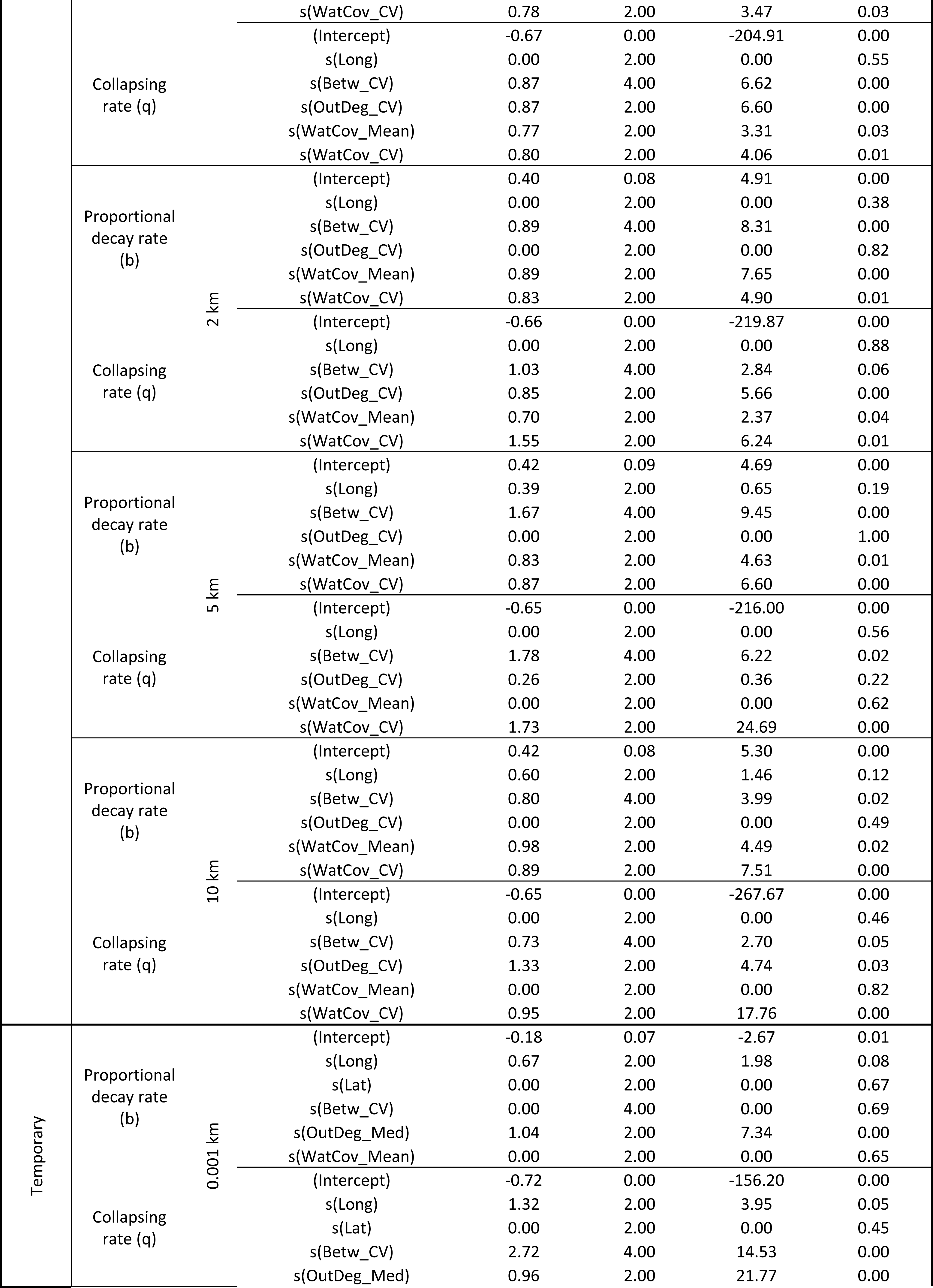
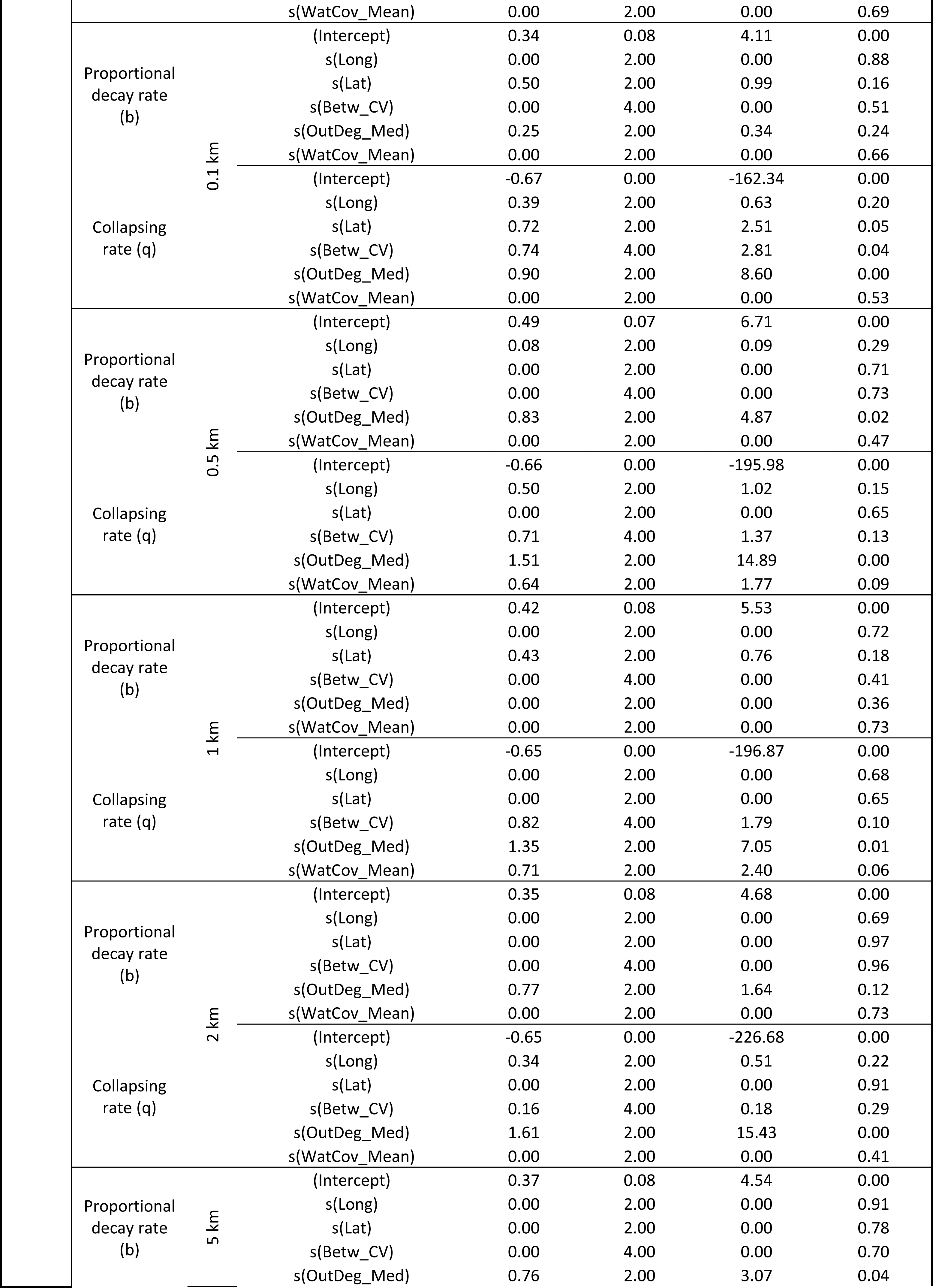
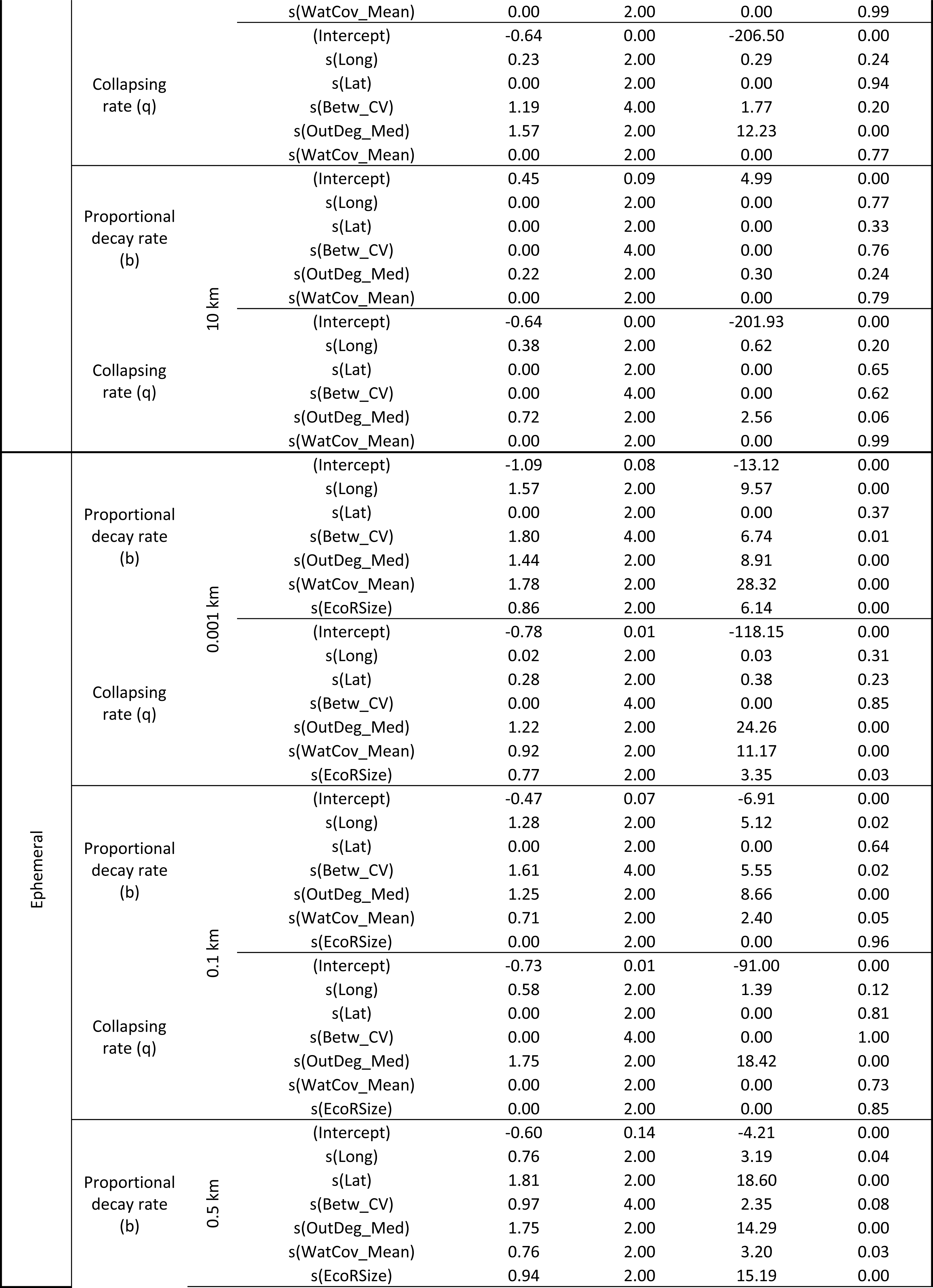
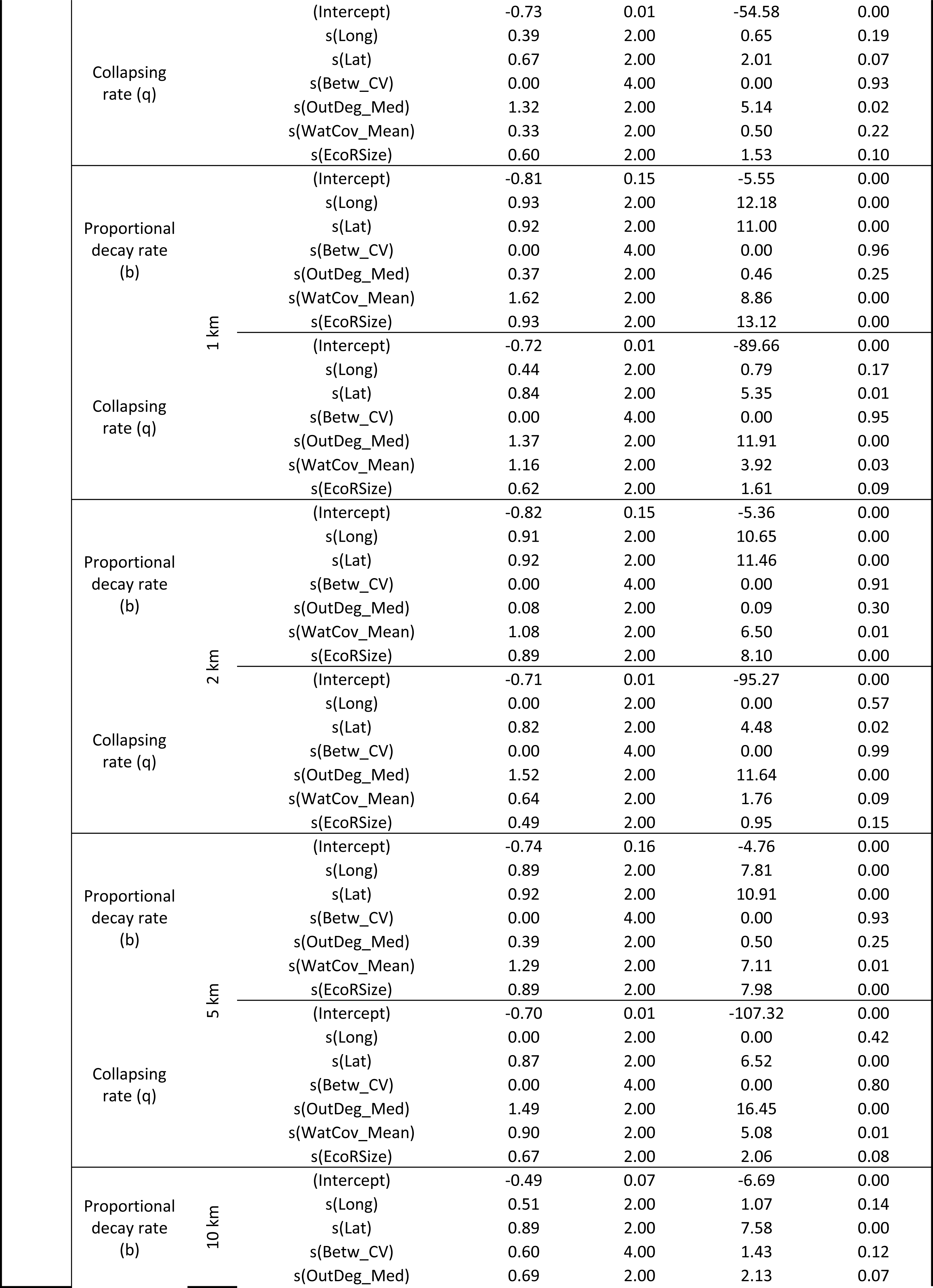
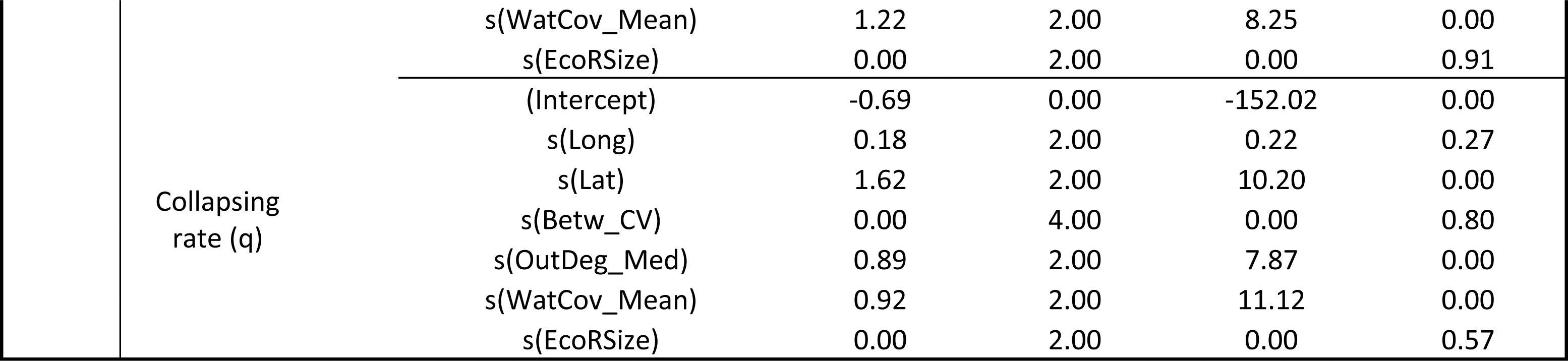
Gam model results for the intercept term (intercept) and the smoother term. Each smoother correspond to the selected descriptors after elimination of highly correlated ones. The complete list of descriptors is ecoregion mean longitude (Long), ecoregion mean latitude (Lat), ecoregion betweenness centrality median (Betw_Med), ecoregion betweenness centrality coefficient of variation (Betw_CV), ecoregion outdegree centrality median (OutDeg_Med), ecoregion outdegree centrality coefficient of variation (OutDeg_CV), mean ecoregion water cover (WatCov_Mean), ecoregion water cover coefficient of variation (WatCov_CV), ecoregion size (EcoRSize). Each model was carried for each type of freshwater (Total, Permanent, Temporary and Ephemera), Diversity decay parameter (Proportional decay rate and Collapsing rate) and the 7 dispersal distances (0.001 km, 0.1 km, 0.5 km, 1 km, 2 km, 5 km, 10 km). A summarizing barplot plot with the same information is also included to facilitate visual comparisons betewen models.

**Supplementary 9:**
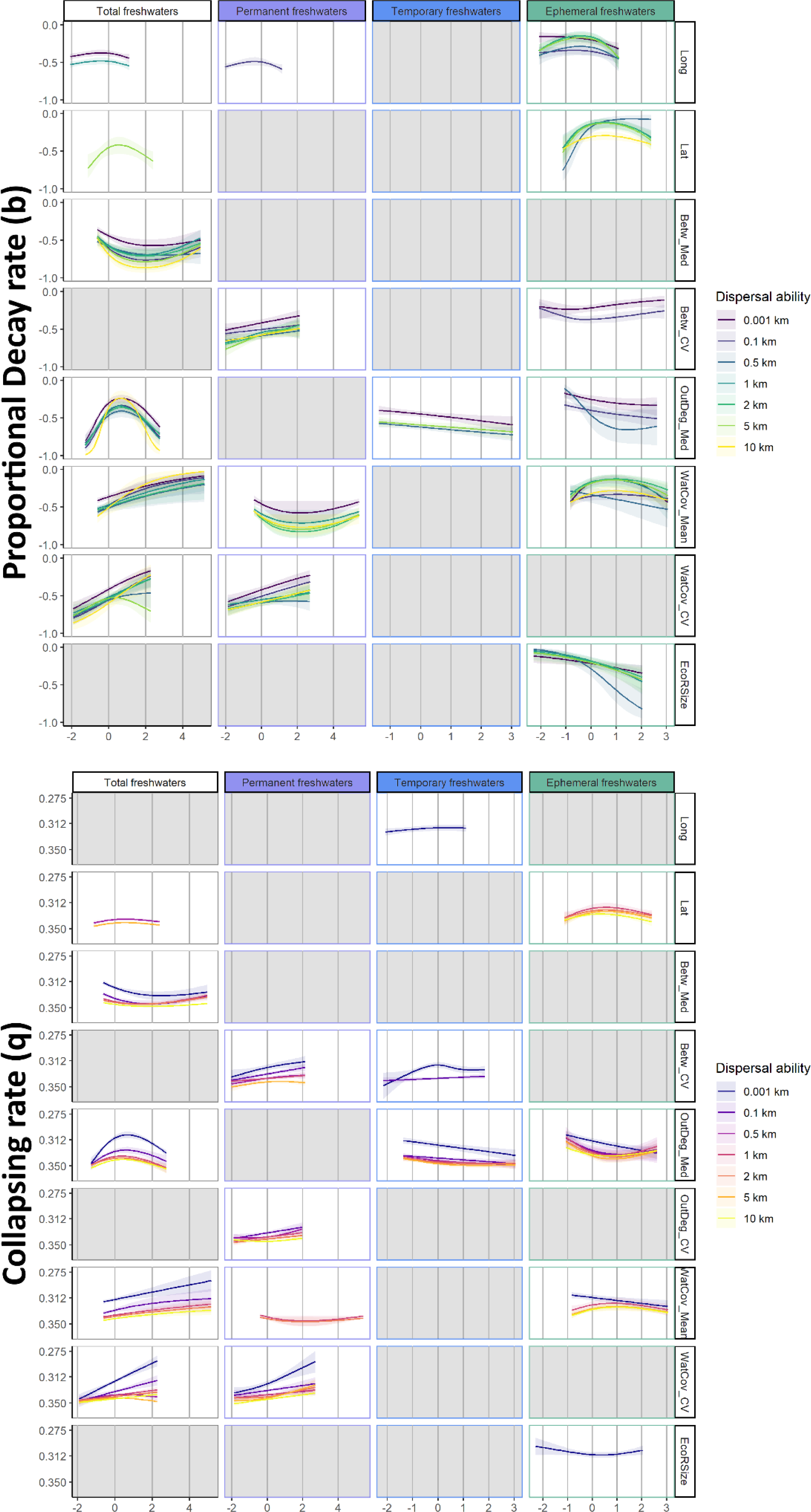
Landscape structural descriptors significant trends against the two diversity decay parameters Proportional decay rate (top plots) and Collapsing rates (bottom plots) and for each of the four freshwater types: Total freshwaters, Permanent freshwaters (purple panels), Temporary freshwaters (blue panels), and Ephemeral freshwaters (green panels). Dispersal abilities are identified with different colour gradients from low (0.001km) to high (10km) with purple-green-yellow gradient for proportional decay rate trends and blue-red-yellow for collapsing rates to ease visual identification. Only scaled landscape structural descriptors with significant trends are represented (x axes): ecoregion mean longitude (Long), ecoregion mean latitude (Lat), ecoregion betweenness centrality median (Betw_Med), ecoregion betweenness centrality coefficient of variation (Betw_CV), ecoregion outdegree centrality median (OutDeg_Med), mean ecoregion water cover (WatCov_Mean), ecoregion water cover coefficient of variation (WatCov_CV), ecoregion size (EcoRSize). Grey panels indicate non-significant relationships between structural driver and diversity decay parameters. Note that y axes are ordered differently ranging from 0 to ™1 for proportional decay rate (b) and from 0.2 to 0.4 for collapsing rate (q) to visually define high resistance to landscape degradation (decay rates close to 0 for both cases) as “high” y axis values and low resistance to landscape degradation as “low” y axis values.

## Notes

### Competing Interest Statement

The authors have declared no competing interest.

